# *Xist* Repeat A coordinates an assembly of SR proteins to recruit SPEN and induce gene silencing

**DOI:** 10.1101/2025.05.21.655143

**Authors:** Jackson B. Trotman, Alessandro Porrello, Scott A. Schactler, Lorenzo E. DeLeon, Quinn E. Eberhard, Samuel P. Boyson, Zhiyue Zhang, Shuang Li, David M. Lee, Sarah E. Kirik, Nadia Sultana, Son N. Nguyen, Michael-Claude G. Beltejar, Min Kyung Shinn, Shao-En Ong, Maria P. Gonzalez-Perez, Scott A. Shaffer, Daniel Dominguez, David M. Shechner, J. Mauro Calabrese

## Abstract

The lncRNA *Xist* represents a paradigm to understand the mechanisms of RNA-mediated gene silencing in mammals, which remain largely unresolved. To induce silencing, *Xist* recruits the RNA-binding protein SPEN through its 5′-proximal Repeat A domain. Yet, how Repeat A recruits SPEN and how SPEN coordinates silencing remain unclear. We report that sequences in Repeat A critical for SPEN recruitment directly bind SR-rich splicing factors. SRSF1, one such factor, is required for optimal SPEN recruitment and its RS-domain recruits SPEN when tethered to *Xist*. SPEN and SR-protein-binding motifs promote Repeat A’s association with many proteins, including the m^6^A machinery and elongating RNA polymerase II. SPEN also represses autosomal genes where its recruitment coincides with SR-protein binding. Our results reveal an unexpectedly essential role for splicing factors in coordinating silencing by *Xist* and suggest that the sensing of SR-protein-rich assemblies is a general mechanism through which SPEN targets genes for repression.

## INTRODUCTION

*Xist* is an 18-kilobase (kb) long noncoding RNA (lncRNA) that orchestrates X-chromosome inactivation, a regulatory process that silences transcription across an entire chromosome early during the development of female eutherian mammals. Once silencing by *Xist* is complete, it is propagated for life, making X-chromosome inactivation the most extensive and stable RNA-mediated silencing event identified. Nevertheless, the mechanisms of *Xist*-induced silencing, its evolutionary origins, and the extent to which analogous silencing events operate on autosomes remain incompletely resolved.^1,2^

*Xist* is distinguished by a unique sequence organization that contains several conserved domains embedded within exons of exceptional length.^3–7^ The most essential of these for gene silencing is Repeat A, a 450-nucleotide (nt) domain near the 5′ end of *Xist*. Repeat A is comprised of eight tandemly arrayed copies of a ∼50-nt monomer that itself is made up of a degenerate, structurally flexible U-rich spacer followed by a GC-rich “core” that is linearly conserved across eutherians.^8,9^ Repeat A’s gene silencing activity depends on its interaction with the RNA-binding protein SPEN.^10–13^

The mechanisms that recruit SPEN to *Xist* and coordinate SPEN-dependent silencing represent longstanding unknowns in our understanding of X-inactivation. Previous work has shown that in the context of a synthetic, consensus Repeat A sequence in an *Xist* cDNA transgene, mutations of the GC-rich cores – but not mutations or truncations of the U-rich spacers – impair silencing as measured indirectly by a cell-survival assay.^14,15^ Perplexingly, however, the same GC-rich-core mutations had no qualitative impact on SPEN binding *in vitro*,^12^ and SPEN directly binds Repeat A through its U-rich spacers.^16–18^ Chemical probing suggests that sequence motifs across Repeat A exist in single-stranded states that bind proteins (Figure S1A),^7,9^ hinting that additional Repeat-A-binding proteins may mediate SPEN recruitment.^19^

Once recruited to Repeat A, SPEN has been proposed to coordinate interactions between histone deacetylates and other histone-modifying enzymes to mediate silencing by *Xist*.^11,20^ However, while deletion of SPEN or Repeat A results in a near-complete loss of *Xist*-induced silencing, inhibition or deletion of SPEN-associated histone deacetylates, including HDAC3, have only minor effects.^20–22^ Thus, Repeat A and SPEN employ yet-to-be-defined mechanisms to induce silencing.^19^ Likewise, the manner and extent to which SPEN regulates gene expression in contexts beyond X-inactivation remain unknown.

Curiously, in addition to its role in silencing, Repeat A harbors other functions. For reasons that remain unclear, Repeat A deletion reduces *Xist* abundance in a manner that depends partly on SPEN.^23–30^ Relatedly, in a transgenic context, Repeat A promotes readthrough of proximal polyadenylation signals, an activity that depends on its GC-rich cores and not SPEN.^27^ Lastly, Repeat A associates with serine-and-arginine-rich (SR) proteins *in vivo* and *in vitro*, and its deletion alters *Xist* splicing patterns.^3,6,27–29,31^ Whether these additional functions are relevant to silencing by Repeat A remains unclear.

The many unknowns highlight a need to study Repeat A in quantitative contexts that report on all aspects of *Xist* biology. Such studies would provide insights into the mechanisms of RNA-mediated gene silencing as well as into the functions of SPEN, a deeply conserved protein recurrently mutated in cancers and neurodevelopmental disorders whose molecular functions remain poorly characterized.^19,32–42^ Below, we demonstrate that SR-protein-binding motifs in Repeat A’s GC-rich cores and at least one SR protein, SRSF1, are critical for recruitment of SPEN and *Xist*-induced gene silencing. The N-terminal region of SPEN is an SR-protein sensor that facilitates its recruitment to both Repeat A and SR-protein-binding motifs transcriptome-wide. SPEN and SR-protein-binding sequences synergistically enable *Xist* to associate with the m^6^A methylation, RNA turnover, and transcriptional elongation machineries. Lastly, SPEN represses autosomal genes that it engages through SR-protein-bound RNA. Our findings provide new insights into the mechanisms and evolutionary origins of *Xist*-induced silencing and suggest that the sensing of SR-protein-rich assemblies is a general mechanism by which SPEN targets genes for repression.

## RESULTS

### Sequences critical for *Xist*-induced silencing, *Xist* production, and *Xist* splicing fidelity are located within the GC-rich cores of Repeat A

To clarify how sequence elements throughout Repeat A impact *Xist* biogenesis and function, we sought to evaluate the effects of new and previously investigated^12,14,15^ mutations in the context of a full-length doxycycline (dox)-inducible *Xist* transgene that includes its exons and introns. *Xist* transgenes were inserted as a single copy into the C57BL/6 (B6) allele of the *Rosa26* locus of an F1-hybrid mouse embryonic stem cell (ESC) line.^27^ In these ESCs, *Xist* expression represses genes on the B6 copy of chromosome 6 (chr6) without affecting expression on the CAST/EiJ (CAST) chromosome (Figure 1A).^27,43^

**Figure 1.**
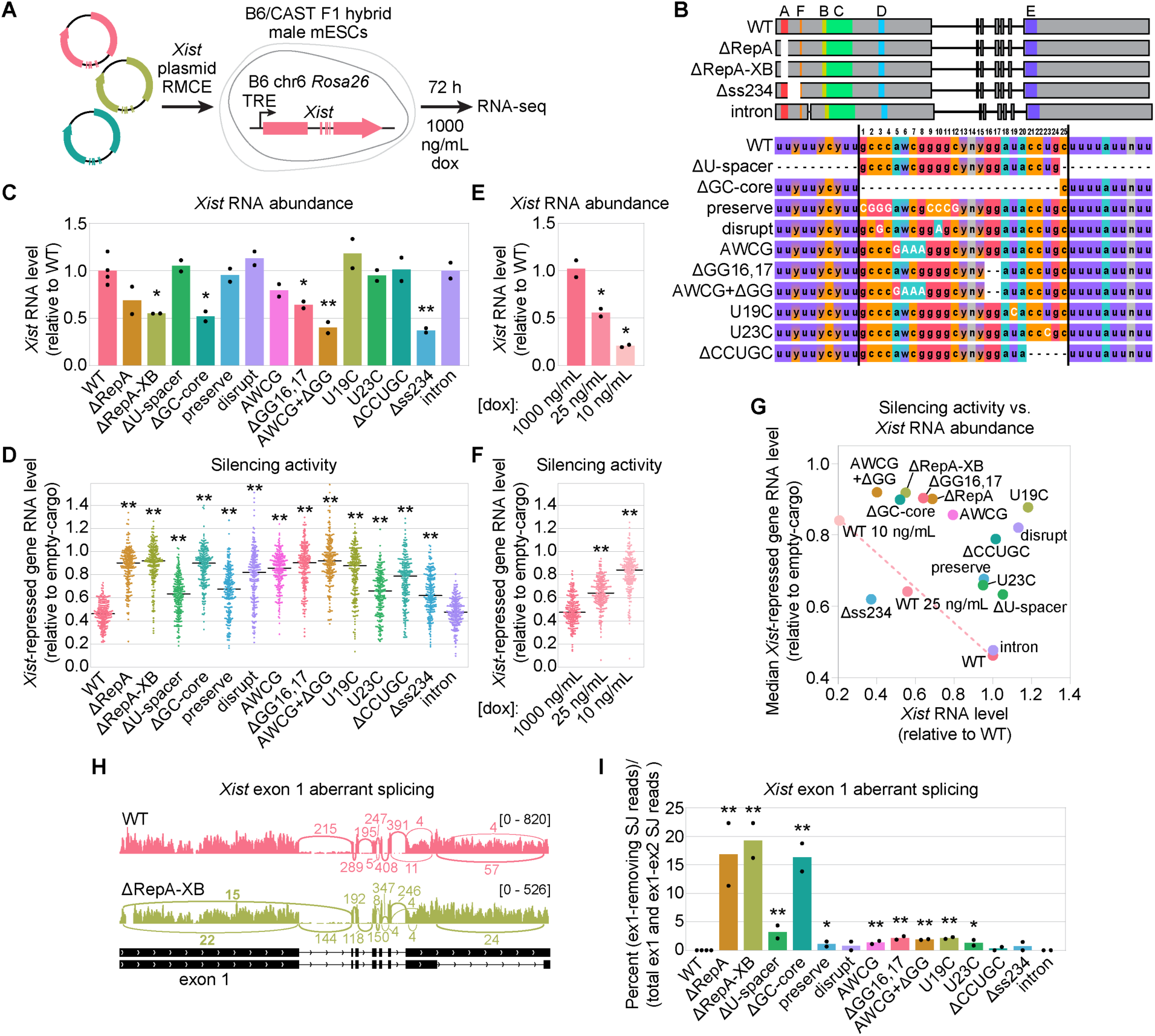
Sequences critical for *Xist*-induced silencing, *Xist* production, and *Xist* splicing fidelity are located within the GC-rich cores of Repeat A. **(A)** *Xist* gene insertion into *Rosa26*.^43^ **(B)** Summary of *Xist* mutants, including mutation positions in each Repeat A monomer. (C, E) *Xist* abundance relative to WT by RNA-seq across mutants (C) or WT dox titration (E). Dots, independent clonal cell lines. (D, F) *Xist* silencing activity by RNA-seq for *Xist* mutants (D) or WT dox titration (F). Dots, sample-averaged expression of B6 chr6 *Xist*-repressed genes relative to empty-cargo control. Black bars, median. **(G)** Median silencing vs. *Xist* RNA abundance. Dashed line, silencing expected for corresponding levels of WT *Xist*. **(H)** Sashimi plots illustrating normal (WT) and aberrant (ΔRepA-XB) *Xist* splicing. **(I)** Quantification of aberrant splicing across genotypes (dots, independent clonal cell lines). SJ, splice junction. For all panels vs. WT (two-sided t-test): (*), *p* < 0.05; (**), *p* < 0.01. See Figures S1, S2.

Using this system, we made two separate excisions of Repeat A and ten smaller mutations or deletions within each monomer of Repeat A (Figure 1B; Figure S2). Additionally, we made a deletion as well as an insertion downstream of Repeat A (Figure 1B). The deletion encompassed a 742-nt region immediately downstream of Repeat A that is predicted to form at least three stable RNA secondary structures (“ss234”)^9,44^ and harbors the highest N6-methyladenosine (m^6^A) density within *Xist*.^26^ This region also contains the only transcribed CpG island in *Xist*, perhaps related to its requirement for readthrough of proximal polyadenylation signals following a 2-kb *Xist* transgene.^27,45,46^ Separately, we inserted a synthetic intron^47^ just downstream of Repeat F. Previous work suggested that ectopic splicing of an intron downstream of Repeat A may attenuate *Xist*-induced silencing,^48^ and we sought to test that hypothesis. Four independent wild-type (WT) *Xist* clonal lines and two independent clonal lines for each *Xist* mutant were selected and rendered dox-inducible via piggyBac-mediated insertion of the reverse tetracycline-controlled transactivator (rtTA).^27^ In parallel, we prepared four independent clonal lines of empty-cargo control ESCs containing the same dox-inducible promoter but no lncRNA sequence inserted.

We first defined a high-confidence set of *Xist*-repressed genes on chr6. We performed RNA-seq for each wild-type *Xist* and empty-cargo cell line treated with dox for 3 days (Figure 1A). Comparing allelic expression between the two genotypes, we identified 204 chr6 genes significantly repressed by *Xist* (adjusted *p*-value < 0.001).^49^ As expected, *Xist*-repressed genes were located along the length of chr6, and were, on average, repressed by about 50% relative to empty-cargo control (Figure S1C). Replicates across genotypes were highly reproducible (Figure S1D,E).

We next performed RNA-seq for our panel of mutants to assess effects on *Xist*-induced gene silencing, *Xist* production, and *Xist* splicing. This revealed a broad range of phenotypes, many of which were unanticipated. As expected, Repeat A deletion led to a near-complete loss of silencing and a ∼50% reduction in *Xist* abundance (Figure 1C,D, Figure S1F). Repeat A deletion also caused previously reported aberrant splicing events that excise the bulk of *Xist* exon 1; an estimated ∼15-20% of *Xist* transcripts in Repeat A deletion mutants lacked a functional exon 1 (Figure 1H,I).^29,50^ Surprisingly, deletion of the U-rich spacers – Repeat A’s presumed SPEN-binding region – only intermediately affected silencing, did not alter *Xist* abundance, and minimally altered *Xist* splicing, whereas deletion of the GC-rich cores phenocopied deletion of Repeat A in all respects (Figure 1C,D,I). Mutations predicted to preserve or disrupt possible secondary structure within the GC-rich cores^14^ did not impact *Xist* abundance and minimally impacted splicing but did have moderate (“preserve” mutant) to severe (“disrupt” mutant) effects on silencing (Figure 1C,D,I). Strikingly, mutation of a predicted single-stranded AWCG motif previously shown to bind splicing factor SRSF1,^28^ deletion of a GG dinucleotide downstream of the AWCG, or a combination of both mutations in the same *Xist* gene phenocopied Repeat A deletion in terms of reduced *Xist* abundance and silencing. These mutations also altered exon 1 splicing but to a much lesser degree than Repeat A or GC-rich core deletion (Figure 1C,D,I). Mutation of U19 or deletion of the CCUGC motif had severe defects on silencing without reducing *Xist* abundance; U19 mutation altered splicing to a similar degree as the AWCG mutation, whereas CCUGC deletion did not (Figure 1C,D,I). Mutation of U23 weakly reduced silencing, weakly affected splicing, and did not alter *Xist* abundance (Figure 1C,D,I). Deletion of ss234 affected both *Xist* abundance and silencing and had no effect on splicing (Figure 1C,D,I). Insertion of the ectopic intron phenocopied wild-type *Xist* in all respects (Figure 1C,D,I); *Xist*-induced silencing was unaffected by variably efficient ectopic intron splicing (Figure S1D,G).

Leveraging our panel of mutants, we sought to determine the extent to which reductions in silencing activity could be attributed to concomitant reductions in *Xist* abundance. We performed RNA-seq in ESCs expressing lower levels of wild-type *Xist* and demonstrated a linear relationship between silencing and *Xist* abundance (Figure 1E-G, Figure S1E), consistent with expectations set by prior works.^43,51,52^ Next, plotting silencing activity as a function of *Xist* RNA levels for all genotypes and expression conditions revealed that every mutant except two (Δss234 and the intron-insertion) affected silencing to a greater degree than could be attributed solely to reduced *Xist* abundance (Figure 1G). These experiments confirm a role for Repeat A in silencing beyond its role in promoting *Xist* production.

Taken together, we observed a striking non-uniformity of phenotypes upon deletion or mutation of Repeat A, and the sequence elements most critical for *Xist*-induced silencing, *Xist* production, and suppression of aberrant splicing are located in the GC-rich cores of Repeat A and not its U-rich spacers.

### SPEN association with Repeat A *in vivo* depends on sequences that it does not bind *in vitro*

SPEN is recruited by Repeat A and is required for *Xist*-induced silencing.^10–12,20^ However, *in vitro*, SPEN directly binds the U-rich spacers of Repeat A and not the GC-rich cores critical for silencing.^12,16,17^ To address this apparent discrepancy, we performed RNA immunoprecipitation from formaldehyde-crosslinked cells, followed by quantitative PCR (RIP-qPCR),^43^ to measure the extent to which SPEN associates with each *Xist* mutant *in vivo* (Figure 2A). Because formaldehyde induces RNA-protein and protein-protein crosslinks, RIP enables detection of both direct and indirect (i.e., protein-bridged) RNA-protein interactions (Figure 2A). We confirmed that our SPEN antibody retrieved Repeat A in a SPEN-dependent manner by performing RIP-qPCR with wild-type and *Spen*-knockout ESCs (Figure 2B; Figure S3A,B).^53^ In parallel, we examined SPEN association across our panel of mutants, normalizing signal by the amount of *Xist* RNA in each corresponding input (Figure 2B). Consistent with the necessity of SPEN for *Xist*-induced silencing, we observed a remarkably strong correlation between SPEN RIP recovery and extent of *Xist*-induced gene silencing across all mutants (Figure 2C; Pearson’s *r* = −0.91; *p* = 5 x 10^-6^). The mutants associating with the least amount of SPEN *in vivo* were ΔRepA; ΔGC-core; ΔGG16,17; and the AWCG+ΔGG mutant (Figure 2B). Recovery of a non-specific control region was uniform across genotypes (Figure 2B). Therefore, the sequences most essential for SPEN-Repeat A association *in vivo* are not U-rich but are instead contained within its GC-rich cores.

**Figure 2.**
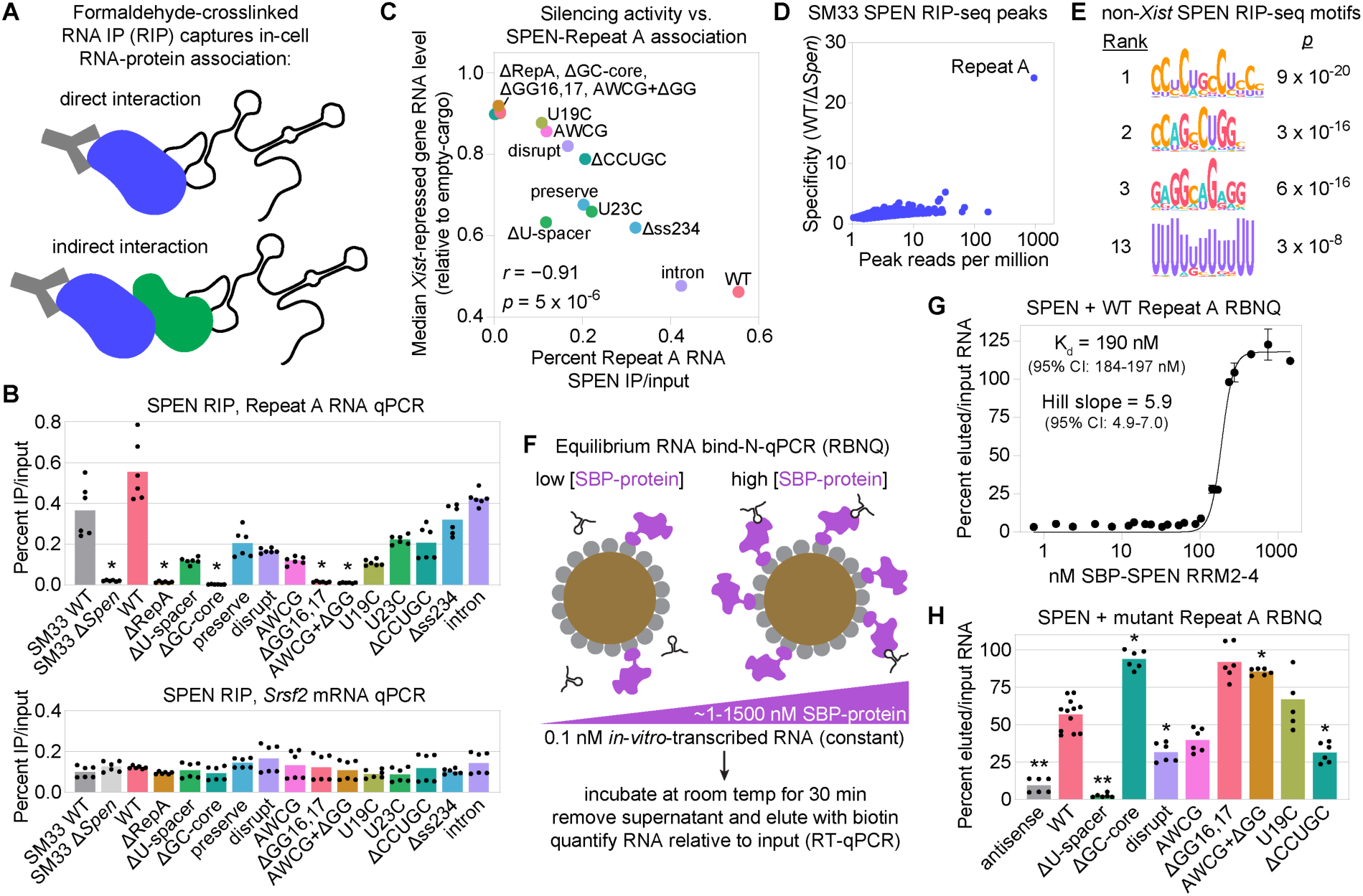
SPEN association with Repeat A *in vivo* depends on sequences that it does not bind *in vitro*. **(A)** RIP schematic.^43^ **(B)** SPEN-Repeat A association across genotypes by input-normalized RIP-qPCR. SM33 WT vs. Δ*Spen* ESCs confirm specificity of SPEN antibody.^53^ *Srsf2* mRNA, control. Dots, triplicate qPCR values from two independent experiments. **(C)** Median silencing vs. SPEN-Repeat A association across genotypes. *r*, *p*: data fit to linear regression. **(D)** Specificity vs. signal for SPEN RIP-seq peaks in SM33 ESCs. **(E)** Motifs from top 1500 non-*Xist*, non-*Spen* SPEN RIP-seq peaks. **(F)** Overview of equilibrium RBNQ. SBP, streptavidin-binding peptide. **(G)** RBNQ with SPEN RRM2-4 and Repeat A. Dots, values from two independent experiments. Error bars, standard deviation of qPCR triplicates. CI, confidence interval. **(H)** RBNQ across Repeat A RNAs using fixed SPEN RRM2-4 concentration. Dots, qPCR triplicates from ≥2 independent experiments. For all panels vs. WT (two-sided t-test): (*), *p* < 0.05; (**), *p* < 0.01. See Figure S3.

To determine whether GC-rich sequences can direct SPEN association with RNAs other than *Xist*, we sequenced RNA recovered by SPEN RIPs from wild-type and *Spen*-knockout SM33 ESCs (Figure 2B). We identified 69,045 genomic regions (“peaks”) enriched in SPEN association. This revealed that Repeat A is by far the most-recovered RNA target of SPEN and also the most specifically recovered from wild-type versus *Spen*-knockout cells (Figure 2D). Nonetheless, we identified thousands of RNA peaks whose wild-type recovery was at least two times greater than knockout across two independent experiments. Analysis of peaks with STREME^54^ revealed enrichments for multiple CUG- and GA-rich motifs (Figure 2E, Figure S3C). A single poly-U motif, ranked 13^th^, was also enriched. These results suggest that, while SPEN can associate with U-rich sequences *in vivo*, it more frequently associates with G- and C-rich sequences, consistent with our *Xist*-mutant RIP results.

Next, we examined how the different mutations impact the relative binding affinity between Repeat A RNA and SPEN *in vitro*. SPEN directly binds RNA through its RNA-recognition motifs 2 through 4 (RRM2-4), located near its N-terminus.^18^ We therefore used a modified version of the RNA-bind-n-qPCR assay (RBNQ)^55^ to measure the equilibrium binding affinity between bacterially purified SPEN RRM2-4 and *in vitro* transcribed Repeat A RNA (Figure 2F; Figure S3D,E). SPEN RRM2-4 bound to wild-type Repeat A RNA cooperatively and with a K_d_ of 190 nM, consistent with prior studies (Figure 2G).^17,18,56^ We next performed RBNQ across our panel of mutant RNAs using a concentration of SPEN RRM2-4 that yielded ∼50% recovery of wild-type RNA, enabling detection of both increases and decreases in binding (Figure 2F). Relative to wild-type Repeat A, U-rich-spacer deletion abrogated binding, consistent with expectations from prior *in vitro* experiments (Figure 2H;^12,16^). However, GC-rich-core deletion increased binding, and mutations within the cores either increased binding, made no difference, or reduced binding slightly but not to the same degree as deletion of the U-rich spacers (Figure 2H). Together, these findings confirm that the sequences within Repeat A that directly bind SPEN *in vitro* (its U-rich spacers) are not the ones most critical for SPEN association *in vivo* (its GC-rich cores).

### Motif-dependent association between Repeat A and SR proteins *in vitro* and *in vivo*

To rationalize the sharp discordance between SPEN’s preferred motifs *in vitro* and *in vivo*, we hypothesized that recruitment of SPEN to Repeat A *in vivo* might require proteins that interact with the GC-rich cores. To identify candidate interacting proteins, we modified a previous approach^3^ in which internally biotinylated wild-type or mutant Repeat A RNA was incubated with ESC nuclear extracts, crosslinked with UV light, and streptavidin-purified prior to being analyzed by mass-spectrometry-based proteomics (Figure 3A, Figure S4A,B). We also used proteomics to quantify the proteins detectable in our nuclear extracts. In total, we identified 644 proteins reproducibly recovered with Repeat A RNA but not no-RNA control (Figure S4B; Figure S4C; Table S1). SPEN was not recovered, presumably due to its low abundance in ESCs (2766^th^ out of 2826 detected nuclear extract proteins; Table S1).

**Figure 3.**
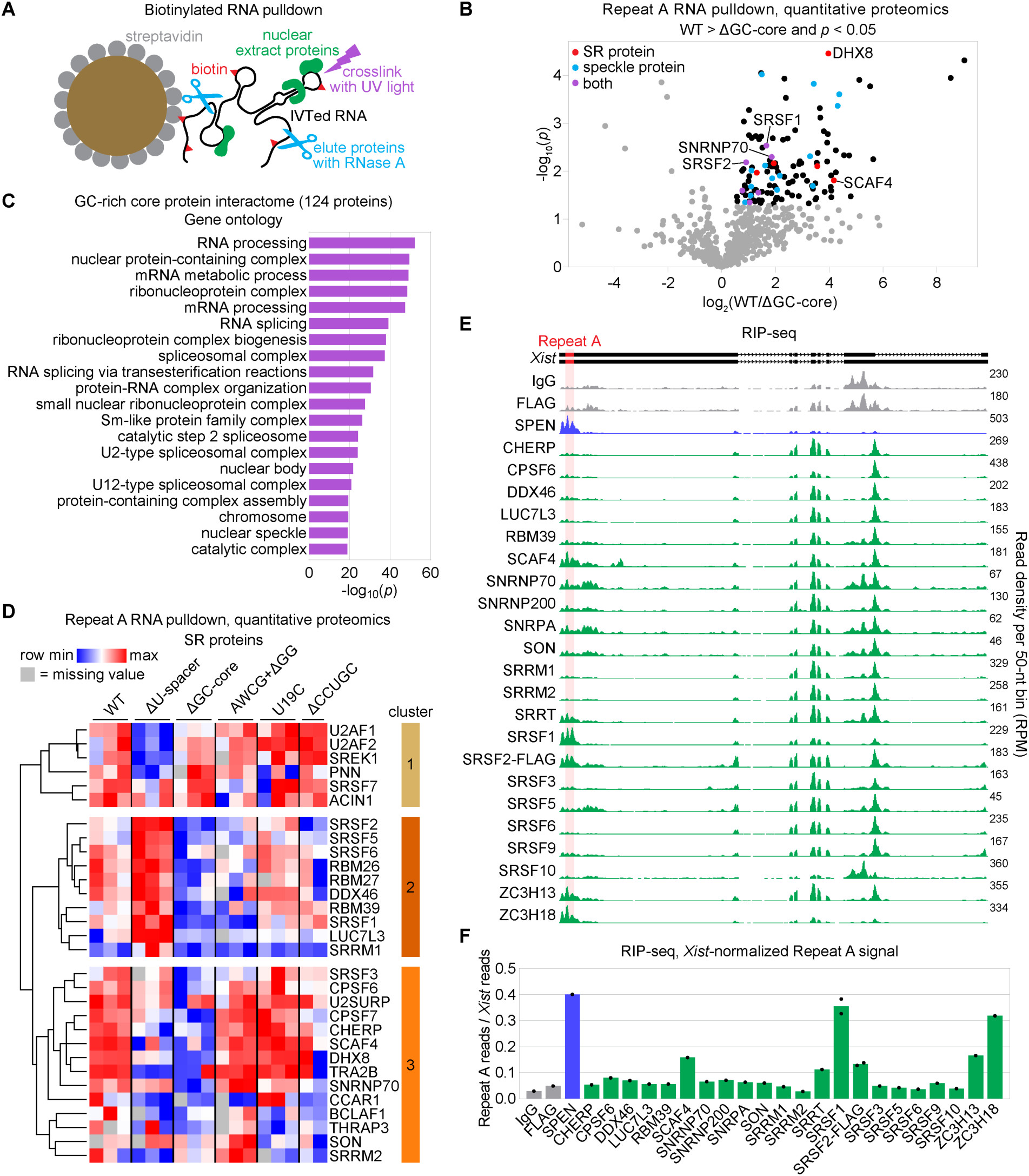
Motif-dependent association between Repeat A and SR proteins *in vitro* and *in vivo*. **(A)** Strategy to identify Repeat A-interacting proteins *in vitro*. **(B)** Proteins enriched by WT over ΔGC-core Repeat A (*p* < 0.05, two-sided t-test, n=3). **(C)** GO terms of GC-core-specific proteins (top 20). **(D)** Relative recovery of SR proteins across Repeat A RNAs. Columns, independent experiments. **(E)** RMCE-*Xist* WT RIP-seq with antibodies against controls, SPEN, and SR and/or RNA-processing proteins. **(F)** RIP-seq signal within Repeat A relative to total *Xist*. Dots, independent experiments. See Figure S4.

Of the 644 Repeat A-recovered proteins, 124 were significantly enriched with wild-type RNA relative to ΔGC-core RNA (*p* < 0.05, two-sided t-test; Figure 3B). Gene ontology analysis of the GC-core-interacting proteins revealed an enrichment in RNA processing and splicing factors, including components of the spliceosome and nuclear speckles (Figure 3C). We noted distinct motif-dependent enrichment patterns of SR proteins, a protein family distinguished by disordered serine- and arginine-rich domains that plays central roles in transcription and RNA processing (Figure 3D).^57,58^ Indeed, relative to wild-type Repeat A, several SR proteins displayed reduced recovery upon deletion of the GC-rich cores and enhanced recovery by RNA containing only GC-rich cores (the ΔU-spacer mutant), strongly suggesting interactions with the GC-rich cores (Figure 3D, cluster 2).

We next sought to determine whether the proteins identified to interact with Repeat A *in vitro* associate with Repeat A *in vivo*. Using RIP-seq, we analyzed a panel of 22 RNA processing factors, including components of the spliceosome and nuclear speckle, with a focus on SR proteins. We analyzed SPEN as a positive control, and nonspecific IgG and anti-FLAG antibodies as negative controls. Repeat A association was enriched at least two-fold over negative controls for five SR proteins: SRSF1, SRSF2, ZC3H13, ZC3H18, and SCAF4 (Figure 3E,F; Figure S4D). RIPs for other proteins more closely resembled negative controls, indicating poor antibody performance or non-specific association with *Xist*. We conclude that *in vivo*, Repeat A associates with SR proteins that *in vitro* interact with Repeat A in a GC core-dependent manner.

### SPEN associates with nuclear speckle components and other SR proteins via its N-terminus

Next, we investigated whether SPEN interacts with SR proteins *in vivo*. We first deleted *Spen* from RMCE-*Xist* WT ESCs and stably re-expressed SPEN from a cDNA harboring an N-terminal Halo tag and a C-terminal V5 tag, enabling immunoprecipitation and western blot detection (Figure S5A,B). Consistent with previous observations,^26^ *Spen* deletion caused a 50% decrease in *Xist* levels (Figure 4A) and a loss of *Xist* silencing activity beyond that expected for comparable lower levels of wild-type *Xist* (Figure 1G, Figure S5C). While ∼18% of *Xist* transcripts displayed aberrant splicing-out of *Xist* exon 1 upon Repeat A deletion (Figure 1H,I), only 4% did so upon *Spen* deletion, indicating that SPEN plays a minor role in preserving *Xist* splicing fidelity (Figure S5D). Expressing Halo-SPEN-V5 in *Spen* knockout cells restored *Xist* abundance and splicing fidelity to wild-type levels and rescued silencing activity, albeit only partially, possibly due to the production of SPEN at lower-than-endogenous levels (Figure 4A; Figure S5C,D).

**Figure 4.**
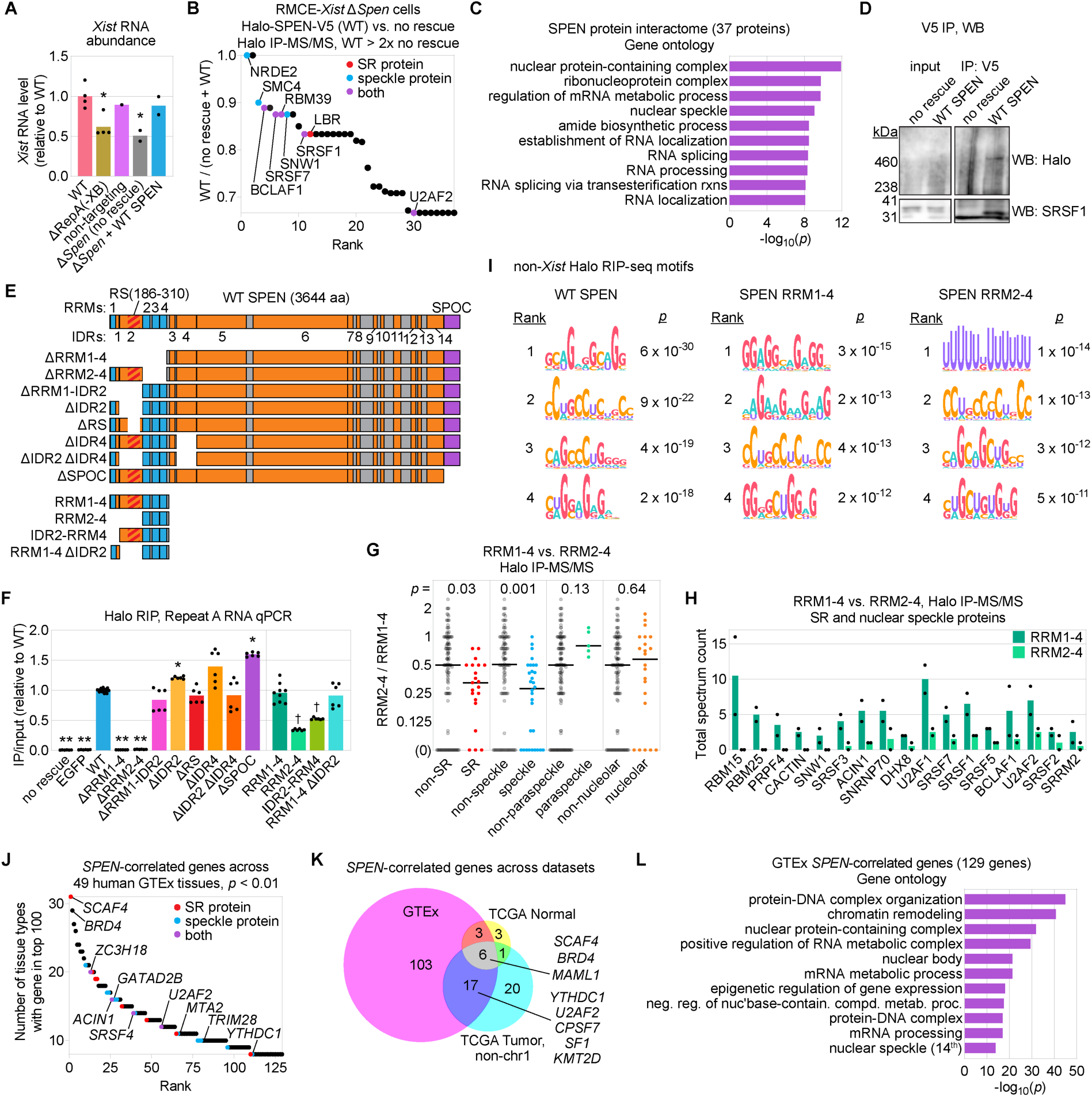
SPEN associates with nuclear speckle components and other SR proteins via its N-terminus. **(A)** *Xist* abundance across genotypes (RNA-seq). Dots, independent clonal lines. **(B)** Halo-SPEN-V5-interacting proteins ranked by proportion of peptides recovered. **(C)** GO terms of SPEN-interacting proteins (top 10). **(D)** V5 IP, Halo and SRSF1 western blots (no-rescue vs. Halo-SPEN-V5 ESCs). **(E)** SPEN domains and deletion mutants. IDRs, MobiDB.^131^ **(F)** SPEN-Repeat A association for WT SPEN vs. mutants by input-normalized Halo RIP-qPCR. Dots, qPCR triplicates (≥2 independent experiments). Asterisks, significantly different from WT. Crosses, significantly different from RRM1-4. **(G)** Enrichment of proteins, by class, in RRM1-4 vs. RRM2-4 IP-MS/MS. “(0),” no recovery with RRM2-4. **(H)** Peptide counts of SR and/or speckle proteins in RRM1-4 or RRM2-4 IP-MS/MS. Dots, independent experiments. **(I)** Motifs from top 1500 non-*Xist*, non-*Spen* Halo RIP-seq peaks across genotypes. **(J)** Significant *SPEN*-correlated genes from GTEx (guilt-by-association; *p* < 0.01 and FDR < 0.1, see Methods). **(K)** Overlap of significant *SPEN*-correlated genes (FDR < 0.1) from the GTEx, TCGA Tumor (non-chr1), and TCGA Normal datasets. **(L)** GO terms of *SPEN*-correlated genes from GTEx (top 10 and 14^th^). Excepting (J), *p*-values, two-sided t-tests: (*/†), *p* < 0.05; (**), *p* < 0.01. See Figure S5.

To identify SPEN-interacting partners, we immunoprecipitated Halo-SPEN-V5 from formaldehyde-crosslinked cells with RIP wash conditions and analyzed recovered proteins via mass spectrometry. Parallel immunoprecipitations from *Spen* knockout cells expressing no Halo-tagged construct (“no rescue”) served as a specificity control. We identified 37 proteins whose signal was enriched at least two-fold over no-rescue control in both replicates, while also observing that SPEN peptides were only detected from its N-terminal half (Figure 4B, Table S2; not shown). Gene ontology analysis revealed significant enrichments with many of the same terms as our Repeat A affinity proteomics, including those related to splicing, RNA processing, and nuclear speckles (Figure 3C, Figure 4C). The enrichment of nuclear speckle components and SR proteins was especially striking, with 8 of the top 19 SPEN-interacting proteins belonging to one or both categories (Figure 4B). One such protein, SRSF1, is an SR and nuclear speckle protein previously found to interact with SPEN,^57–60^ which we confirmed via co-immunoprecipitation (Figure 4D).

We next examined how individual parts of SPEN contribute to its interaction with Repeat A and other proteins. SPEN is a ∼400-kilodalton protein with 4 N-terminal RRMs and a C-terminal SPOC domain thought to mediate interactions with protein effectors (Figure 4E).^20^ The rest of SPEN primarily consists of 14 intrinsically disordered regions (IDRs) defined by MobiDB (Figure 4E).^61^ Reminiscent of canonical SR proteins, SPEN contains a bona fide RS domain within its RRM-adjacent IDR2.^57,62^ IDR4, downstream of RRM4, is also arginine-rich, with many RE/ER dipeptides that can mimic phosphorylated RS dipeptides (Figure S5E,F).^63^ These regions were of particular interest due to the roles that RS domains play in mediating interactions among SR proteins.^57,58^ We constructed a panel of 12 Halo-and-V5-tagged SPEN deletion mutants and analyzed association with Repeat A via RIP-qPCR (Figure 4E). As expected, wild-type SPEN recovered Repeat A robustly relative to no-rescue and EGFP-Halo controls (Figure 4F). Deletion of RRM1-RRM4 completely abrogated SPEN’s association with Repeat A, as did deletion of RRM2-4, consistent with previous results.^20,52,64^ In contrast, deletion of the RRM1-IDR2 region had no impact on Repeat A association, nor did deletion of IDR2, the smaller RS domain, IDR4, or a combined deletion of IDR2 and IDR4. Intriguingly, deletion of the SPOC domain caused a ∼50% increase in SPEN-Repeat A association. Lastly, we found that RRM2-4 alone was sufficient to associate with Repeat A but at a level ∼3-fold lower than the larger RRM1-4 construct. This difference was mostly attributable to RRM1, although the addition of IDR2 to RRM2-4 also caused a minor increase in Repeat A association (Figure 4F). These results confirm that RRMs 2-4 are essential for Repeat A recognition and demonstrate that SPEN’s N-terminus (RRM1-IDR2) promotes Repeat A association in a truncated version of SPEN and likely in combination with downstream IDRs in the context of full-length SPEN.

Previous work demonstrated no difference in direct Repeat A binding affinity between RRM1-4 and RRM2-4, implying that SPEN’s N-terminus functions outside of RNA-binding.^18^ To investigate this function, we used Halo immunoprecipitation and proteomics to compare protein association between the RRM1-4 and RRM2-4 constructs. Likely due to higher expression of these constructs than full-length SPEN (Figure S5B), we identified 146 and 106 proteins enriched with RRM1-4 and RRM2-4, respectively, at least two-fold over no-rescue control in two independent replicates (Table S2). GO analysis of the seventy-five proteins enriched with RRM1-4 at least two-fold over RRM2-4 revealed strong enrichments for terms related to splicing and nuclear speckles (Figure S5G). Indeed, compared to RRM2-4, RRM1-4 recovered significantly higher levels of nuclear speckle and SR proteins – including known Repeat A interactors SRSF1, SRSF2, U2AF1, U2AF2, and RBM15 – but not paraspeckle or nucleolar proteins (Figure 4G,H).^65^ In parallel, we performed RIP-seq to determine motifs enriched in the top non-*Xist* peaks for Halo-tagged full-length SPEN, RRM1-4, and RRM2-4 transcriptome-wide. Like endogenous SPEN (Figure 2E), full-length Halo-SPEN-V5 associated most strongly with GA- and GC-rich, SR-protein-like binding motifs (Figure 4I). Notably, while RRM1-4 similarly enriched for SR-protein-like binding motifs, RRM2-4 enriched most strongly for poly-U motifs, consistent with the latter’s direct RNA-binding preference (Figure 2H, Figure 4I). Therefore, relative to RRM2-4, RRM1-4 associates more strongly with SR proteins, SR-protein-binding motifs, and Repeat A itself.

Lastly, we investigated the proteins genetically associated with SPEN via an orthogonal method, performing “guilt-by-association” analysis with data from the Genotype-Tissue Expression (GTEx) project.^66–69^ We identified 129 genes whose expression was significantly correlated with *SPEN* across 49 GTEx tissues (FDR < 0.1, see Methods), implying that they may function within the same or related pathways (Figure 4J, Table S3). Additional guilt-by-association analyses of normal and tumor tissue data from The Cancer Genome Atlas (TCGA) identified many of the same genes and pathways (Figure 4K, Figure S5H, Table S3), including *MAML1* and *KMT2D*, genes previously linked to SPEN in the context of NOTCH signaling.^70,71^ Genes encoding SPEN-interacting proteins (merged list of WT, RRM1-4, and SPOC domain interactors; Table S2)^20^ were significantly enriched in *SPEN*-correlated genes (*p* = 0.0004, two-tailed binomial test), further underscoring our ability to identify meaningful connections to SPEN via guilt-by-association. Consistent with our findings above, we observed strong enrichment for GO terms associated with splicing and nuclear speckles, along with terms and proteins related to transcriptional activation and repression (Figure 4J-L, Figure S5H). These include the top hit, SCAF4, an RS-domain-containing transcriptional antiterminator enriched over Repeat A (Figure 3E);^72^ BRD4, a critical transcriptional elongation factor;^73^ TRIM28 and NuRD (GATAD2B, MTA2), epigenetic modifiers previously linked to SPEN- and Repeat A-dependent silencing;^20,74^ ZC3H18, an SR protein and RNA exosome adapter enriched over Repeat A (Figure 3E);^75,76^ the SR proteins ACIN1, SRSF4, and U2AF2; and YTHDC1, a (D/E)R-rich, nuclear speckle-associated m^6^A-binding protein and exosome adapter also enriched over Repeat A (Figure 3E,F, Figure 4H, Figure S7).^77–80^ Thus, proteomic, genomic, and guilt-by-association analyses connect SPEN with multiple aspects of the transcriptional process and nominate the N-terminal region of SPEN as an SR protein sensor.

### SRSF1 directly binds a conserved Repeat A motif to recruit SPEN to *Xist in vivo*

To determine whether SR-rich splicing factors play a role in recruiting SPEN to Repeat A or other RNAs, and for the following reasons, we focused on SRSF1: (i) SRSF1 interacts with Repeat A *in vitro* and *in vivo*, and, after SPEN, was the most Repeat-A-enriched protein we observed by RIP-seq (Figure 3, Figure S4D);^3,6,28,65,81^ (ii) Repeat A’s CAWCGGG motif resembles a consensus SRSF1 motif determined by SELEX,^31^ (iii) by proteomics, SRSF1 binding was reduced in the silencing-defective ΔGC-core and AWCG+ΔGG mutants (Figure 1D, Figure 3D); (iv) the GA-rich motifs enriched transcriptome-wide in SPEN RIP-seq peaks closely resemble canonical SRSF1 binding motifs (Figure 2E, Figure 4I);^58,82,83^ and (v) we and others detect physical interactions between SRSF1 and SPEN (Figure 4A,C,I).^59,60,84^

We first performed SRSF1 RIP-qPCR across our panel of *Xist* mutants (Figure 5A). We included wild-type and *Spen*-knockout SM33 cells, which revealed that SRSF1 associates with Repeat A two to three times more strongly in the absence of SPEN (Figure 5A). We observed that SRSF1 association was lost or reduced upon deletion of Repeat A, deletion of the GC-rich cores, or mutations within nucleotides 1 and 17 of the GC-rich cores (Figure 1B, Figure 5A). Conversely, SRSF1 association was unchanged by deletion of the U-rich spacers and increased with mutations between GC-rich-core nucleotides 19 and 25 (Figure 1B, Figure 5A). Levels of SRSF1 and SPEN association across mutants were positively correlated (Pearson’s *r* = 0.47; *p* = 0.09); moreover, excluding mutations within GC-rich-core nucleotides 19 and 25 yielded a strong, highly significant correlation (Pearson’s *r* = 0.84; *p* = 6 x 10^-4^; Figure 5B). The CCUG(21-24) sequence of the GC-rich cores is a canonical SRSF2-binding motif;^85,86^ however, deletion of this sequence did not impact SRSF2-Repeat A association (Figure S6A). Together, these results are consistent with a role for SRSF1 in recruiting SPEN to Repeat A and suggest an SRSF1- and SRSF2-independent role for the 3’ end of the GC-rich cores in recruiting SPEN.

**Figure 5.**
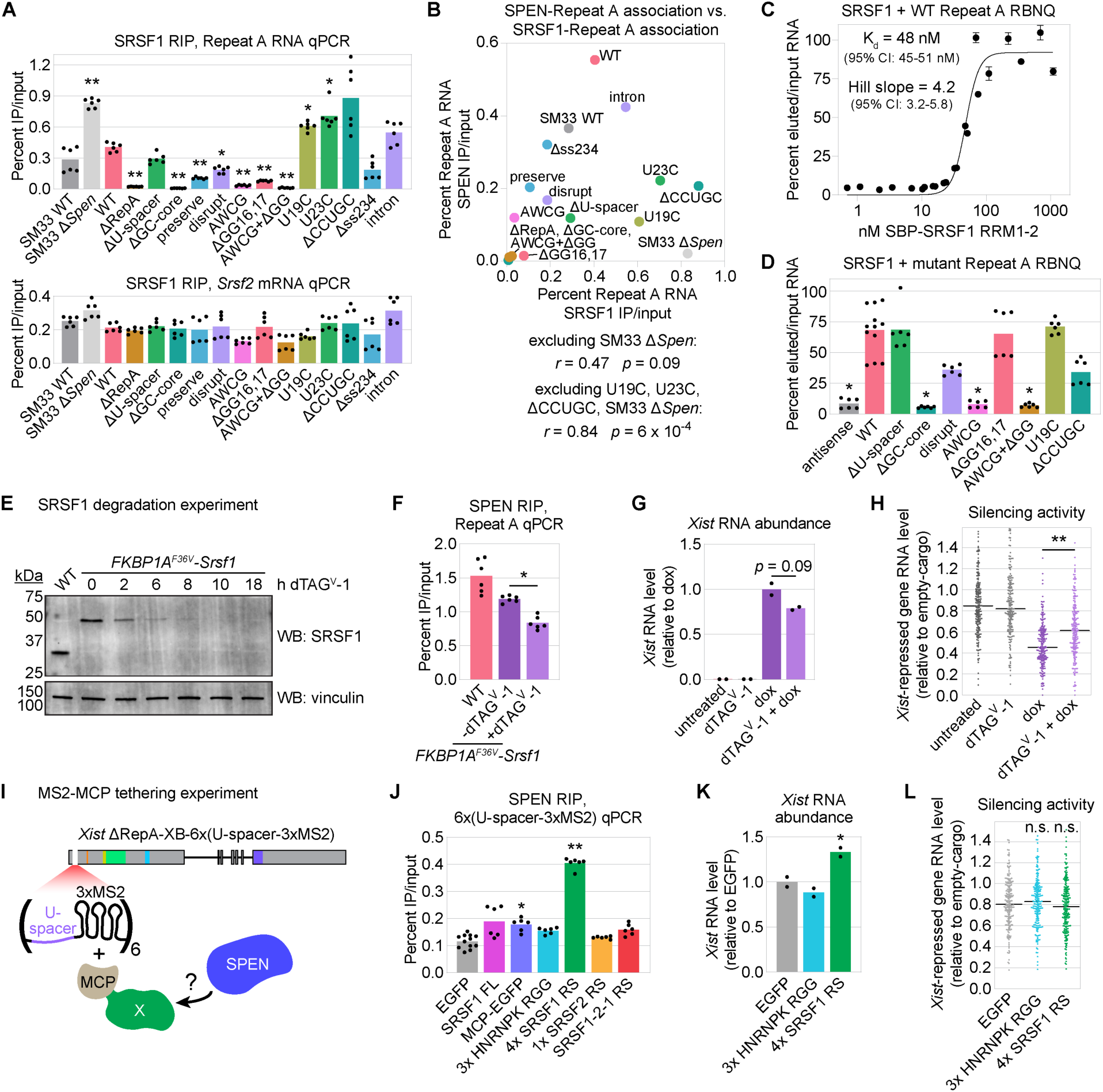
SRSF1 directly binds a conserved Repeat A motif to recruit SPEN to *Xist in vivo*. **(A)** SRSF1-Repeat A association in SM33 or RMCE-*Xist* ESCs by input-normalized RIP-qPCR. *Srsf2* mRNA, control. Dots, qPCR triplicates from two independent experiments. **(B)** SPEN- vs. SRSF1-Repeat A association (RIP-qPCR). *r*, *p*: data fit to linear regression. **(C)** RBNQ with SRSF1 RRM1-2 and Repeat A. Dots, values from two independent experiments. Error bars, standard deviation of qPCR triplicates. **(D)** RBNQ across Repeat A RNAs using fixed SRSF1 RRM1-2 concentration. Dots, qPCR triplicates values, ≥2 independent experiments. **(E)** FKBP1A^F36V^-SRSF1 degradation by dTAG^V^-1 in cells derived from RMCE-*Xist* WT. **(F)** SPEN-Repeat A association in WT or *FKBP1A^F36V^*-*Srsf1* cells by input-normalized RIP-qPCR. Dots, qPCR triplicates from independent clonal lines. **(G)** *Xist* abundance (RNA-seq) in *FKBP1A^F36V^*-*Srsf1* cells, with/without dox and with/without dTAG^V^-1. Dots, independent lines, relative to dox(+), dTAG^V^-1(-) condition. **(H)** *Xist* silencing activity as in Figure 1D,F, for *FKBP1A^F36V^*-*Srsf1* cells treated as in (G). **(I)** MCP-MS2 tethering to assess SPEN recruitment to *Xist* ΔRepA-XB-6x(U-spacer-3xMS2). **(J)** SPEN-6x(U-spacer-3xMS2) association across MCP genotypes by input-normalized RIP-qPCR. Dots, qPCR triplicates from ≥2 independent experiments. **(K)** *Xist* abundance (RNA-seq) across genotypes. Dots, independent experiments. **(L)** *Xist* silencing activity as in Figure 1D,F across MCP genotypes. “n.s.,” not significantly different. *P*-values, two-sided t-tests relative to WT or EGFP unless specified by black bars: (*), *p* < 0.05; (**), *p* < 0.01. See Figure S6.

We next determined the extent to which SRSF1 directly binds Repeat A *in vitro* by performing equilibrium RBNQ with SRSF1’s RNA-binding domains (RRM1-2; Figure S6B). Like SPEN, SRSF1 bound Repeat A cooperatively, yet with a four-fold stronger K_d_ than that of SPEN, despite being ∼1000-fold more abundant in ESCs (Figure 2G, Figure 5C).^87^ Holding the concentration of SRSF1 RRM1-2 constant in binding reactions with Repeat A mutants, we observed patterns similar to *in vivo* SRSF1 association (Figure 5A), with the ΔGC-core, AWCG, and AWCG+ΔGG mutants exhibiting loss of SRSF1 binding (Figure 5D). Interestingly, the ΔGG16,17 mutant, which reduced SRSF1 association by RIP (Figure 5A) and reduced *in vitro* binding in another study,^28^ did not reduce binding by RBNQ (Figure 5D). Our results demonstrate that the direct interaction between SRSF1 and Repeat A is strong and dependent on the single-stranded AWCG5-8 motif present in each GC-rich core.

To directly test whether SRSF1 is necessary for recruiting SPEN to *Xist*, we tagged the 5′ end of both *Srsf1* alleles in RMCE-*Xist* WT cells with *FKBP1A^F36V^* to enable its degradation upon addition of the small molecule dTAG^V^-1 (Figure 5E, Figure S6C).^88–90^ To determine the impact of SRSF1 depletion on SPEN-Repeat A recruitment, we treated *FKBP1A^F36V^-Srsf1* cells with dTAG^V^-1 for 18 hours prior to inducing *Xist* expression. Using SPEN RIP-qPCR, we observed that SRSF1 depletion caused a 50% decrease in SPEN-Repeat A association relative to wild-type cells and a 30% decrease relative to untreated *FKBP1A^F36V^*-tagged controls, indicating that SRSF1 is necessary for optimal SPEN recruitment to Repeat A (Figure 5F). By RNA-seq, acute depletion of SRSF1 caused a mild ∼20% decrease in *Xist* RNA levels and significantly weakened *Xist* silencing activity (Figure 5G,H). While a previous report suggested a role for SRSF1 in *Xist* splicing fidelity,^28^ we observed no aberrant *Xist* splicing events upon SRSF1 depletion (Figure S6D and not shown). These results reveal SRSF1 is necessary to recruit SPEN to *Xist*.

To determine whether SRSF1 is sufficient to recruit SPEN to Repeat A, we devised an *in vivo* tethering assay in which SRSF1 or other proteins were fused to the MS2 bacteriophage coat protein (MCP), and within *Xist*, Repeat A was replaced with a synthetic sequence consisting of six units, each with a natural U-rich spacer followed by three copies of the MCP-cognate MS2 stem-loop (6x(U-spacer-3xMS2); Figure 5I). Initially, we MCP-tagged full length SRSF1 and compared effects to cells expressing untagged EGFP (Figure S6E). Using SRSF1 RIP-qPCR, we confirmed successful tethering of the MCP-SRSF1 fusion construct to the 6x(U-spacer-3xMS2) sequence but did not observe a significant increase in SPEN association relative to EGFP control (*p* = 0.12; Figure 5J, Figure S6F). We speculated that full-length SRSF1 may not robustly recruit SPEN in this system due to steric effects and/or loss of heterotypic cooperativity due to presumed binding sites for multiple proteins in the 25-nt GC-rich core being replaced with monovalent, 95-nt 3xMS2 sequences. We therefore expressed a fusion construct consisting of MCP, an SV40 nuclear localization signal, a 3xFLAG tag, and four concatenated copies of SRSF1’s RS domain, reasoning that the 4x-RS domain might better mimic cooperative cofactor recruitment by the GC-rich core (Figure 5F).^58^ We additionally tested analogously tagged constructs of EGFP, three copies of the HNRNPK arginine-glycine-rich (RGG) domain, one copy of the SRSF2 RS domain, or alternating copies of the SRSF1 and SRSF2 RS domains (Figure S6E). We confirmed tethering of these constructs to *Xist* by FLAG RIP-qPCR (Figure S6G). Strikingly, relative to EGFP, 4xSRSF1 RS recruited four times more SPEN to synthetic Repeat A (*p* = 1 x 10^-4^; Figure 5J). The 4xSRSF1 RS construct nonetheless recruited four-fold less SPEN than wild-type *Xist* and did not induce gene silencing (Figure 5L, Figure S6H). However, 4xSRSF1 RS tethering also caused a 33% increase in levels of *Xist* RNA over EGFP control, likely a consequence of increased SPEN recruitment (Figure 4A, Figure 5K). Consistent with this, across Repeat A mutants, we observed that *Xist* RNA abundance significantly correlated with levels of association with both SRSF1 and SPEN (Figure S6H,I). In summary, using a reconstructed system, we find that the RS domain of SRSF1 is sufficient to increase levels of *Xist* RNA and recruit SPEN to a version of *Xist* lacking its GC-rich cores. Altogether, our results reveal that SRSF1 is a core component of the Repeat A ribonucleoprotein (RNP) complex that recruits SPEN to *Xist*.

### SPEN and SR-protein-binding motifs promote interactions between Repeat A and the m^6^A, RNA turnover, and transcriptional elongation machineries

Our own and previous proteomic analyses detected interactions between SPEN and proteins involved in transcription, splicing, and m^6^A RNA methylation, the last of which has been implicated in the degradation and transcriptional silencing of chromatin-associated RNAs.^20,91,92^ Likewise, SR proteins can interact with RNA polymerase II, the splicing and m^6^A machineries, and the exosome adapters ZC3H18 and YTHDC1, which themselves exhibit association with Repeat A (Figure 3E, Figure S7).^75,79,93–96^ Our observations that Repeat A and the N-terminus of SPEN are SR-protein-binding modules raised the possibility that Repeat A may use SR proteins and SPEN to engage with the transcriptional and RNA processing machineries upstream of or in parallel to inducing transcriptional silencing by chromatin modification.

To investigate this possibility, we used *Xist* oligonucleotide-mediated proximity-interactome mapping (O-MAP) to assess how *Spen* deletion impacts the proteins present within the *Xist* microenvironment (Figure 6A).^97^ We identified 1802 *Xist*-proximal proteins significantly enriched relative to no-probe controls in either RMCE-*Xist* WT or Δ*Spen* cells: 810 exhibited reduced recovery upon *Spen* deletion, 34 exhibited enhanced recovery, and 958 were unchanged (*p* < 0.05, two-sided t-test; Figure 6B, Table S4). We consider the first group “SPEN-dependent” for inclusion in the *Xist* microenvironment and the latter two “SPEN-independent.” The large number of SPEN-dependent proteins may reflect, in part, reduced *Xist* abundance in the absence of SPEN (Figure 4A).^24,26^ Consistent with expectations, the well-characterized SPEN-binding protein RBPJ was SPEN-dependent, whereas proteins that bind *Xist* outside Repeat A, including HNRNPK, HNRNPU, PTBP1, and MATR3, were SPEN-independent (Table S4).^98–100^ Strikingly, consistent with the notion that Repeat A binds SR proteins without SPEN (Figure 5A), 40 of the 51 SR proteins detected as *Xist*-proximal were SPEN-independent,^57^ including SRSF1/3/5/6/7/9, TRA2A/B, U2AF1/2, SCAF1/4/8, SNRNP70, and ACIN1 (Table S4). Conversely, the previously identified Repeat A-interacting proteins RNF20/40 were SPEN-dependent, as were nearly all components of the m^6^A methyltransferase complex (METTL3/14, RBM15, WTAP, ZC3H13, VIRMA; Figure 6B).^10,78^ Additionally, several components of the RNA exosome (EXOSC2/7/8, MTREX) and RNA polymerase II (POLR2A/C/E/H/J) were SPEN-dependent, along with many repressive and activating transcription factors and epigenetic modifiers (Figure 6B, Table S4). In sum, these data suggest that *Xist*’s association with SR proteins is largely SPEN-independent, whereas *Xist*’s association with the m^6^A, RNA exosome, and transcriptional machineries is more SPEN-dependent.

**Figure 6.**
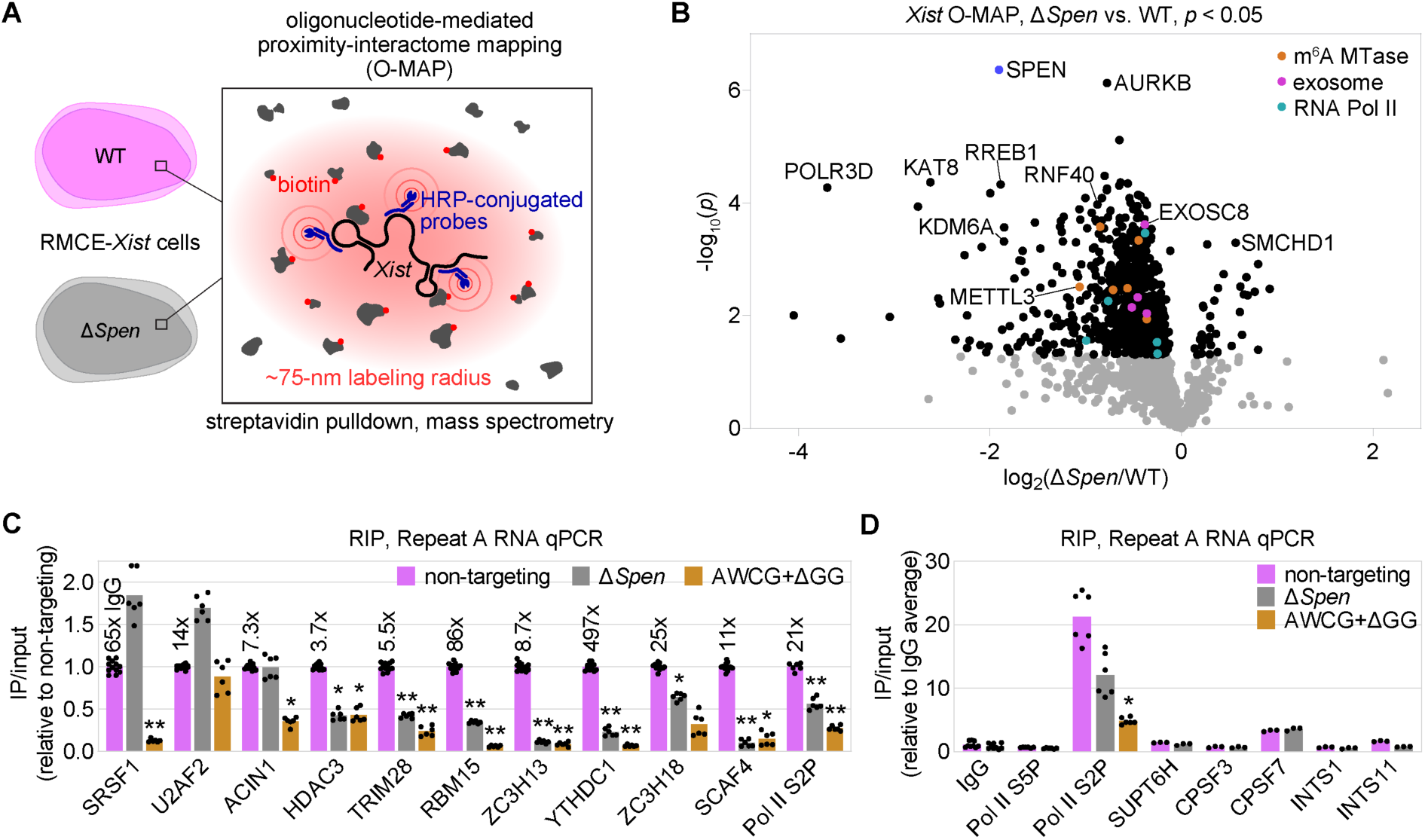
SPEN and SR-protein-binding motifs promote interactions between Repeat A and the m^6^A, RNA turnover, and transcriptional elongation machineries. **(A)** O-MAP schematic; HRP, horseradish peroxidase.^97^ **(B)** Proteins differentially recovered by O-MAP from WT and Δ*Spen* cells (*p* < 0.05, two-sided t-test, n=4). **(C)** Repeat A association (input-normalized RIP-qPCR) in non-targeting-control and Δ*Spen* ESCs derived from RMCE-*Xist* WT, and RMCE-*Xist* AWCG+ΔGG cells. Values relative to non-targeting controls. Dots, qPCR triplicates from ≥2 independent experiments. Values above non-targeting-control bars, experiment-averaged IP/input relative to IgG control. **(D)** RIP-qPCR as in (C), shown relative to IgG. *P*-values, two-sided t-tests relative to WT or non-targeting controls: (*) indicates *p* < 0.05; (**) indicates *p* < 0.01. See Figure S7.

To probe these observations further, we used RIP-qPCR to examine the extent to which proteins involved in RNA metabolism and transcription physically associate with Repeat A in a SPEN- or SR-protein-binding motif-dependent fashion (Figure S7). Consistent with O-MAP and previous RIP (Figure 5A, Figure 6B), the association of SRSF1 with Repeat A was SPEN-independent but lost with the *Xist* AWCG+ΔGG mutant, a pattern similar for the SR protein ACIN1 (Figure 6C). Repeat A association with the SR protein and spliceosome adapter U2AF2 was independent of both SPEN and the AWCG+ΔGG mutation, consistent with its preference for the U-rich spacers (Figure 3D, Figure 6C). For most other proteins, associations with Repeat A were dually dependent on SPEN and Repeat A’s SR-protein-binding motifs. These included HDAC3 and TRIM28, factors previously implicated in silencing by *Xist* and SPEN;^11,20,21,74^ m^6^A methyltransferase complex components RBM15 and ZC3H13; nuclear exosome adapters YTHDC1 and ZC3H18; the transcription-elongation and antitermination factor SCAF4; and elongating (serine-2-phosphorylated) RNA polymerase II itself (Figure 6C). In contrast, initiating (serine-5-phosphorylated) RNA polymerase II; the transcription elongation factor SUPT6H; and components of the CPSF/CFIm and Integrator complexes did not associate with Repeat A beyond IgG controls (Figure 6D). Together, our data demonstrate that SPEN and SR-protein-binding motifs enable Repeat A to associate with multiple factors involved in transcription and RNA processing.

### SPEN represses transcription of autosomal genes by targeting Repeat-A like RNP features

SPEN is an ancient protein that predates the emergence of *Xist* and is essential for development in contexts beyond X-inactivation.^32,101,102^ We therefore hypothesized that SPEN targets genes independently of *Xist* and that its recruitment is mediated by features shared with Repeat A. To identify potential *Xist*-independent targets of SPEN, we used EU-RNA-seq^103,104^ to measure nascent transcription in male (non-*Xist*-expressing) E14 mouse ESCs with or without a ∼40-kb inactivating deletion in *Spen*.^12,27^ Upon *Spen* deletion, the transcription of more genes was upregulated (850) than downregulated (594; adjusted *p* < 0.05, DESeq2^49^), consistent with a repressive role for SPEN (Figure 7A, Table S5). In agreement with previous reports,^17,105^ we found that *Spen* itself was strongly upregulated in the absence of functional SPEN, suggesting that SPEN represses its own transcription. Additionally, twice as many X-linked genes, including *Rlim*, *Mecp2*, and *Huwe1*, were upregulated upon *Spen* deletion than expected by chance (57 vs. 29, *p* = 4 x 10^-8^, chi-squared test), suggesting a sensitivity of X-linked genes to SPEN-mediated repression.

**Figure 7.**
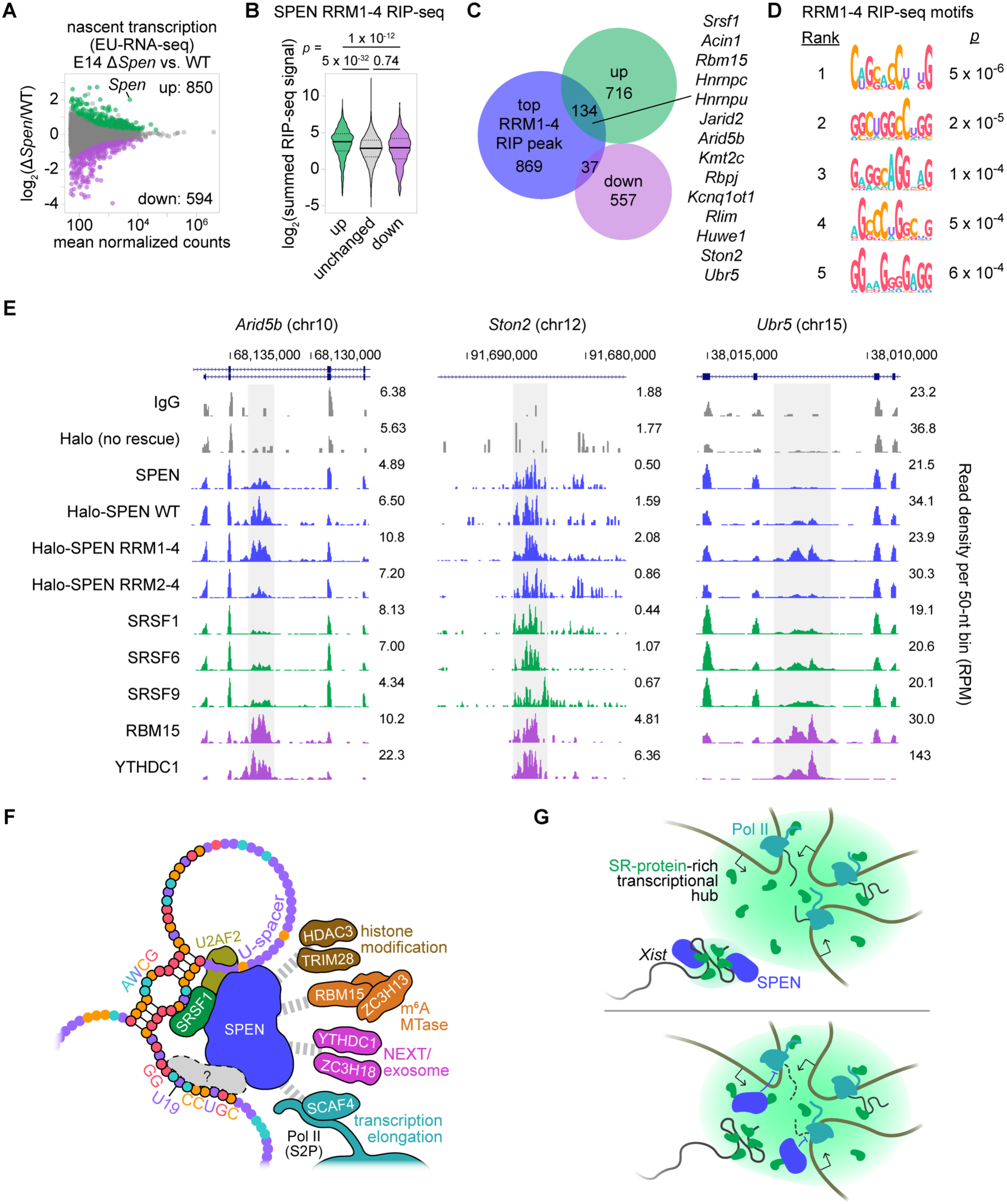
SPEN represses transcription of autosomal genes by targeting Repeat-A like RNP features. **(A)** MA plot of differential nascent transcription between E14 WT and Δ*Spen* cells. **(B)** Violin plot of summed SPEN RRM1-4 RIP-seq signal for genes in each differential expression category. **(C)** Overlaps of genes containing top-1500 SPEN RRM1-4 RIP-seq peaks with up- and downregulated genes in Δ*Spen* vs. WT EU-RNA-seq. **(D)** Top motifs in the 189 RRM1-4 RIP-seq peaks within upregulated genes. **(E)** RIP-seq tracks surrounding peaks of SPEN RRM1-4 (grey) in upregulated genes. **(F)** Model of Repeat A repeats 6 and 7 from structural model in Lu et al.^16,132^ SPEN is recruited to Repeat A predominantly by proteins that bind motifs in the GC-rich core, whereas the U-rich spacers play minor roles. SPEN and GC-rich-core-bound proteins enable association with factors involved in transcriptional elongation and RNA processing. **(G)** Sensing of SR-protein-rich transcriptional hubs by Repeat A and SPEN.

We next investigated whether *Spen*-dependent changes in transcriptional output were connected to recruitment of SPEN. To best capture SPEN-RNA interactions, we analyzed SPEN RRM1-4 RIP-seq data from Δ*Spen* cells because RRM1-4 is more highly expressed than full-length SPEN and lacks silencing domains that could reduce levels of target RNAs (Figure 4I, Figure S5B). We found that transcripts of upregulated genes associated with significantly more SPEN RRM1-4 than those of downregulated or unchanged genes (Figure 7B, Table S5). Likewise, genes containing the top 1500 RRM1-4 RIP-seq peaks (Figure 4I) were 2.5 times more likely to be transcriptionally upregulated than downregulated upon *Spen* deletion (*p* = 3 x 10^-30^, chi-squared test; Figure 7C, Table S5). Intriguingly, among the 134 upregulated genes with a top 1500 RRM1-4 RIP-seq peak (our highest-confidence SPEN targets) were those with roles in *Xist* biology (*Srsf1*, *Rbm15*, *Hnrnpc*, *Hnrnpu*), epigenetic modification (*Jarid2*, *Arid5b*, *Kmt2c*), and NOTCH signaling (*Rbpj*). Like Repeat A, the top RRM1-4 peaks within upregulated genes were enriched in GC- and GA-rich SR-protein-like motifs and SR-protein RIP-seq signal (Figure 7D,E). For example, RRM1-4 peaks within the SPEN-repressed genes *Arid5b*, *Ston2*, and *Ubr5* coincided with RIP-seq signal for SRSF1, SRSF6, and SRSF9; these sites were also enriched in m^6^A machinery association, further resembling Repeat A (Figure 7E, Figure S7). At these same regions, RIP-seq signal for SPEN RRM2-4 was weaker than RRM1-4, consistent with the notion that SPEN is targeted to autosomal transcripts via interactions with SR proteins (Figure 4G,H, Figure 7E).

## DISCUSSION

Our study resolves major unknowns regarding the mechanism by which *Xist* recruits SPEN and reveals what is to our knowledge the first direct mechanistic link between core splicing factors and the coordination of RNA-mediated gene silencing in mammals. We find that the sequences in Repeat A most essential for gene silencing do so by binding canonical splicing factors such as SRSF1, which in turn recruit SPEN. These same sequences are potent suppressors of splicing and, together with SPEN, bridge Repeat A with transcriptional repressors and activators, including the m^6^A machinery and elongating RNA polymerase II itself (Figure 7F).

Our findings indicate that Repeat A is a splicing factor-binding module that recruits SPEN predominantly through bridged interactions with a subset of SR proteins. While SRSF1 is one, our data suggest that yet-to-be-identified proteins also bind the AWCG and downstream motifs in Repeat A, and that these proteins can substitute for or work in concert with SRSF1 to recruit SPEN. SR proteins have overlapping binding specificities and are orders of magnitude more abundant than SPEN in cells.^58,60,72,106^ Thus, it stands to reason that SR proteins assemble on Repeat A prior to recruiting SPEN, likely as *Xist* is transcribed. Moreover, SR proteins and SPEN interact with elongating RNA polymerase II,^59,93–95,107^ findings we corroborated. We speculate that in the context of nascent *Xist*, interactions among SR proteins, SPEN, and the transcriptional machinery are partly if not fully responsible for the role of Repeat A in promoting *Xist* production. Concurrently, nuclear speckles and foci rich in speckle-related proteins are hubs of transcription and splicing.^94,108–111^ Thus, after mature *Xist* is processed and released from its own locus, SR-protein- and SPEN-driven interactions might grant the *Xist* RNP access to actively transcribed regions (Figure 7G).^20^

Once near an actively transcribed region, our data suggest that *Xist* is capable of silencing via pathways parallel to or independent of chromatin modification. Our observation that Repeat A physically associates with elongating but not initiating RNA polymerase II is consistent with work demonstrating that the chromatin-bound fraction of RNA polymerase II is destabilized in a SPEN-dependent manner during X-chromosome inactivation.^112^ Whether SPEN directly destabilizes RNA polymerase II and/or induces degradation of nascent RNA requires further study. While we did not observe association between Repeat A and CPSF/CFIm or Integrator, we did detect associations between SPEN, Repeat A, the m^6^A machinery, and the nuclear exosome, the latter of which degrades chromatin-associated RNAs.^75–77,91,92,113^ In plants, the SPEN homologue FPA induces degradation of nascent RNAs subsequent to transcriptional silencing, supporting the possibility that SPEN-family proteins are involved in clearance of nascent RNA.^114–116^ In either case, a model whereby SPEN represses transcription prior to directing chromatin modification could rationalize why HDAC3, which is bound to X-linked promoters prior to exposure to *Xist*, does not potentiate silencing until after exposure to *Xist*.^81^ Concordantly, by O-MAP, the presence of several proteins within the *Xist* microenvironment, including HDAC3, was SPEN-independent; whereas, by RIP-qPCR, the same proteins exhibited SPEN-dependent associations, suggesting that SPEN strengthens and integrates interactions between *Xist* and specific factors nearby (Figure 7F). These factors may even include other molecules of SPEN pre-engaged with *Xist* target genes: *Rlim*, one of the first and most robustly-silenced X-linked genes,^21^ exhibited SPEN-dependent repression in male ESCs that do not express appreciable levels of *Xist*.

SPEN and SR proteins are deeply conserved, predating the emergence of *Xist* by several hundred million years.^19,117^ We surmise that SPEN’s ability to recognize SR-protein-rich assemblies also predates *Xist*. Our data indicate that, in contexts outside X-inactivation, SPEN maintains homeostatic gene expression by transcriptionally dampening genes whose nascent RNPs are enriched in SR proteins and components of nuclear speckles and spliceosomes. This same mechanism might additionally enable SPEN to silence transcription of SR-protein-rich nascent RNPs upon which the spliceosome has stalled. A splicing-quality-control function could explain SPEN’s role in the silencing of transposons, particularly if they lack efficient splicing signals.^17,116^

By extension, SPEN’s recognition of SR-protein-rich assemblies was likely the evolutionary force that gave rise to Repeat A. Repeat A is replete with GC-rich consensus SR-protein-binding motifs and U-rich spacers that resemble canonical polypyrimidine tracts, which each associate with spliceosome-recruiting proteins, including SRSF1, SRSF2, U2AF1, and U2AF2. Yet, Repeat A lacks consensus 5’ and 3’ splice sites, does not appreciably associate with core spliceosome components (e.g., SNRNP70, SNRPA, or SNRNP200; Figure 3E), and is not spliced itself. Rather, it has a local suppressive effect on splicing (Figure 1H,I). Thus, disruption of splicing near the 5’ end of proto-*Xist*,^118^ either by chance mutation or transposon insertion,^17,119^ followed by expansion of a proto-Repeat A monomer, may represent the path by which SPEN recruitment and transcriptional silencing ability arose in present-day *Xist*.

Genetic and proteomic screens have identified SRSF1, U2AF1, and other SR proteins and RNA-binding proteins as factors essential for the establishment or maintenance of mammalian heterochromatin, yet the mechanisms remain unclear.^120–122^ In that regard, our work may provide a relevant mechanistic paradigm. We have shown that Repeat A nucleates interactions with a core set of RNA-binding proteins that enable *Xist* to recruit more SPEN than any other transcript in the cell to subsequently silence nearby genes. At loci undergoing heterochromatinization, lowly abundant RNAs may nucleate analogous interactions with RNA-binding proteins, which could serve as context-specific adapters that cooperate with DNA-based recruitment elements to stabilize epigenetic modifiers locally on chromatin.

Lastly, our discoveries provide important perspectives on SPEN in development and disease. Haploinsufficiency of *SPEN*, located in the rearrangement-prone 1p36 region of the human genome, causes neurodevelopmental disorders.^32^ *SPEN* is also recurrently mutated in cancers and harbors a mutational profile generally consistent with that of a tumor suppressor.^33–41^ Even missense mutations in *SPEN* are selected against in humans.^42^ Functionally, SPEN regulates NOTCH-, WNT-, growth factor-, and hormone-dependent gene expression, but the mechanisms remain ambiguous.^41,99,101,102,123–130^ In these contexts and others, the possibility that SPEN represses transcription through the sensing of SR-protein-rich assemblies should now be evaluated.

## ACKNOWLEDGMENTS

We thank Brian Golitz, Noah Sciaky, and Andrew Snipes of the UNC CRISPR Screening Facility for sequencing. This work was supported by NIH grants R01GM136819, R01GM121806, and R35GM153293; NSF grant DBI-2228805; and the Yang Family Biomedical Scholar Fund (J.M.C.); NIH grants R01GM138799 (to S.A.S., L.E.D., and D.M.S.), R21CA288806 (to S.E.O), R35GM142864 (to D.D.), R01HL160825 (to D.M.S.), and a Chan-Zuckerberg Institute Collaborative Pairs project award (to D.M.S.). J.B.T. was supported by T32CA217824 and a Marzluff Postdoctoral Fellowship from the RNA Discovery Center. S.B.P. was supported by T32GM135128. S.A.S. was supported by T32GM007750 and the ISCRM Fellows Program. M.K.S was supported by K99GM152778. Table S3 results are in part based upon data from (https://www.cancer.gov/tcga). This work used an EASY-nLC1200 UHPLC and Thermo Scientific Orbitrap Fusion Lumos Tribrid mass spectrometer purchased from NIH-SIG grant S10OD021502 to S.E.O.

## AUTHOR CONTRIBUTIONS

J.B.T., A.P., D.M.S., and J.M.C. conceived the study; J.B.T., A.P., S.A.S, L.E.D., Q.E.E., S.P.B., Z.Z., S.L., D.M.L., N.S., S.N.N., and M.-C.G.B. performed the experiments and/or data analysis; S.E.H., M.K.S., S.-E.O., M.P.G.-P., S.A.S., D.D., and D.M.S. provided reagents and advice; J.B.T., D.M.S., and J.M.C. acquired funding; and J.B.T. and J.M.C. wrote the paper with feedback and appropriate methods contributed by co-authors.

## SUPPLEMENTARY FIGURES AND LEGENDS

**Figure S1. Related to Figure 1.**
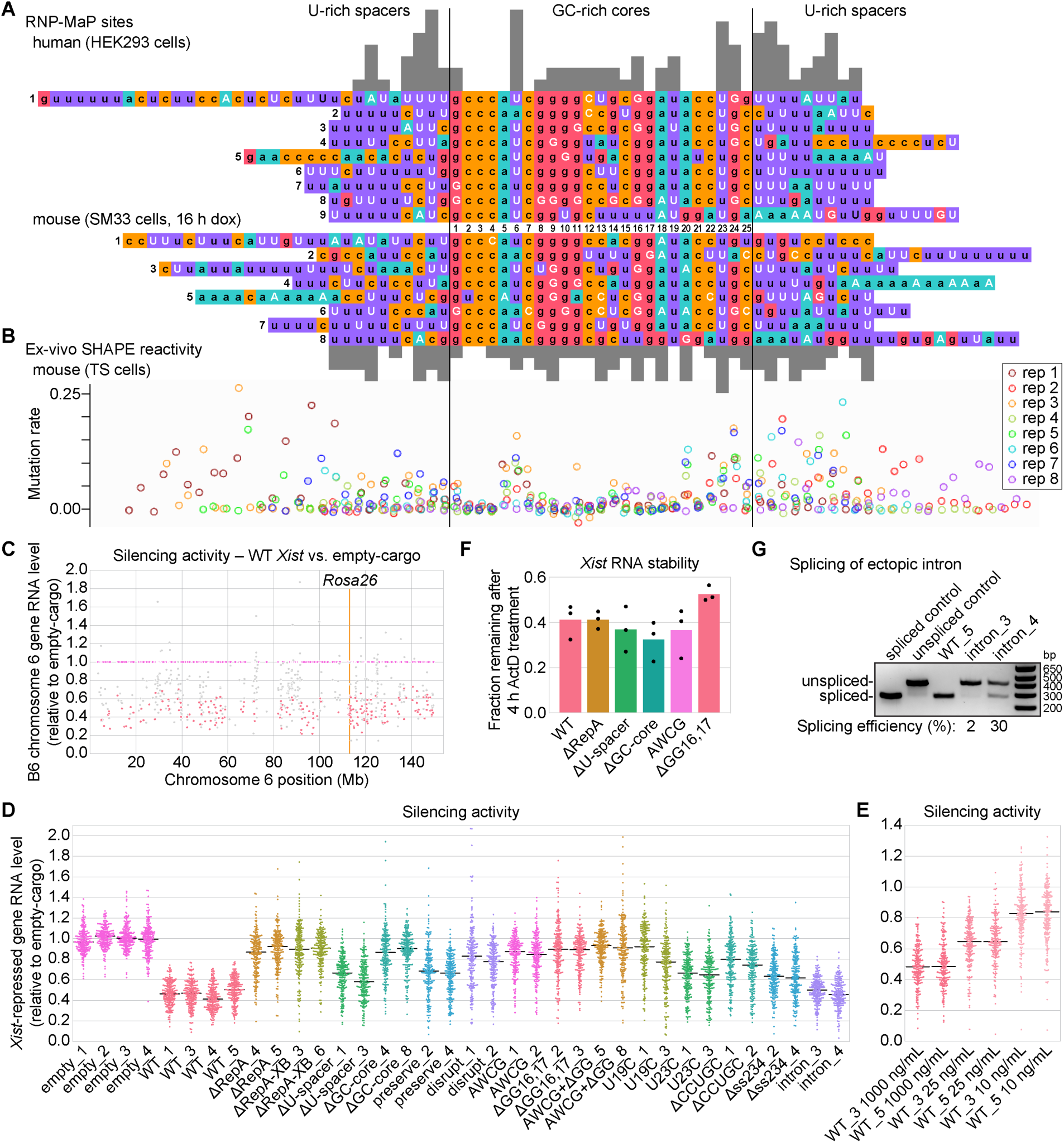
**(A)** Sequences of human (top) and mouse (bottom) Repeat A, with each repeat monomer centered on its conserved GC-rich core sequence. Protein-interacting nucleotides detected by RNP-MaP^7^ are highlighted in white and tallied for each position (grey bars above and below). **(B)** Per-nucleotide ex-vivo SHAPE reactivity for Repeat A in mouse trophoblast stem (TS) cells.^9^ **(C)** *Xist*-induced silencing of genes along the B6 allele of chr6, determined by RNA-seq. The average expression level of each expressed gene in RMCE-*Xist* WT cells (grey and pink dots) is plotted relative to the average expression level in RMCE-empty cells (magenta dots). The 204 genes with significantly altered expression (*p*_adj_ < 0.001; DESeq2)^43^ are depicted by pink dots and designated as “*Xist*-repressed genes.” **(D and E)** Silencing activity for *Xist*-repressed genes in individual clonal cell lines. Average values for each gene in each genotype/condition are shown in Figure 1D,F. Black bars, median values. **(F)** *Xist* RNA stability across WT and Repeat A mutants, using RT-qPCR to measure differences following treatment for 4 h with actinomycin D (ActD). Dots, average values from three independent experiments. **(G)** Semiquantitative PCR using intron-flanking primers to estimate splicing efficiency of the ectopic intron inserted downstream of Repeat A in independent cell lines. Spliced and unspliced controls used RMCE-*Xist* WT and intron plasmids, respectively, as PCR templates.

**Figure S2. Related to Figure 1.**
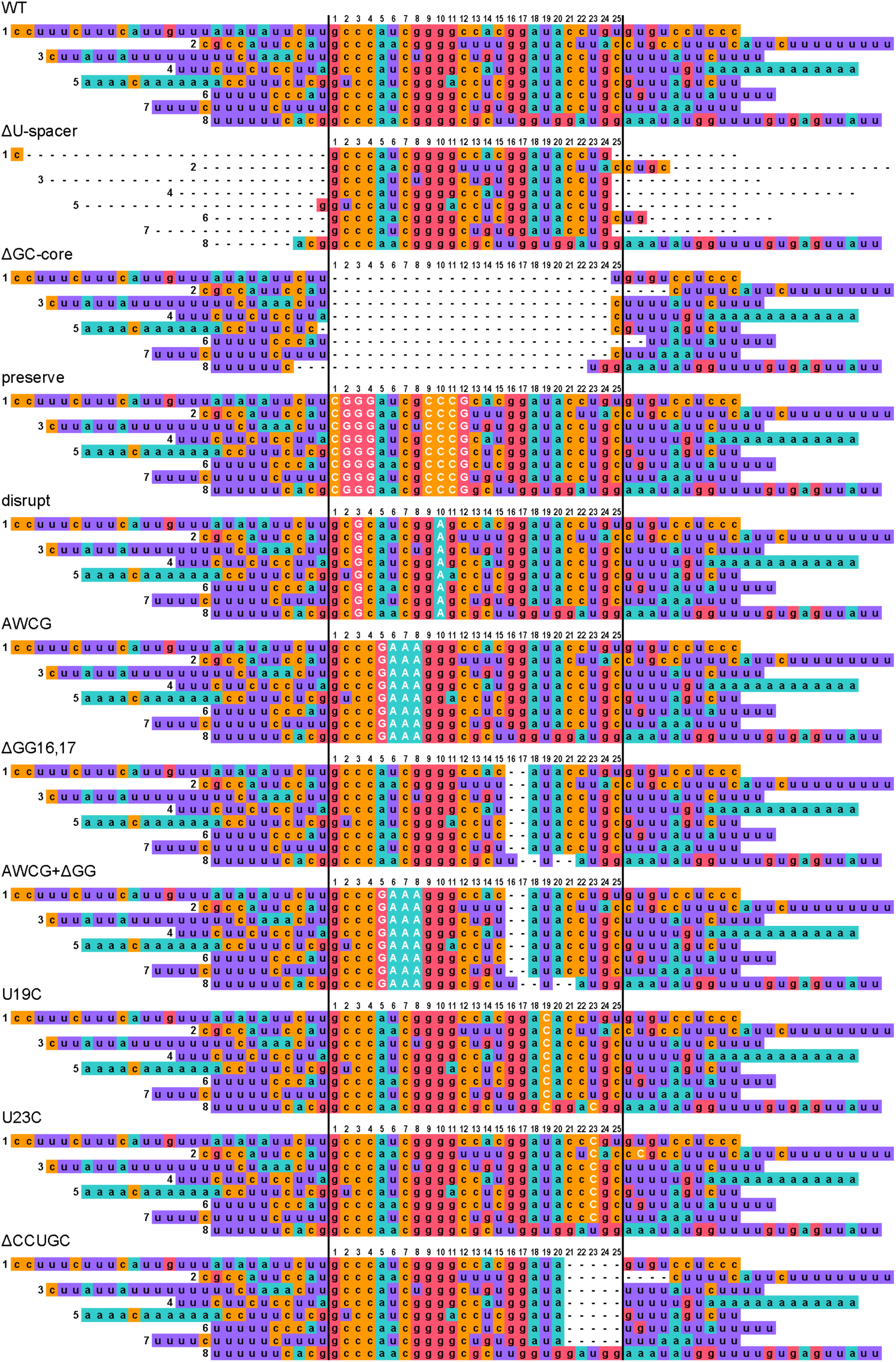
Repeat A mutant sequences, with deletions shown as dashes and substitutions shown in white. Each repeat monomer sequence is aligned and centered on its GC-rich core (flanked by black lines), with position of each GC-rich core nucleotide numbered above.

**Figure S3. Related to Figure 2.**
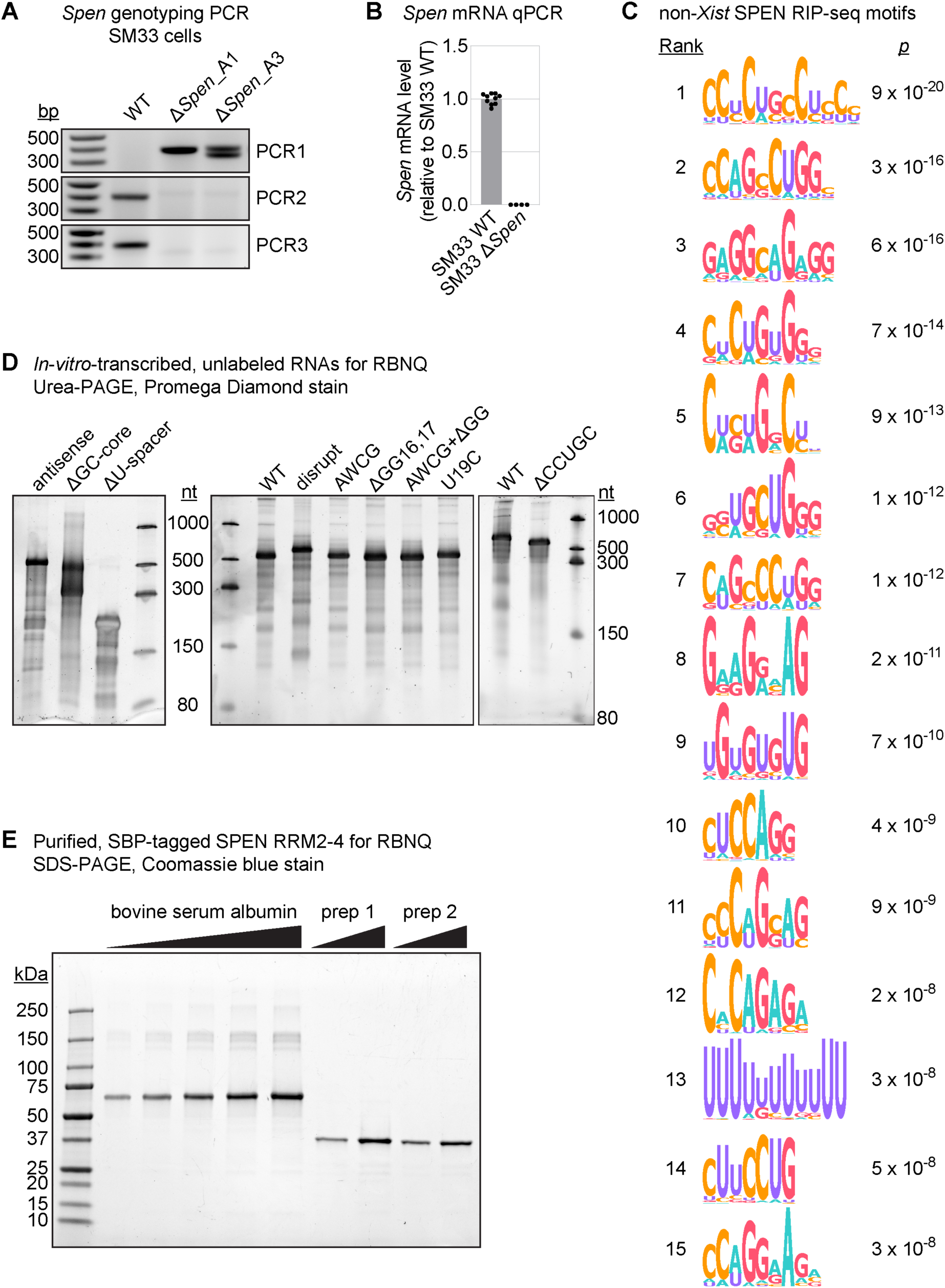
**(A)** Genotyping PCR for independent clonal lines of SM33 Δ*Spen* cells generated by CRISPR-Cas9. PCR1 was performed using primers flanking either side of the deletion; PCR2 and PCR3 were performed using primers flanking the upstream and downstream gRNA target sites. **(B)** RT-qPCR measuring *Spen* mRNA in SM33 WT and Δ*Spen* cells. Values are relative to SM33 WT. Dots represent average qPCR-replicate values for independent cell lines. **(C)** Top 15 most significant sequence motifs enriched in the 1500 most abundant non-*Xist*, non- *Spen* SPEN-specific peaks from SPEN RIP-seq, determined by STREME. **(D)** Promega Diamond-stained urea-PAGE gels of in-vitro-transcribed Repeat A RNAs used for equilibrium RBNQ experiments. **(E)** Coomassie blue-stained SDS-PAGE gel of two independent preps of SBP-tagged SPEN RRM2-4 used for replicates of equilibrium RBNQ experiments.

**Figure S4. Related to Figure 3.**
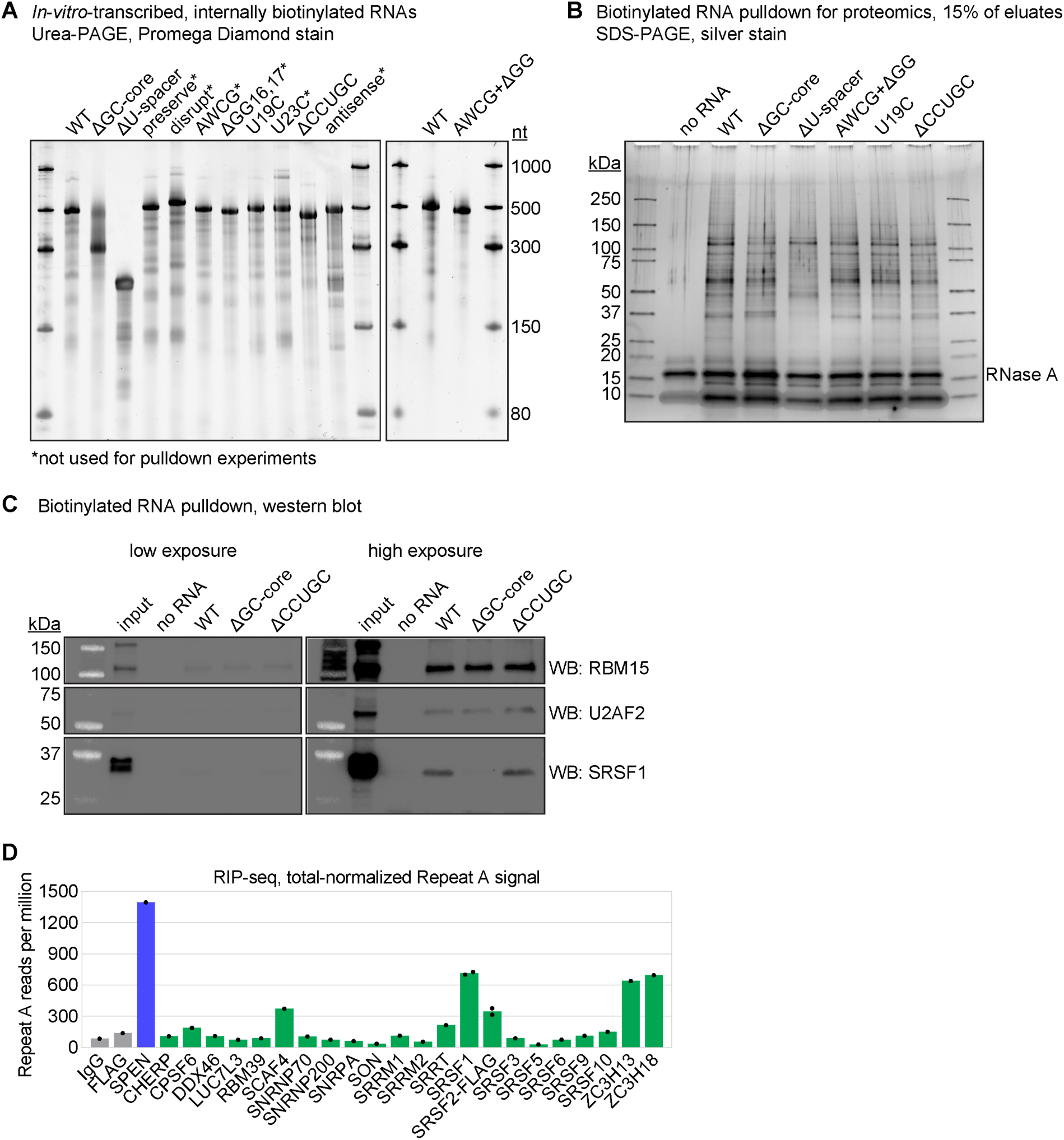
**(A)** Promega Diamond-stained urea-PAGE gels of in-vitro-transcribed, internally biotinylated Repeat A RNAs used for biotinylated RNA pulldown experiments. **(B)** Representative silver-stained SDS-PAGE gel of proteins recovered by biotinylated Repeat A mutant pulldowns or no-RNA control pulldown. The remaining 85% of each sample was analyzed by quantitative proteomics. **(C)** Validation of quantitative proteomics results by biotinylated RNA pulldown and western blotting for the proteins RBM15, U2AF2, and SRSF1. **(D)** Quantification of the proportion of RIP-seq signal in Figure 3E within Repeat A relative to total reads per million. Dots, values from independent experiments.

**Figure S5. Related to Figure 4.**
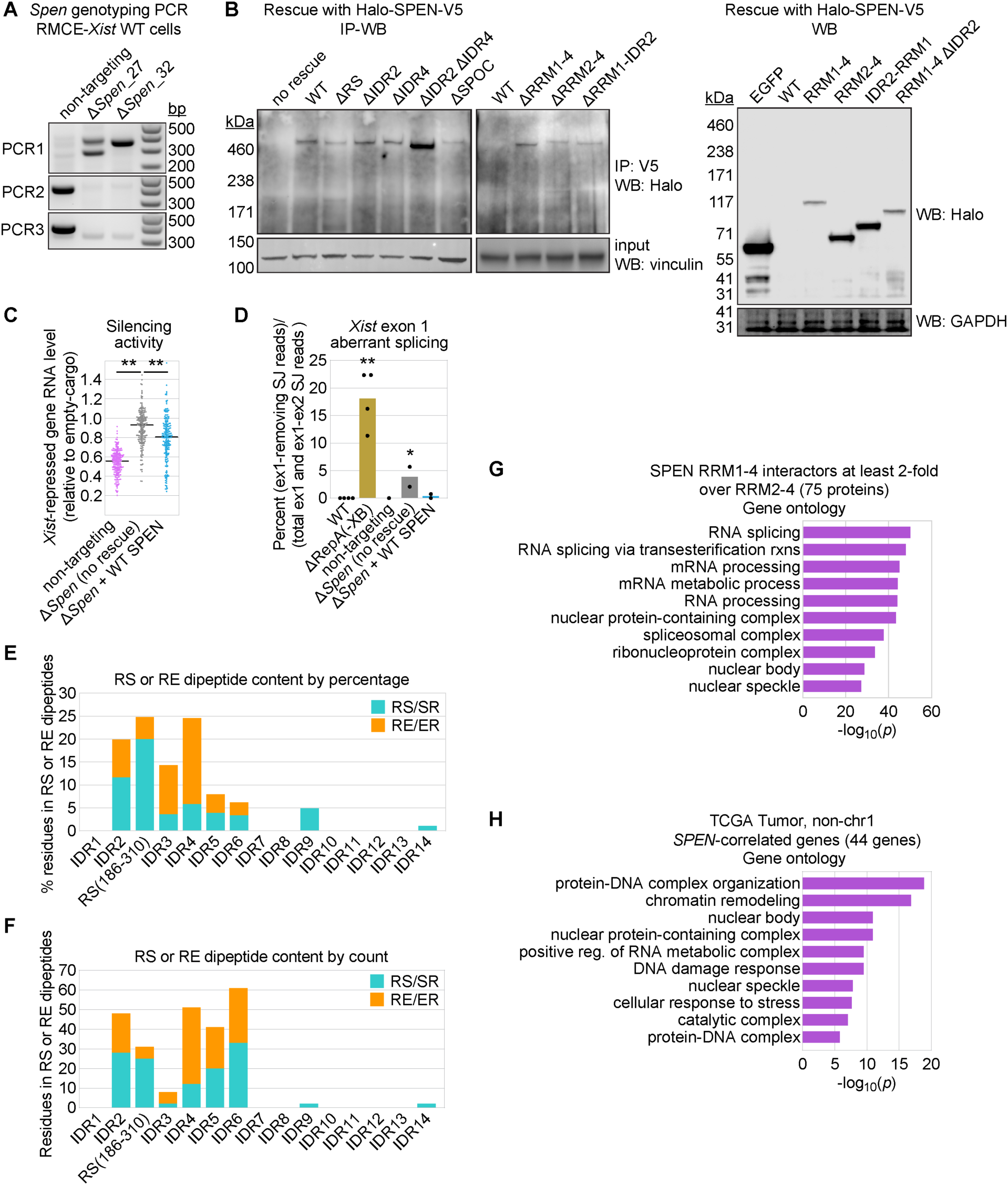
**(A)** Genotyping PCR for independent clonal lines of RMCE-*Xist* WT Δ*Spen* cells or non-targeting gRNA-treated control cells generated by CRISPR-Cas9. PCR1 was performed using primers flanking either side of the deletion; PCR2 and PCR3 were performed using primers flanking the upstream and downstream *Spen* gRNA target sites. **(B)** Confirmation of expression of WT and mutant Halo-SPEN-V5 proteins by IP-WB or WB. **(C)** *Xist* silencing activity determined by RNA-seq, as in Figure 1D,F, for non-targeting control cells and Δ*Spen* cells with and without WT Halo-SPEN-V5 expression. Asterisks, significant differences by two-sided t-test. **(D)** Quantification of exon 1 aberrant splicing for each RMCE-*Xist* cell line (dots), as done in Figure 1I. Asterisks, values significantly different from WT by two-sided t-test. **(E and F)** Quantification of RS/SR and RE/ER dipeptide content in each IDR of SPEN by percentage of residues (E) or overall residue count (F). **(G)** Gene ontology analysis of the 75 proteins recovered with SPEN RRM1-4 two-fold more than with RRM2-4 in two independent experiments, showing the 10 most significantly enriched terms. **(H)** Gene ontology analysis of the 44 non-chromosome 1 (chr1) genes with expression significantly correlated with *SPEN* across tissue types in the TCGA Tumor dataset (FDR < 0.1). An additional 59 chr1 genes were significantly correlated with *SPEN* in this dataset but were omitted from this analysis because their correlations with *SPEN*, a gene on chr1, cannot be decoupled from cancer-associated copy-number variations.

**Figure S6. Related to Figure 5.**
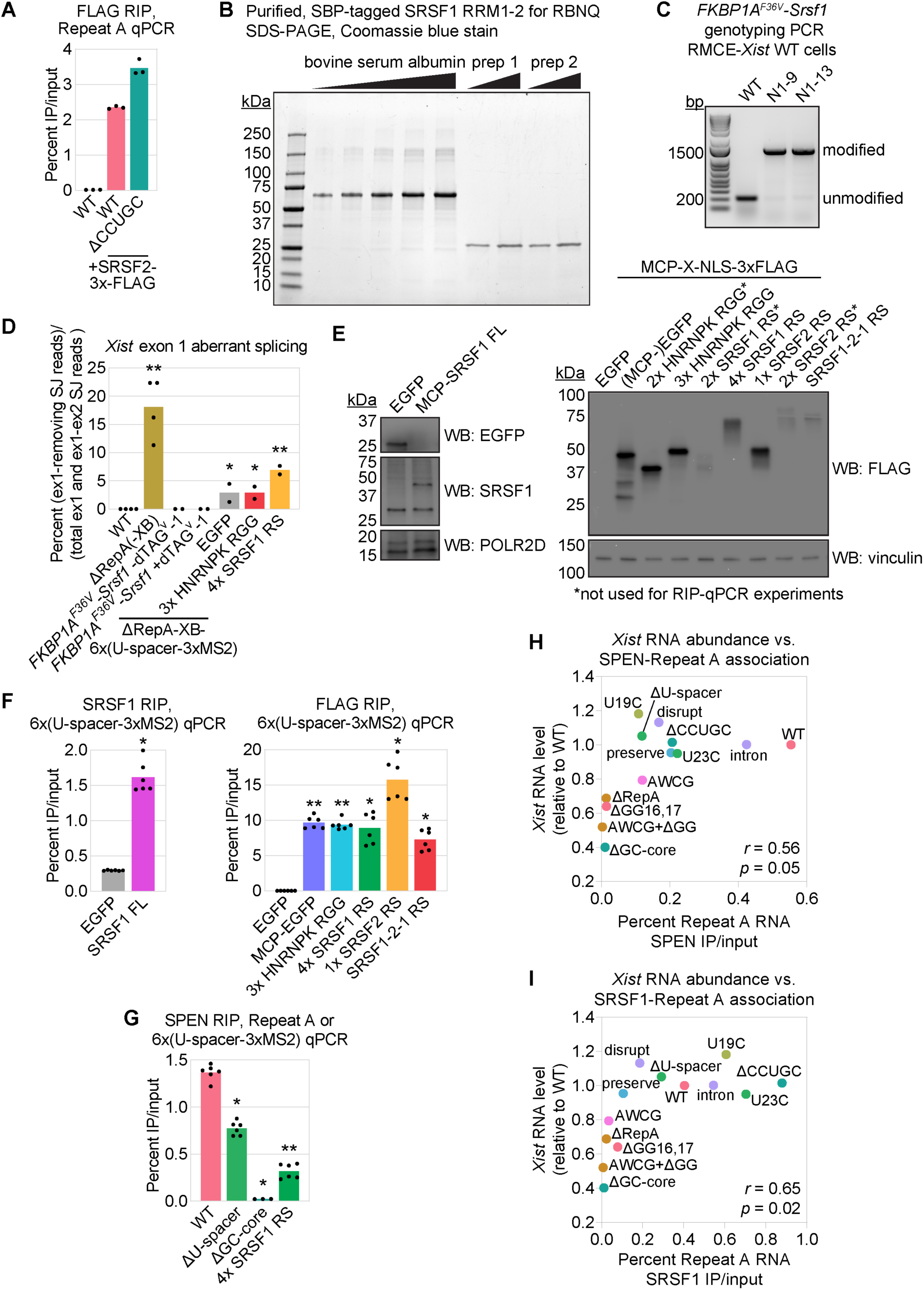
**(A)** Levels of Repeat A recovery by input-normalized FLAG RIP-qPCR from RMCE-*Xist* WT cells expressing no FLAG-tagged construct and RMCE-*Xist* WT and ΔCCUGC cells expressing SRSF2-3xFLAG. Dots, qPCR triplicate values from one experiment. **(B)** Coomassie blue-stained SDS-PAGE gel of two independent preps of SBP-tagged SRSF1 RRM1-2 used for replicates of equilibrium RBNQ experiments. **(C)** Genotyping PCR for parental RMCE-*Xist* WT control cells or independent clonal lines of *FKBP1A^F36V^*-*Srsf1* cells generated by CRISPR-Cas9 and homology-directed repair. PCR was performed using primers flanking either side of the *FKBP1A^F36V^* degron tag insertion. **(D)** Quantification of exon 1 aberrant splicing for each RMCE-*Xist* cell line (dots), as done in Figure 1I. Asterisks, values significantly different from WT by two-sided t-test. **(E)** Western blot confirmation of expression of untagged EGFP or various MCP-tagged proteins in RMCE-*Xist* ΔRepA-XB-6x(U-spacer-3xMS2) cells. **(F)** Confirmation of MCP-tagged protein tethering by input-normalized RIP-qPCR using antibodies against SRSF1 (left) or FLAG (right). Cells expressing untagged EGFP served as a negative control. Dots, qPCR triplicate values from two independent experiments. Asterisks, significantly different from untagged EGFP control by two-sided t-test. **(G)** Levels of SPEN association with WT, mutant, or synthetic Repeat A in RMCE-*Xist* WT, ΔU-spacer, or ΔGC-core cells or RMCE-*Xist* ΔRepA-XB-6x(U-spacer-3xMS2) cells, determined by input-normalized RIP-qPCR. Dots, qPCR triplicate values from two independent experiments (one that included ΔGC-core). Asterisk, significantly different from WT by two-sided t-test (two-sided one-sample t-test for ΔGC-core). **(H and I)** Plots of *Xist* RNA abundance as a function of SPEN-Repeat A association level (H) or SRSF1-Repeat A association level (I). Values of *r* and *p* reflect fit to a linear regression models with the indicated data. The ss234 deletion reduces *Xist* abundance independent of Repeat A and was thus omitted.

**Figure S7. Related to Figure 6.**
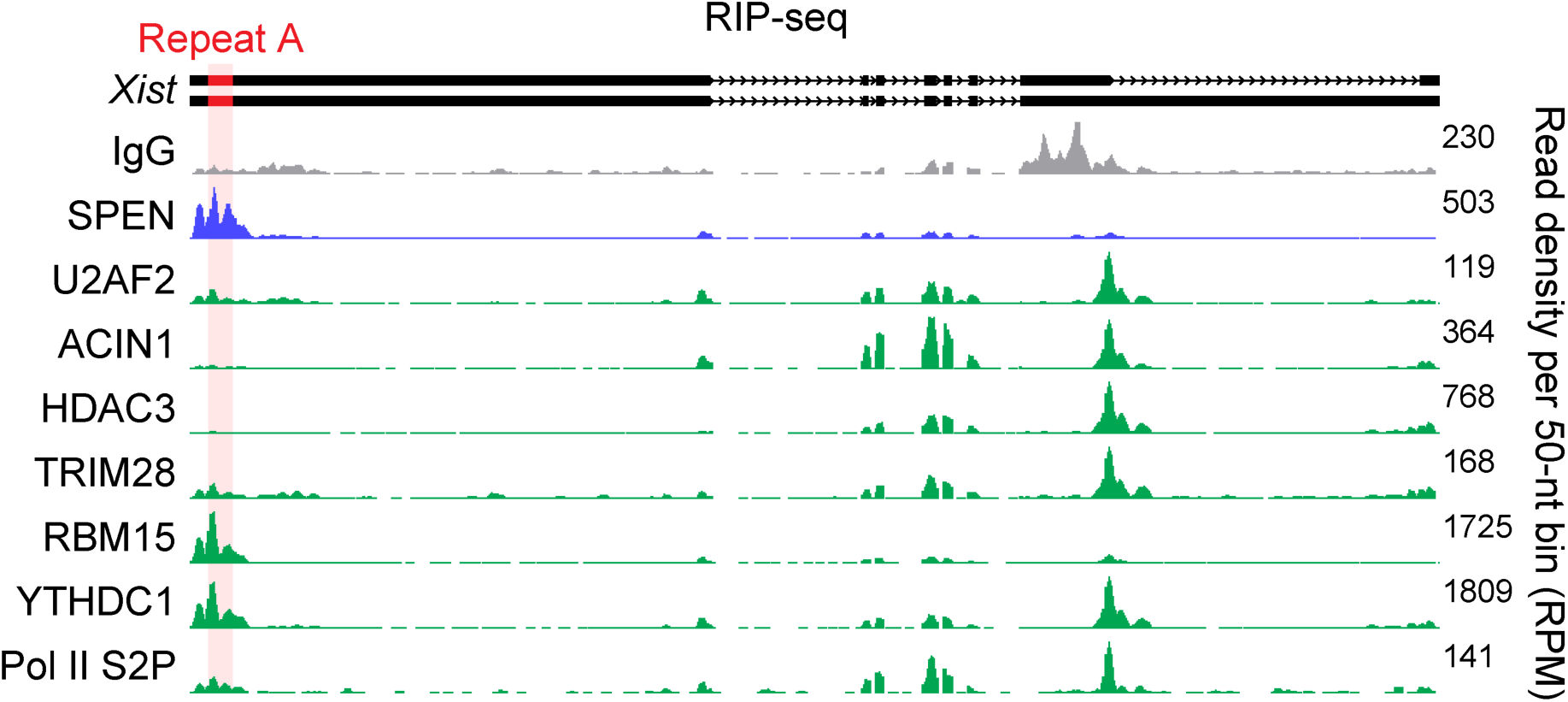
RIP-seq from RMCE-*Xist* non-targeting gRNA control cells, with antibodies used in Figure 6C. IgG (grey) and SPEN (blue) RIP-seq tracks from Figure 3E are included as controls. Note that for some RIPs, enrichment over Repeat A relative to the rest of *Xist* was low; nonetheless, Repeat A was recovered at levels significantly higher than IgG control in input-normalized RIP-qPCR (Figure 6C).

## SUPPLEMENTARY TABLES

**Table S1. Quantitative proteomics of biotinylated Repeat A RNA pulldowns. Related to Figure 3.**

**Table S2. Halo-SPEN IP proteomics. Related to Figure 4.**

**Table S3. S*PEN*-correlated genes by guilt-by-association. Related to Figure 4.**

**Table S4. WT vs. Δ*Spen Xist* O-MAP proteomics. Related to Figure 6.**

**Table S5. E14 WT vs. Δ*Spen* EU-RNA-seq and SPEN RRM1-4 RIP-seq. Related to Figure 7.**

**Table S6. Oligonucleotides and antibodies used in this study. Related to Figures 1-7.**

## MATERIALS AND METHODS

### Cell culture

Male E14 mouse embryonic stem cells (mESCs, kind gift of D. Ciavatta), male B6/CAST F1-hybrid *Rosa26*-RMCE mESCs,^27^ and male SM33 mESCs (with dox-inducible promoter at the endogenous *Xist* locus; kind gift of K. Plath; ^52^), were grown on gelatin-coated plastic dishes in a humidified Thermo Fisher Forma Series II water-jacketed incubator at 37 °C and under 5% CO_2_. Cells were cultured in growth medium consisting of DMEM high glucose plus sodium pyruvate (Gibco 11995-065), 15% ESC-qualified fetal bovine serum (Gibco 26140-079), 0.1 mM non-essential amino acids (Gibco 11140-050), 100 U/mL penicillin-streptomycin (Gibco 15140-122), 2 mM L-glutamine (Gibco 25030-081), 0.1 mM β-mercaptoethanol (Sigma-Aldrich 63689), and 1:500 LIF conditioned media produced from Lif-1Cα (COS) cells (kind gift of N. Hathaway).

Fresh growth medium was supplied daily, and cells were split every two days. To do so, cells were washed once with 1x phosphate-buffered saline (PBS; Corning 46-013-CM), dissociated by adding pre-warmed (37 °C) 0.125% trypsin-EDTA (Gibco 2520072; diluted from 0.25% with 1x PBS) for ∼3-5 min, and quenched by adding two volumes of pre-warmed “standard” quenching medium (DMEM high glucose plus sodium pyruvate containing 10% fetal bovine serum, 100 U/mL penicillin-streptomycin, 2 mM L-glutamine and, 0.1 mM β-mercaptoethanol). Cells were gently dissociated to single-cell suspension using a P1000 pipet before centrifuging the desired number of cells (most typically a 1:6 split) in a conical tube at 200 x g for 5 min at room temp. Supernatants were aspirated, and cells were resuspended in pre-warmed growth medium via serological pipet before adding to tissue-culture-treated plastic dishes (Genesee Scientific) pre-treated with 0.1% (w/v) gelatin (Sigma G9391).

### Plasmid construction

Repeat A mutants were first generated in plasmid pTETRISv1 ^27^ as follows. ∼1.2-kb DNA sequences were synthesized (Genewiz) corresponding to the 5′ end of *Mus musculus Xist* and containing the desired Repeat A mutations (sequences shown in Figure S2). These sequences were inserted into pTETRISv1-*Xist*-2kb WT ^27^ digested with KpnI and NdeI. The insert sequences of the resulting plasmids were verified by Sanger sequencing.

To generate pCARGO-RMCE-*Xist*, ΔU-spacer, ΔGC-core, disrupt, AWCG, AWCG+ΔGG, U19C, U23C, and ΔCCUGC, mutant sequences were PCR-amplified using pTETRISv1-*Xist*-2kb mutants as template and inserted into pCARGO-RMCE-*Xist* digested with NotI and PmlI (cloned by Genewiz).

To generate pCARGO-RMCE-*Xist* ΔRepA (deletion of *Xist* nt 307-732), preserve, ΔGG(16,17), and Δss234 (deletion of *Xist* nt 731-1472), mutant sequences were PCR-amplified from corresponding pTETRISv1-*Xist*-2kb mutants using Q5 High-Fidelity DNA Polymerase (NEB M0491) and primers JT351 and JT352 (Table S6). PCR products (∼0.02 pmol) were inserted into 0.01 pmol NotI-and-PmlI-digested pCARGO-RMCE-*Xist* using NEBuilder HiFi DNA Assembly (NEB E2621). Assembly products were diluted 4-fold with water and transformed into MAX Efficiency Stbl2 cells (Invitrogen 10268019), which were grown overnight at 30 °C with ampicillin selection.

To generate pCARGO-RMCE-*Xist* ΔRepA-XB, pGEM-*Xist*-2kb ^44^ was first modified via site-directed mutagenesis to simultaneously delete Repeat A (*Xist* nt 309-736) and replace it with XmaI and BsiWI restriction sites, generating pGEM-*Xist*-2kb ΔRepA-XB. This was done using NEB Q5 High-Fidelity DNA Polymerase, primers JT353 and JT354, and KLD enzyme mix (NEB M0554). The mutated *Xist* sequence was then PCR-amplified using NEB Q5 High-Fidelity DNA Polymerase and primers JT351 and JT352 and inserted into NotI-and-PmlI-digested pCARGO-RMCE-*Xist* using NEBuilder HiFi DNA Assembly as described above.

To generate pCARGO-RMCE-*Xist* intron, a 328-bp dsDNA gBlock (IDT, JT355, Table S6) was synthesized corresponding to *Xist* sequence containing a 133-bp chimeric β-globin/immunoglobulin intron ^47^ inserted between *Xist* bases 1919 and 1920 (NCBI NR_001463.3). This dsDNA fragment was inserted into PmlI-and-BsaBI-digested pCARGO-RMCE-*Xist* using NEBuilder HiFi DNA Assembly as described above.

To generate pCARGO-RMCE-*Xist* ΔRepA-XB-6x(U-spacer-3xMS2), the following 6x(U-spacer-3xMS2) sequence was first synthesized by Genewiz and inserted into pUC-GW-Kan, including the flanking 5′ XmaI and 3′ BsiWI sites (uppercase, not bold):

CCCGGG**TTTCTTTCATTGTTTATATATTCTT**cgggtggaggatcaccccacccgacacttcacaatcaggg tggaggaacaccccaccctccagacacattacacagaagcaccatcagggcttctg**CTTTTCATTCTTTTTTTTTCTTA TTATTTTTTTTTCTAAACTT**cgggaggacgatcacgcctcccgaatatcggcatgttgcgtggagcatcagcccacgca gccaatcagagtccgcgaagagcatcagccttcgcg**TTTTATTCTTTTTTTCTTCTCCTTA**caggaacagcatcagc gttcctgcccagtacccaaccccatgcagcatcagcgcatgggccccaagaatacatccgagcaccaacagggctcgga**GTTT AGTCTTTTTTTCCCAT**tccatggaccatcaggccatggactctcaccaacaatcgaagcagcatcagcgcttcgaaacact cgagcattgggtggacgatcacgccaccca**TGTTATTATTTTTTTTTCTTTTTCTTTT**gacgagcaccaccagggc tcgtcgttccacgtccaacgggatcacgatcacggatcccgcagctacatcactagcatgcacgatcacggcatgct**TTTAAATT TTTTTTTTCACG**tgggttcagcaccagcgaacccactcctacctcaaaggcaagcaggatcaccgcttgcccattccaacata cctggtacagcatcagcgtaccagCGTACG.

This sequence contains, in order, the endogenous mouse U-rich spacers that precede the GC-rich cores of Repeat A repeats 1, 3, 4, 6, 7, and 8 (bold, uppercase). 3xMS2 sequences, synonymized to prevent recombination (lowercase, underlined), and linkers (lowercase, not underlined) between each U-rich spacer were derived from Addgene plasmid #84561 ^133^. The 6x(U-spacer-3xMS2) sequence was liberated with XmaI and BsiWI and ligated, via NEB Quick Ligase (NEB M2200S), into XmaI-and-BsiWI-digested pCARGO-RMCE-*Xist* ΔRepA-XB.

For expression and purification of proteins N-terminally tagged with glutathione-S-transferase (GST; cleavable) and streptavidin-binding peptide (SBP), sequences coding for SPEN and SRSF1 RRMs were cloned into pDD001 (lab of D. Dominguez) as follows. To generate pDD001-SPEN-RRM2-4, an *Escherichia coli*-optimized coding sequence corresponding to *Mus musculus* SPEN amino acids 335-620 was first synthesized by Genewiz and then PCR-amplified with NEB Q5 High-Fidelity DNA Polymerase and primers JT396 and JT397 and inserted into NcoI-and-NotI-digested pDD001 using NEBuilder HiFi DNA Assembly. To generate pDD001-SRSF1-RRM1-2, SRSF1 RRM1-2 sequence (*Mus musculus* SRSF1 amino acids 1-197) was PCR-amplified from template plasmid pGEX6P-1-SF2 (Addgene #99020, a gift from Honglin Chen, ^134^) using NEB Q5 High-Fidelity DNA Polymerase and primers JT388 and JT390 and inserted into NcoI-and-NotI-digested pDD001 using NEBuilder HiFi DNA Assembly. The sequences of the pDD001 plasmids were confirmed by diagnostic digestion with NcoI and NotI and by Sanger sequencing of the full insert sequences.

The mammalian expression plasmid pCL116(PuroR) – which enables piggyBac-mediated insertion, puromycin-based selection, and doxycycline-induced expression of genes of interest (EGFP in the parental plasmid) – was generated by removing the *HygroR* sequence in pCL116 (“pb-miRE-tre,” from which Addgene plasmid #196087 was derived^135^) with NheI and SnaBI and replacing it with *PuroR* sequence (from pTETRISv1, Addgene plasmid #126033 ^44^) generated by DNA synthesis (Genewiz).

To generate pCL116(PuroR)-SRSF2-3xFLAG, DNA was synthesized (Genewiz) containing mouse-optimized (IDT Codon Optimization Tool, https://www.idtdna.com/pages/tools/codon-optimization-tool) SRSF2 coding sequence followed by a 3xFLAG tag (MDYKDHDGDYKDHDIDYKDDDDK), and this sequence was inserted into pCL116(PuroR)-HA-MCP-SRSF1_v2 (described below) digested with AgeI and SalI (to remove HA-MCP-SRSF1).

To generate pCL116(PuroR)-Halo-SPEN-V5 WT, the Halo-SPEN-V5 sequence from PyPP-CAG-Halo-full-length-Spen-V5 (kind gift of K. Plath ^64^) was PCR-amplified using NEB Q5 High-Fidelity DNA Polymerase and primers JT418 and JT419 and inserted into SalI-and-NcoI-digested (to remove EGFP) pCL116(PuroR) using NEBuilder HiFi DNA Assembly. The RS domain was deleted by Genewiz. To generate the remaining SPEN deletion mutants in pCL116(PuroR)-Halo-SPEN-V5, a strategy was used in which two unique restriction enzymes were used to remove sequence from pCL116(PuroR)-Halo-SPEN-V5 encompassing the desired deletion(s), and NEBuilder HiFi DNA Assembly was used to reconstitute the removed region such that it contained the desired deletion(s). To generate the plasmid backbones for NEBuilder HiFi DNA Assembly, pCL116(PuroR)-Halo-SPEN-V5 was digested with the following pairs of restriction enzymes: NotI and MfeI for RRM1-4, RRM2-4, IDR2-RRM4, and RRM1-4 ΔIDR2; KpnI and MfeI for ΔSPOC; and NotI and PmlI for all others except IDR2-RRM4 and RRM1-4 ΔIDR2, which were generated using NotI-and-MfeI-digested pCL116(PuroR)-Halo-SPEN(RRM2-4)-V5. PCR fragments for NEBuilder HiFi DNA Assembly were generated using with NEB Q5 High-Fidelity DNA Polymerase, pCL116(PuroR)-Halo-SPEN-V5 as template, and primer pairs detailed in Table S6. The following PCR fragments (Table S6) were used to construct each mutant: ΔRRM1-4, fragment 8; ΔRRM2-4, fragments 9 and 10; ΔRRM1-IDR2, fragment 14; ΔIDR2, fragments 1 and 2; ΔIDR4, fragments 3 and 4; ΔIDR2 ΔIDR4, fragments 1, 4, and 5; ΔSPOC, fragments 6 and 7; RRM1-4, fragments 11 and 12; RRM2-4, fragments 12 and 13; IDR2-RRM4, fragment 15; RRM1-4 ΔIDR2, fragments 1 and 16. Assembly reaction products were transformed into NEB high-efficiency 5-alpha competent cells (NEB C2987H), which were grown in LB containing ampicillin. The SPEN mutations by amino acid (aa) residue are as follows: ΔRRM1-4, Δaa1-590; ΔRRM2-4, Δaa335-590; ΔRRM1-IDR2, Δaa1-334; ΔIDR2, Δaa99-330; ΔRS, Δ186-310; ΔIDR4, Δaa694-900; ΔIDR2 ΔIDR4, both Δaa99-330 and Δaa694-900; ΔSPOC, Δaa3478-3644*; RRM1-4, aa1-620 only; RRM2-4, aa335-620 only; IDR2-RRM4, aa99-620 only; RRM1-4 ΔIDR2, aa1-620 with Δaa99-330 (*note that the T2201-A2202 dipeptide in the Uniprot mouse SPEN entry is missing in this plasmid; as a result, SPOC is technically aa 3476-3642 in this plasmid).

To modify the endogenous *Srsf1* loci for dTAG-mediated degradation, two plasmids were constructed, the first containing Cas9 and a guide RNA (gRNA) targeted to the immediate 5’ end of *Srsf1* (gRNA sequence determined with Benchling CRISPR Guide RNA Design tool; https://www.benchling.com/crispr), and the second containing a homology-directed repair (HDR) donor with the PuroR-P2A-2xHA-FKBP1A^F36V^ tag. The first plasmid (PX459_v2-Srsf1-N1-gRNA) was constructed by annealing gRNA-sequence oligos JT459 and JT460 and inserting this sticky-ended dsDNA into BbsI-digested PX459 V2.0 (Addgene #62988, a gift from Feng Zhang,^136^) via NEB Quick Ligase (NEB M2200S). The second plasmid (pGW-Amp-Srsf1-N_term-dTAG-HDR) was created by DNA synthesis (Genewiz) in pGW-Amp, containing the ∼1 kb PuroR-P2A-2xHA-FKBP1A^F36V^ sequence from Addgene plasmid #91793 ^89^, flanked on either side with ∼1 kb of sequence homologous to the endogenous *Srsf1* sequence upstream and downstream of its start codon ^90^.

To generate pCL116(PuroR)-MCP-SRSF1_v2, a coding sequence containing mouse-optimized (IDT Codon Optimization Tool) HA-MCP-SRSF1 was synthesized by Genewiz and inserted into AgeI-and-SalI-digested pCL116(PuroR). The 5’ HA-MCP sequence was taken from Addgene plasmid #31230 ^137^. An XbaI restriction site was included between MCP and SRSF1 to enable cloning of other MCP-tagged constructs.

To generate pCL116(PuroR)-MCP-X-NLS-3xFLAG constructs, pCL116(PuroR)-MCP-SRSF1_v2 was digested with XbaI and SalI to remove SRSF1, which was replaced with synthesized (Genewiz) DNA fragments containing mouse-optimized (IDT Codon Optimization Tool) X-NLS-3xFLAG sequence, where NLS is the SV40 nuclear localization signal (PKKKRKV), 3xFLAG is three copies of a FLAG tag (MDYKDHDGDYKDHDIDYKDDDDK), and X is one of the following: EGFP (sequence from Addgene plasmid #13031), three copies of the mouse HNRNPK RGG domain (aa 247-330), four copies of the mouse SRSF1 RS domain (aa 196-248), one copy of the mouse SRSF2 RS domain (aa 93-221), or one copy of the SRSF1 RS domain followed by one copy of the SRSF2 RS domain followed by one copy of the SRSF1 RS domain.

Large-scale preparations of all pCARGO-RMCE-*Xist* plasmids were purified from MAX Efficiency Stbl2 cells (Invitrogen 10268019) grown overnight at 30 °C with ampicillin selection (100 μg/mL ampicillin sodium salt, Fisher Scientific BP1760), using the NucleoBond BAC100 Maxiprep Kit (Macherey-Nagel 740579). To limit any potential physical shearing during handling, these large (∼29 kb) plasmids below were mixed only by flicking and inverting tubes, taking care to avoid vortexing or extensively mixing by pipet. Pelleted plasmids were resuspended in 120-150 μL of Invitrogen TE Buffer before analyzing purity (A260/A280 > 1.8) with a NanoDrop Lite and DNA concentration with a Qubit 2.0 Fluorometer (dsDNA high-sensitivity or broad-range kit, Thermo Fisher Q32854 and Q32853). Integrity and sequence composition of all pCARGO-RMCE-*Xist* large-scale plasmid preparations were confirmed by diagnostic digestion with KpnI and XhoI, by whole-plasmid Oxford Nanopore sequencing (Plasmidsaurus), and by Sanger sequencing of the regions surrounding Repeat A, Repeat B, and the junction between intron 3 and exon 4, where whole-plasmid sequencing erroneously called an AG-to-AA mutation at the 3′ splice site for several plasmids. Large-scale preparations of all other plasmids were purified from High-Efficiency 5-alpha competent cells (NEB C2987H) or Subcloning Efficiency DH5α competent cells (Invitrogen 18265017) using the PureLink HiPure Plasmid Midiprep Kit (Invitrogen K210004), quantified via Qubit 2.0 fluorometer (dsDNA broad-range kit, Thermo Fisher Q32853), and cloned sequences were confirmed by whole-plasmid Oxford Nanopore sequencing (Plasmidsaurus).

For a complete list of oligonucleotides used in this study, see Table S6.

### Generation of RMCE cell lines

Prior to electroporation, plasmid mixtures were prepared containing 6 μg PB-TRE-Cas9 (for transient hygromycin resistance; Addgene #126029 ^138^), pOG-Cre (for RMCE-cargo integration; Addgene #233265; ^27^; 2.2 μg for mixtures with pCARGO-RMCE-*Xist* and 4.1 μg for mixtures with pCARGO-RMCE-empty for a molar ratio of 1:2.4 for pOG-Cre:pCARGO-RMCE), and 19 μg pCARGO-RMCE-*Xist* (wild-type or mutant) or 6.25 μg pCARGO-RMCE-empty. Each plasmid mixture was ethanol-precipitated, air-dried, and thoroughly resuspended in 10 μL of TE Buffer (Invitrogen 12090015). The 10-μL plasmid mixtures were mixed with 1 million B6/CAST F1-hybrid *Rosa26*-RMCE mESCs (trypsin-dissociated single-cell suspension in 100 μL Neon Buffer R, from Invitrogen MPK10025) and electroporated using a Neon Transfection System (Invitrogen) with one 40-ms pulse of 1000 V before seeding sparsely onto a confluent 10-cm dish of gamma-irradiated drug-resistant (DR4) MEF feeder cells (ATCC SCRC-1045) in growth medium lacking penicillin-streptomycin. Growth medium was not changed the day after electroporation but was replaced daily thereafter. Two days after electroporation, growth medium was replaced with normal growth medium (containing penicillin-streptomycin). Starting three days after electroporation and lasting for four days, cells were selected in growth medium containing 150 μg/mL hygromycin B (Roche 10843555001). Starting seven days after electroporation and lasting for four days, cells were selected in growth medium containing 150 μg/mL hygromycin B and 3 μM ganciclovir (InvivoGen sud-gcv). Nine to eleven days after electroporation, individual colonies (now clearly visible by eye) were picked with aid of a light microscope fitted with a 4X objective, dissociated with 0.125% trypsin-EDTA, and each transferred to a 24-well plate well pre-seeded confluently with gamma-irradiated DR4 MEF feeder cells. Clonal lines of cells were grown until sufficiently confluent (>50%), at which point they were passaged for subsequent genotyping (see “Genotyping PCR” section), preparing cryogenic freeze-down stocks, and introducing the reverse tetracycline-controlled transactivator (rtTA) via plasmid transfection. Every colony screened by genotyping PCR (66 for this study) was positive for having received the cargo cassette.

For introduction of rtTA, four independent RMCE-empty-cargo cell lines, four independent RMCE-*Xist* wild-type cell lines, and two independent RMCE-*Xist* cell lines of each mutant were each seeded at a density of 750,000 cells per well of a 6-well plate. The following day, each cell line was given fresh growth medium and transfected with a plasmid mixture containing 1 μg PB-rtTA (Addgene #126034; ^138^) and 1 μg pUC19-piggyBac transposase ^138^ using Lipofectamine 3000 (Thermo Fisher L3000001). Starting around 24 h after transfection and lasting at least 10 days, cells were selected in growth medium containing 200 μg/mL G418 sulfate (Thermo Fisher 10131035).

### Generation of SM33 Δ*Spen* cell lines

To generate SM33 Δ*Spen* cells, 400,000 SM33 cells were seeded in a 6-well plate well and transfected the following day, via Lipofectamine 3000, with a plasmid mixture containing 0.2 μg of PB-TRE-Cas9 (for transient hygromycin resistance; Addgene #126029; ^138^) and 0.8 μg of a 1:1:1:1 mixture of pX330 plasmids (Addgene #42230; ^139^) expressing Cas9 and 4 different *Spen*-targeting gRNAs.^27^ One day after transfection, cells began 3-day treatment with 150 μg/mL hygromycin B to select for transfected cells. Three days after transfection, cells were split and seeded sparsely (500, 1500, 3000, or 6000 cells) onto confluent 10-cm dishes of gamma-irradiated DR4 MEF feeder cells (ATCC SCRC-1045). Ten days after transfection, individual colonies of cells were picked and subsequently expanded and genotyped (see “Genotyping PCR” section) as done above. Of 48 colonies screened by genotyping PCR for *Spen* deletion, eight contained homozygously deleted alleles.

### Generation of RMCE-*Xist* WT and ΔCCUGC SRSF2-3xFLAG cell lines

To generate cells with doxycycline-inducible expression of SRSF2-3xFLAG, RMCE-*Xist* WT (clone 4) cells and RMCE-*Xist* ΔCCUGC (clone 1) cells already containing rtTA were seeded at a density of 1.5 million cells in a 6-cm dish. The following day, cells were given fresh growth medium and transfected with a plasmid mixture containing 3 μg pCL116(PuroR)-SRSF2-3xFLAG and 1 μg pUC19-piggyBac transposase ^138^ using Lipofectamine 3000 (Thermo Fisher L3000001). Starting around 24 h after transfection and lasting at least 5 days, cells were selected in growth medium containing 2 μg/mL puromycin dihydrochloride (Corning 61-385-RA) and 200 μg/mL G418 sulfate (Thermo Fisher 10131035). These cells were cultured in 2 μg/mL puromycin dihydrochloride and 200 μg/mL G418 sulfate to maintain selective pressure.

### Generation of RMCE-*Xist* Δ*Spen*, non-targeting gRNA control, and Halo-SPEN-V5 rescue cell lines

To generate RMCE-*Xist* Δ*Spen* cells, 500,000 RMCE-*Xist* Δ*HygroR* cells (chosen to enable addition of expression constructs with hygromycin selection; ^135^) were transfected, via Lipofectamine 3000, with a plasmid mixture containing 0.4 μg of pCL116(PuroR) [for transient puromycin resistance] and 1.6 μg of a 1:1:1:1 mixture of pX330 plasmids (Addgene #42230; ^139^) expressing Cas9 and 4 different *Spen*-targeting gRNAs ^27^. In parallel, RMCE-*Xist* non-targeting gRNA control cells were generated by transfecting 500,000 RMCE-*Xist* Δ*HygroR* cells, via Lipofectamine 3000, with a plasmid mixture containing 0.4 μg of pCL116(PuroR) [for transient puromycin resistance], 0.8 μg of PB-TRE-Cas9 (Addgene #126029; ^138^), and 0.8 μg of PB-rtTA(BsmBI)-non-targeting-gRNA (gRNA sequence from TrueGuide sgRNA Negative Control, non-targeting 1 [Invitrogen A35526]; ^140^). Cells were seeded to 6-well plate wells. One day after transfection, cells began 3-day treatment with 2 μg/mL puromycin to select for transfected cells; non-targeting control cells were additionally treated for 1 day with 1 μg/mL dox for transient expression of Cas9. Two days after transfection, cells were split and seeded sparsely (500 or 2000 cells) onto confluent 10-cm dishes of gamma-irradiated DR4 MEF feeder cells (ATCC SCRC-1045). Eight days after transfection, individual colonies of cells were picked and subsequently expanded and genotyped (see “Genotyping PCR” section) as done above. Of 16 colonies screened by genotyping PCR for *Spen* deletion, three contained homozygously deleted alleles.

For introduction of rtTA, two independent RMCE-*Xist* Δ*Spen* cell lines and one RMCE-*Xist* non-targeting gRNA control cell line were each seeded at a density of 500,000 cells per well of a 6-well plate. The following day, each cell line was given fresh growth medium and transfected with a plasmid mixture containing 1 μg PB-rtTA (Addgene #126034; ^44^) and 1 μg pUC19-piggyBac transposase ^44^ using Lipofectamine 3000 (Thermo Fisher L3000001). Starting around 24 h after transfection and lasting at least 10 days, cells were selected in growth medium containing 200 μg/mL G418 sulfate (Thermo Fisher 10131035). These cells and those derived from them were cultured in 200 μg/mL G418 sulfate to maintain selective pressure.

To generate EGFP-Halo or Halo-SPEN-V5 rescue cell lines, RMCE-*Xist* Δ*Spen* (clone 27) cells already containing rtTA were seeded at a density of 400,000 cells per well of a 6-well plate. The following day, cells were given fresh growth medium and transfected with a plasmid mixture containing 1.5 μg pCL116-EGFP-Halo or pCL116(PuroR)-Halo-SPEN-V5 (WT or SPEN mutant) and 0.5 μg pUC19-piggyBac transposase ^44^ using Lipofectamine 3000 (Thermo Fisher L3000001). Starting around 24 h after transfection and lasting at least 5 days, cells were selected in growth medium containing 150 μg/mL hygromycin B (for EGFP-Halo; Roche 10843555001) or 2 μg/mL puromycin dihydrochloride (for Halo-SPEN-V5; Corning 61-385-RA). These cells were cultured in 150 μg/mL hygromycin B or 2 μg/mL puromycin dihydrochloride (as appropriate) and 200 μg/mL G418 sulfate to maintain selective pressure.

### Generation of RMCE-*Xist FKBP1A^F36V^*-*Srsf1* cell lines

To generate RMCE-*Xist FKBP1A^F36V^*-*Srsf1* cell lines, RMCE-*Xist* WT (clone 4) cells already containing rtTA were seeded at a density of 1.5 million cells in a 6-cm dish. The following day, cells were given fresh growth medium and transfected with a plasmid mixture containing 2 μg PX459_v2-Srsf1-N1-gRNA and 2 μg pGW-Amp-Srsf1-N_term-dTAG-HDR using Lipofectamine 3000 (Thermo Fisher L3000001). One day after transfection, all cells were split and seeded onto a confluent 10-cm dish of gamma-irradiated DR4 MEF feeder cells (ATCC SCRC-1045). Two days after transfection, cells began selection with 2 μg/mL puromycin dihydrochloride (Corning 61-385-RA) and 200 μg/mL G418 sulfate (Thermo Fisher 10131035), which lasted at least 10 days. Seven days after transfection, individual colonies were picked and subsequently expanded and genotyped (see “Genotyping PCR” section) as done above. Of 10 colonies screened by genotyping PCR, two contained homozygously edited *Srsf1* alleles.

Note that at the time of picking colonies, many more colonies – at least several hundred – had survived than the anticipated ∼50 to 100 ^90^, so care was taken to pick individual colonies before further growth would have made their isolation infeasible. This deviation from expectation may be attributable to Lipofectamine-mediated transfection being more efficient than the electroporation method in ^90^; as a result, any efforts to repeat this procedure with Lipofectamine-mediated transfection are advised to seed cells at a range of lower densities.

Parallel work to generate C-terminally-tagged *Srsf1*-*FKBP1A^F36V^*cell lines yielded no edited colonies (0/12 screened), possibly a consequence of poor gRNA (GCTCTCGTACATAAGATGAT) performance or cell inviability upon allele editing. Likewise, work to N-terminally tag *Srsf1* using a different gRNA (GCCGCCGCCGCCATGTCGGG) yielded no edited colonies (0/10 screened).

### Generation of RMCE-*Xist* cells for MS2-MCP tethering assay

To generate cells for MS2-MCP tethering, RMCE-*Xist* WT ΔRepA-XB-6x(U-spacer-3xMS2) cells (clone 1 for EGFP and SRSF1-FL; clone 4 for all others) were seeded at a density of 500,000 (clone 1) or 700,000 cells (clone 4) per well of a 6-well plate. The following day, each cell line was given fresh growth medium and transfected with a plasmid mixture containing 0.8 μg of the appropriate pCL116(PuroR) expression plasmid (parental plasmid for EGFP expression), 0.8 μg of PB-rtTA (Addgene #126034; ^44^), and 0.4 μg of pUC19-piggyBac transposase ^44^), using Lipofectamine 3000 (Thermo Fisher L3000001). Starting around 24 h after transfection and lasting at least 10 days, cells were selected in growth medium containing 2 μg/mL puromycin dihydrochloride (Corning 61-385-RA) and 200 μg/mL G418 sulfate (Thermo Fisher 10131035).

### Genotyping PCR

Genomic DNA from individual clonal lines was prepared by first incubating 50-100% confluent 24-well-plate wells of cells (washed once with PBS) with 488 μL of a mixture containing 400 μL gDNA Lysis Buffer (100 mM Tris-HCl pH 8.0, 200 mM NaCl, 5 mM EDTA-NaOH pH 8.0, 0.2% (w/v) SDS), 80 μL of 20 mg/mL Proteinase K solution (Meridian Bioscience BIO-37084), and 8 μL of 5 mg/mL linear acrylamide (Thermo Fisher AM9520). Plates were wrapped in Parafilm and incubated at 55 °C (hybridization oven) for 1-2 h before mixing the contents of each well by pipet, transferring to 1.7-mL centrifuge tubes, and continuing to incubate at 55 °C overnight. The next day, samples were incubated at 100 °C for 1 h before placing on ice, adding 960 μL ice-cold 100% ethanol, inverting several times to mix, and incubating on ice 30 min with occasional inversion. Tubes were then centrifuged at 16,100 x g for 10 min at 4 °C to pellet genomic DNA. DNA samples were washed once with 1 mL ice-cold 80% ethanol, centrifuged again (16,100 x g for 5 min at 4 °C), and DNA pellets were air-dried before resuspending in 100 μL ice-cold TE Buffer (Invitrogen 8019005).

Genotyping PCR was performed using the Apex Taq polymerase kit (Genesee Scientific 42-801B2), in 25-μL reactions containing 1.5 μL genomic DNA, 1x Ammonium Buffer, 1 μM genotyping primers (Table S6), 0.4 mM dNTPs, 1.5 mM MgCl_2_, and 0.1 U/μL (0.5 μL of 5 U/μL stock) Apex Taq polymerase. Reactions were run on a thermal cycler with the following protocol: 95 °C for 3 min, 30 cycles of (95 °C for 30 s, 60 °C for 30 s, 72 °C for 1 min), 72 °C for 5 min. Reaction products were run on a 1% agarose-TAE gel containing ethidium bromide and visualized with a Bio-Rad ChemiDoc MP imager (Ethidium Bromide setting).

### RNA-seq sample preparation

Prior to harvesting RMCE cell lines (with rtTA) were treated with 10-1000 ng/mL doxycycline for 72 hours, with the exception of *FKBP1A^F36V^*-*Srsf1* cells, which were treated for 18 hours with 500 nM dTAG^V^-1 (Fisher Scientific 69-145) or equivalent volume of DMSO (Sigma-Aldrich D2650) as a vehicle control, followed by 26 hours with 1000 ng/mL doxycycline hyclate (Sigma-Aldrich D9891-1G) while maintaining the dTAG^V^-1 or DMSO treatment. Each 6-cm dish of cells was washed twice with 3 mL ice-cold PBS and harvested with 1 mL TRIzol (Thermo Fisher 15596026). Total RNA was prepared via standard TRIzol-chloroform extraction, per manufacturer’s instructions, and resuspended in RNase-free water. RNA concentrations were measured via Qubit 2.0 fluorometer (RNA high-sensitivity or broad-range kit, Thermo Fisher Q32852 and Q10211), and integrity of RNA was confirmed by visualizing rRNA bands on an agarose gel. Single-read, strand-specific (reverse-stranded) libraries for RNA sequencing were prepared using the KAPA RNA HyperPrep Kit with RiboErase (HMR) (Roche 08098140702) and KAPA Unique Dual-Indexed Adapters (Roche 08861919702) or TruSeq adapters, with each library’s input consisting of 9 μL 100 ng/μL total RNA (900 ng) mixed with 1 μL of a 1:250 dilution of ERCC RNA spike-in controls (Thermo Fisher 4456740). RNA fragmentation was performed at 94 °C for 6 min, ligation was performed using 7-μM adapter stocks, and libraries were amplified using 11 cycles of PCR. Libraries were quantified using a Qubit 2.0 fluorometer (dsDNA broad sensitivity kit, Thermo Fisher Q32850), and average fragment sizes were determined by agarose gel electrophoresis. Libraries were pooled and sequenced on an Illumina NextSeq500 platform with a 75-cycle NextSeq 500/550 High Output Kit v2.5 (Illumina 20024906) or Illumina NextSeq1000 platform with a 100-cycle NextSeq 1000/2000 P2 Reagent Kit v3 (Illumina 20046811) or 100-cycle NextSeq 1000/2000 P2 XLEAP-SBS Reagent Kit (Illumina 20100987).

### Quantification of *Xist* levels by RNA-seq

For each sample, RNA-seq reads (fastq files) were aligned with STAR (v2.7.9a, ^141^) to the mouse GRCm38 (mm10) genome (with reference to GTF file gencode.vM25.basic.annotation.gtf from https://www.gencodegenes.org/mouse/release_M25.html ^142^), using multi-sample two-pass alignment that incorporated novel splice junctions discovered among all samples in a given experiment ^141^. STAR was run with the option “--outFilterMultimapNmax 1” to consider only uniquely mapping reads. For each sample, reads mapping to *Xist* were counted with featureCounts (Subread v2.0.4, ^143^) and normalized by dividing by the total number of reads uniquely mapped by STAR.

### Allelic RNA-seq, differential gene expression analysis, and measurement of *Xist* silencing activity

Analysis was performed as described previously ^43^. RNA-seq reads were aligned to mm10, and independently, also aligned to a version of mm10 that had been modified to incorporate CAST single-nucleotide polymorphisms (SNPs) downloaded from the Sanger Mouse Genomes Project on 7/30/2020 ^144^. Alignment was performed with STAR (v2.7.9a, ^141^), using multi-sample two-pass mapping and the option “--outFilterMultimapNmax 1” to consider only uniquely mapping reads. Using a custom Perl script (intersect_reads_snps16.pl) and a file containing a list of CAST SNPs (downloaded from the Sanger Mouse Genomes Project on 7/30/2020, ^144^), aligned reads in the B6 and CAST SAM files were parsed to identify reads clearly originating from either the B6 or CAST allele (i.e. reads that overlap at least 1 B6/CAST SNP). Reads marked as either B6 or CAST were then assigned to genes using a custom Perl script (ase_analyzer10.pl) and the GTF file gencode.vM25.basic.annotation.gtf ^142^. The number of B6-specific reads for each gene from each sample were then compiled.

DESeq2 (v1.32.0, ^49^) was used to identify the set of B6 chr6 genes significantly repressed by wild-type *Xist*. Prior to running DESeq2, a pre-filtering step was used in which genes with fewer than an average of five B6 reads per sample (among four wild-type *Xist* and four empty-cargo control samples) were excluded. Using R, DESeq2 was run to compare the wild-type *Xist* samples against the empty-cargo control samples. Default settings were used except selecting a significance threshold of p_adj_ < 0.001, which yielded 204 chr6 genes significantly repressed by WT *Xist*.

To assess *Xist* silencing activity for each sample, B6-specific reads were compiled for each gene as described above. Using Python, the B6-specific reads for each gene were normalized by dividing by the average number of empty-cargo B6-specific reads for that gene; therefore, a value of 1 represents no silencing, and a value of 0 represents complete silencing. The distributions of empty-cargo-normalized expression values for the 204 wild-type *Xist*-repressed genes were visualized by generating swarmplots with the Seaborn package in Python ^145^.

### *Xist* splicing analysis by RNA-seq

RNA-seq reads were aligned to mm10 using STAR (v2.7.9a, ^141^), using multi-sample two-pass mapping and the option “--outFilterMultimapNmax 1” to consider only uniquely mapping reads. The option “--outSAMtype BAM SortedByCoordinate” were included to output alignments in the BAM format. For memory purposes, Samtools (v1.17, ^146^) was used to generate smaller BAM files containing only chrX reads as well as corresponding index (.bai) files.

ChrX BAM files and corresponding index files were loaded into IGV (v2.8.9, ^147^), and the viewer was navigated to the *Xist* locus. Sashimi plots were generated to visualize strand-specific (reverse-stranded, with “forward” option) junction-spanning reads, setting “Junction coverage minimum” to a value of 2 to remove spurious junction-spanning reads. Extents of aberrant splicing for each sample were calculated by dividing the number of aberrant, exon-1-removing junction-spanning reads by the total number of junction-spanning reads within exon 1 and those spanning exons 1 and 2.

### RNA immunoprecipitation (RIP)

After treatment with 1000 ng/mL doxycycline hyclate (24 h for SM33 cells, 72 h for all RMCE cells except 26 h for *FKBP1A^F36V^*-*Srsf1* cells following 18 h pre-treatment with 500 nM dTAG^V^-1), cells were crosslinked with formaldehyde in 10- or 15-cm dishes as follows. Each 10- (15)-cm dish was washed twice with 6 (10) mL ice-cold PBS before adding 8 (12) mL ice-cold, freshly prepared 0.3% (w/v) formaldehyde (Thermo Scientific Pierce 28906) in PBS and incubating at 4 °C for 30 min. Dishes were then moved to room temp and quenched by adding 1 (1.5) mL 2 M glycine to each dish, gently tilting to mix completely, and incubating for 5 min. Each dish was washed twice with 6 (10) mL ice-cold PBS before harvesting cells by scraping into 5 mL ice-cold 0.1% Tween 20 (Fisher Scientific BP337) in PBS and transferring the cell suspensions to pre-chilled conical tubes on ice. Cells were pelleted by centrifugation (1200 x g, 5 min, 4 °C), supernatants were carefully aspirated, and cell pellets were gently resuspended by pipet in a volume of ice-cold PBS to yield a solution of approximately 10 million cells per mL. (At this point, the sizes of all cell pellets within an experiment were judged relative to one another to ensure that an equivalent number of cells would be used in downstream analyses. For reference, 90% confluent 15-cm dishes routinely yielded approximately 75 million cells.) To pre-chilled 1.7-mL tubes on ice, 1 mL (10-million-cell) aliquots of frequently, gently mixed cell suspension was added. Cells were pelleted by centrifugation (1200 x g, 5 min, 4 °C), supernatants were carefully aspirated, and tubes were snap-frozen in liquid nitrogen before storing at −80 °C.

For each immunoprecipitation, 25 μL well-mixed slurry volume of Protein A/G PLUS-Agarose beads (Santa Cruz sc-2003) was washed three times with 1 mL room-temp 0.5% (w/v) bovine serum albumin (BSA; Fisher Scientific BP1600) in PBS. For each wash, 1.7-mL tubes were gently inverted three times and pelleted by centrifugation (2000 x g, 1 min, room temp), and supernatants were removed carefully, leaving ∼10-20 μL to ensure no loss of beads. To washed beads, 300 μL 0.5% (w/v) BSA in PBS was added, followed by an amount of antibody specified in Table S6. Tubes were incubated overnight at 4 °C with end-over-end rotation.

The following day, formaldehyde-crosslinked cell pellets (10 million cells) were thawed on ice and resuspended with 10 gentle up-down strokes of a P1000 pipet in 508 μL ice-cold RIPA Buffer (50 mM Tris-HCl pH 8.0, 150 mM KCl, 5 mM EDTA-NaOH pH 8.0, 1% [v/v] Triton X-100 [Fisher Scientific BP151], 0.5% [w/v] sodium deoxycholate, 0.1% [w/v] sodium dodecylsulfate) containing the following components added immediately before use: 0.5 μL per mL 1 M dithiothreitol (DTT), 10 μL per mL protease inhibitor cocktail (Sigma P8340), and 5 μL per mL 20 U/μL SUPERase-In RNase inhibitor (Invitrogen AM2696). Keeping tubes on ice, each cell suspension was sonicated twice for 30 s with a Sonics Vibra-Cell 130-watt VCX 130 probe-tip sonicator (30% amplitude output), with a 1-min rest on ice between sonication pulses. Tubes were centrifuged (15000 x g, 15 min, 4 °C), and 510 μL of each supernatant (cleared cell lysate) was transferred to a new pre-chilled tube on ice and mixed with 510 μL ice-cold fRIP Buffer (25 mM Tris-HCl pH 7.5, 150 mL KCl, 5 mM EDTA, 0.5% [v/v] IGEPAL CA-630 [Fisher Scientific ICN19859650]) containing the following components added immediately before use: 0.5 μL per mL 1 M DTT, 10 μL per mL Sigma protease inhibitor cocktail, and 5 μL per mL 20 U/μL SUPERase-In. A 25-μL aliquot of each 1020-μL cleared cell lysate mixture was taken and stored at −20 °C to be processed later as “5% input” samples.

The antibody-bound beads from the day before were washed twice with 1 mL ice-cold fRIP Buffer; for each wash, 1.7-mL tubes were gently inverted three times and pelleted by centrifugation (2000 x g, 1 min, 4 °C), and supernatants were carefully removed, leaving ∼10-20 μL to ensure no loss of beads. To each tube of washed beads, 490 μL of an ice-cold 1:1 mixture of RIPA Buffer and fRIP Buffer containing freshly added 1 M DTT (0.5 μL per mL), Sigma protease inhibitor cocktail (10 μL per mL), and 20 U/μL SUPERase-In (5 μL per mL) was added. To each tube, 490 μL of appropriate cleared cell lysate mixture (equivalent to ∼5 million cells) was added, and tubes were incubated overnight at 4 °C with end-over-end rotation. In the case of the SPEN RIP-qPCR experiments in Figure 2B, the remaining ∼500 μL cleared cell lysate mixtures were snap-frozen in liquid nitrogen and stored at −80 °C to use later in the SRSF1 RIP-qPCR experiments in Figure 5A, using the same sample-matched input samples for normalization.

The next day, beads were pelleted by centrifugation (2000 x g, 1 min, 4 °C), and supernatants were carefully removed. The beads were washed once with 1 mL ice-cold fRIP Buffer and resuspended in 1 mL ice-cold Pol II ChIP Buffer (50 mM Tris-HCl pH 7.5, 140 mM NaCl, 1 mM EDTA-NaOH pH 8.0, 1 mM EGTA-NaOH pH 8.0, 1% [v/v] Triton X-100, 0.1% [w/v] sodium deoxycholate, 0.1% [w/v] sodium dodecylsulfate) before carefully transferring to a new 1.7-mL tube, ensuring no loss of beads. Samples were rotated end-over-end at 4 °C for five min, pelleted by centrifugation (2000 x g, 1 min, 4 °C), and supernatants were carefully removed. Samples were washed twice more with 1 mL Pol II ChIP Buffer, once with 1 mL High-Salt Pol II ChIP Buffer (50 mM Tris-HCl pH 7.5, 500 mM NaCl, 1 mM EDTA-NaOH pH 8.0, 1 mM EGTA-NaOH pH 8.0, 1% [v/v] Triton X-100, 0.1% [w/v] sodium deoxycholate, 0.1% [w/v] sodium dodecylsulfate), and once with 1 mL LiCl Buffer (20 mM Tris pH 8.0, 1 mM EDTA-NaOH pH 8.0, 250 mM LiCl, 0.5% [v/v] IGEPAL CA-630, 0.5% [w/v] sodium deoxycholate); each wash included a 5-min end-over-end rotation at 4 °C. In the final wash, samples were transferred to a new 1.7-mL tube. The final wash was carefully removed, first with a P1000 pipet and then with a P20 pipet, to ensure no loss of beads while leaving a volume of ∼10 μL LiCl Buffer or less on the beads.

Input-sample Reverse Crosslinking buffer was prepared on ice by mixing, for each sample, 33 μL 3X Reverse Crosslinking Buffer (3x PBS, 6% [w/v] N-lauroylsarcosine, 30 mM EDTA-NaOH pH 8.0), 41 μL RNase-free water, 5 μL 100 mM DTT, 1 μL 20 U/μL SUPERase-In, and 10 μL 20 mg/mL proteinase K (Thermo Fisher 25530049). Bead-sample Reverse Crosslinking buffer was prepared on ice by mixing, for each sample, 33 μL 3X Reverse Crosslinking Buffer (3x PBS, 6% [w/v] N-lauroylsarcosine, 30 mM EDTA-NaOH pH 8.0), 66 μL RNase-free water, 5 μL 100 mM DTT, 1 μL 20 U/μL SUPERase-In, and 10 μL 20 mg/mL proteinase K. The 25-μL 5% input samples were thawed on ice and mixed thoroughly by pipet with 90 μL Input-sample Reverse Crosslinking buffer. To washed beads, 115 μL bead-sample reverse crosslinking buffer was added forcefully to resuspend the beads, but the mixture was not otherwise mixed. To reverse crosslinks, samples were incubated for 1 hr at 42 °C, 1 hr at 55 °C, and 30 minutes at 65 °C. During these incubations, bead-containing samples had their beads resuspended every 15 to 30 min by forcefully ejecting ∼100 μL of the supernatant onto the settled beads, a practice done to avoid losing beads to the inside of pipet tips. After heating, 1 mL TRIzol was added to each sample, and samples were inverted several times to mix thoroughly and stored at −80 °C.

The TRIzol-containing samples were thawed at room temp, 200 μL chloroform was added to each, and samples were inverted and vortexed for 3 to 5 s (or alternatively, shaken vigorously all together in a “tube rack sandwich” for 15 s), and centrifuged at 15,000 x g for 15 min at 4 °C. A 580-μL volume of the aqueous phase was mixed with 580 μL 100% ethanol, and samples were inverted, vortexed, and pulsed down before applying (in two steps to accommodate the entire volume) to Zymo-Spin IC Columns (from RNA Clean & Concentrator-5 Kit, Zymo R1013) and centrifuging (16,100 x g, 30 s, room temp). A 400-μL volume of RNA Wash Buffer (Zymo R1013) was added to each column and centrifuged (16,100 x g, 30 s, room temp). To each column, a 40-μL volume of a mixture containing 5 μL DNase I (1 U/μL) and 35 μL DNA Digestion Buffer (Zymo R1013) was added; samples were incubated at room temp for 20 min. A 400-μL volume of RNA Prep Buffer (Zymo R1013) was then added, and columns were centrifuged (16,100 x g, 30 s, room temp). Columns were washed again as before with 700 μL RNA Wash Buffer and then with 400 μL RNA Wash Buffer. The flowthrough was discarded and columns spun again for 2 minutes to remove all traces of wash buffer. Columns were transferred to a clean 1.7-mL tube, 15 μL RNase-free water was added to each, and after incubating at room temp for 5 min, samples were centrifuged (16,100 x g, 1 min, room temp). Eluted RNA was stored at −80 °C for subsequent analysis by RT-qPCR (RIP-qPCR) or sequencing (RIP-seq).

Replicate RNA immunoprecipitation experiments used cell pellets that were crosslinked on different days, and the immunoprecipitation, RNA isolation, and RT-qPCR steps were repeated on different days. The exception to this was the SPEN RIP-qPCR experiment in Figure 5F, in which two independent clonal lines of *FKBP1A^F36V^*-*Srsf1* cells were crosslinked and analyzed on the same days.

### Reverse transcription and quantitative PCR analysis of RIPs (RIP-qPCR)

Reverse transcription was performed with the Applied Biosystems High-Capacity cDNA Reverse Transcription Kit (Thermo Fisher 4368814) according to manufacturer instructions: 20-μL randomer-primed reverse transcription reactions were prepared in 0.2-mL PCR strip-tubes on ice, each containing 0.5 μL 40 U/μL RNaseOUT RNase Inhibitor (Thermo Fisher 10777019) and 5 μL of column-purified RNAs from 5% input or 100% RIP samples. Reverse transcription was performed with the following thermal cycler program: 25 °C for 10 min, 37 °C for 120 min, 85 °C for 5 min, hold at 4 °C. All reverse transcription products were first diluted 2-fold by thoroughly mixing with 20 μL nuclease-free water. For robust quantification via qPCR, standard curves were constructed by preparing 4-fold serial dilutions from each 5% input sample (10 μL previous standard mixed with 30 μL nuclease-free water), generating a total of 6 standards per standard curve.

To set up qPCR, a master mix was generated that contained, for each 10-μL qPCR reaction: 5 μL iTaq Universal SYBR Green Supermix (Bio-Rad 1725124), 3 μL nuclease-free water, 0.5 μL 10 μM forward primer, and 0.5 μL 10 μM reverse primer (see Table S6 for primer sequences). In Hard-Shell 96-Well PCR Plates (Bio-Rad HSP9601), qPCR reactions were prepared in technical triplicate by combining 9 μL qPCR master mix and 1 μL of standard-curve or RIP reverse transcription product. Standard-curve qPCR reactions from each 5% input and qPCR reactions for each 5% input’s corresponding RIP(s) were always run on the same plate. Plates were firmly sealed with Microseal ‘B’ PCR Plate Sealing Film (Bio-Rad MSB1001), mixed several times by inversion and low intensity vortexing, and briefly pulsed down in a centrifuge (up to 3000 x g) to collect the volumes and remove bubbles. Reactions were analyzed on a Bio-Rad C1000 Touch Thermal Cycler equipped with a CFX96 Real-Time System with the following standard program: 95 °C for 10 min, 40 cycles of (95 °C for 15 s, 60 °C for 30 s [annealing], 72 °C for 30 s, plateread), followed by melt curve analysis: 95 °C for 10 s, 65 °C for 31 s, 61 cycles of (65 °C for 5 s +0.5 °C/cycle Ramp 0.5 °C/s, plateread). A variant of this standard program in which the annealing step was set to 56 °C was used for Repeat A-targeting qPCR primers to accommodate their lower annealing temperatures (see Table S6). All qPCR primer pairs were confirmed via agarose gel electrophoresis to specifically amplify a single product of the expected size. Data were analyzed using Bio-Rad CFX Manager software (v3.1), first setting “Cq Determination Mode” to “Regression” and then using each standard curve (Cq vs. % input, with the average value from the most concentrated standard set to 5% input) to convert Cq values to “% input” values for each technical replicate of each RIP sample. Each qPCR technical triplicate value was plotted as a dot in GraphPad Prism (v9.5.0), with the average values being represented by bars. Significance was determined via two-tailed t-tests comparing the triplicate-averaged values from independent biological replicates.

### RIP-seq library preparation and sequencing

Single-read, strand-specific (reverse-stranded) libraries for RNA sequencing were prepared using the KAPA RNA HyperPrep Kit with RiboErase (HMR) (Roche 08098140702) and KAPA Unique Dual-Indexed Adapters (Roche 08861919702) or TruSeq adapters, with each library’s input consisting of 9 μL column-purified RNA from RIP and 1 μL of a 1:250 dilution of ERCC RNA spike-in controls (Thermo Fisher 4456740). RNA fragmentation was performed at 85 °C for 5 min, diluted adapter stocks were 1.5 μM, and libraries were amplified using 13 to 21 cycles of PCR depending on the sample, with the number of cycles determined by qPCR to compare against positive-control pre-amplification libraries that had been previously amplified, quantified, and sequenced. Libraries were quantified using a Qubit 2.0 fluorometer (dsDNA broad sensitivity kit, Thermo Fisher Q32850), and average fragment sizes were determined by agarose gel electrophoresis. Libraries were pooled and sequenced on an Illumina NextSeq500 platform with a 75-cycle NextSeq 500/550 High Output Kit v2.5 (Illumina 20024906) or Illumina NextSeq1000 platform with a 100-cycle NextSeq 1000/2000 P2 Reagent Kit v3 (Illumina 20046811) or 100-cycle NextSeq 1000/2000 P2 XLEAP-SBS Reagent Kit (Illumina 20100987).

### RIP-seq wiggle track generation

RIP-seq reads were aligned to mm10 using STAR (v2.7.9a, ^141^), using single-pass mapping and the option “--outFilterMultimapNmax 1” to consider only uniquely mapping reads. Output SAM files from each alignment were parsed by strand using Samtools (v1.15 or v1.20): reads from Samtools view -F 0x10 were retained as negative-stranded reads, and reads from Samtools view -f 0x10 were retained as positive-stranded reads. For each strand-specific SAM file, wiggle files were generated using the custom Perl script bigsam_to_wig_mm10_wcigar2.pl, which binned reads into 50-nt bins. Wiggle files were uploaded to the UCSC Genome Browser to visualize data. For viewing reads over *Xist*, browser tracks were configured such that “Data view” was set to “auto-scale to data view” and “Windowing function” was set to “mean.” Views over *Xist* (which is on the mm10 negative strand) were exported in PDF file format and flipped horizontally in Adobe Illustrator to display data from 5′ to 3′. Track heights were RPM-normalized by dividing the auto-scaled heights (reads per 50-nt bin) by the total number of reads (in millions) uniquely mapped by STAR for that sample.

To quantify reads over Repeat A or the entirety of *Xist*, featureCounts (Subread v2.0.4, ^143^) was used. For Repeat A reads, a SAF file was generated with a region defined as chrX:103482476-103482926, and featureCounts was run with options “-s 2 -F SAF -a Repeat_A.saf.” For *Xist* reads, featureCounts was run with options “-t exon -g gene_id -a mm10_genes.gtf,” and *Xist* reads were collected with the Unix command grep ‘Xist’.

### RIP-seq peak calling

SPEN and Halo RIP-seq FASTQ files were aligned with STAR (2.7.10a, ^141^) to a STAR genomic index built from the primary assembly of the GRCm38 (mm10) genome with reference to the GENCODE vM25 basic annotation GTF. Additional flags of “--outMultimapperOrder Random” and “--outSAMmultNmax 1” were used. Alignments were then filtered to remove multimapping reads (MAPQ scores less than 30) with Samtools (v1.21) view -q 30. For all RIP-seq experiments, the filtered alignments were split by strand, reads from Samtools view -F 0x10 were retained as negative-stranded reads, and reads from Samtools view -f 0x10 were retained as positive-stranded reads. The strand information was then randomly re-assigned through a custom Perl script (macs_strand_rand_sam.pl),^135^ and these strand-randomized alignments were converted to BAM files and provided as input to the MACS2 (v2.2.7.1; ^148^) callpeak function with the flags “--keep-dup all”, “--broad”, “--broad-cutoff 0.3”, and “--max-gap 76”. To identify the number of reads under each peak, the MACS2 peaks for each RIP-seq experiment were converted from broadPeak to SAF file formats and then provided as input to featureCounts (Subread v2.0.6, ^143^) with the additional flags “-F SAF and -s 2,” as the data is reversely stranded. Additionally, the same peaks were also given to featureCounts with the appropriate control sample (SM33 Δ*Spen* or RMCE-*Xist* Δ*Spen* no-rescue) alignment. The number of reads under each peak was converted to RPM and the peak set was filtered to retain only peaks where the RPM of each replicate was greater than two times the RPM value of the control in that same peak region.

### RIP-seq motif analysis

To identify motif enrichment that excluded RIP signal from *Spen* and *Xist* transcripts, the coordinates for both genes were used to make a miniature custom GTF file that was used in BEDTools intersect (v2.29, ^149^) as -b input. The -v flag was also used to exclude intersecting peaks, resulting in a new final set of peaks in BED file format. Next, scaled signal over control for each peak region was calculated by squaring the RIP RPM value and dividing that by the sum of the RIP RPM and the control-sample RIP RPM. Peaks were then sorted by decreasing numerical scaled signal values, and the top 1500 peaks were retained. The sequences for the top 1500 ranked peaks were extracted from the GRCm38 primary assembly genome FASTA by BEDTools getfasta command with the -s flag. A paired set of 1500 length-matched control sequences was generated for the top 1500 peak sequences of each RIP experiment by calculating the nucleotide probabilities from the nucleotide content of the genome, which weighted random nucleotide selection in random sequence creation. Finally, enriched motifs were established with STREME (MEME Suite v5.2.0, ^150^) for the top 1500 peak sequences for each RIP experiment individually, with the corresponding length-matched control sequences provided with the -n flag, a maximum window of --maxw 12, minimum window of --minw 4, --nmotifs 20, and the --rna and -p flags. To determine motifs enriched within the 189 SPEN RRM1-4 RIP peaks in the 134 genes upregulated upon *Spen* deletion, the same protocol was repeated except that STREME was run using MEME Suite v5.5.7 against a set of 1890 length-matched control sequences.

### Protein purification

To 400 mL pre-warmed lysogeny broth (LB; Research Products International L24041) containing 100 μg/mL ampicillin sodium salt (Fisher Scientific BP1760-25) and 34 μg/mL chloramphenicol (Acros Organics 227920250) in a 1-L flask, 4 mL of overnight culture of Rosetta *Escherichia coli* cells (MilliporeSigma 70953) transformed with pDD001-SPEN-RRM2-4 or pDD001-SRSF1 RRM1-2 was added. Cultures were grown at 37 °C with 180 rpm shaking until OD_600_ reached 0.4-0.6, after which cultures were cooled to 17 °C, and then isopropyl β-D-1-thiogalactopyranoside (IPTG; Sigma I6758) was added to a final concentration of 0.5 mM. After ∼16-h induction at 17 °C with 180 rpm shaking, cells were pelleted by centrifugation (5000 x g, 10 min, 4 °C) and gently resuspended by swirling in 20 mL GST Lysis Buffer (20 mM Tris-HCl pH 7.5, 20 mM NaCl, 1% [v/v] Triton X-100; containing freshly added SigmaFast EDTA-free protease inhibitor cocktail [1 tablet per 100 mL]) with freshly added 2 mM DTT and 2 mM phenylmethylsulfonyl fluoride (PMSF). Resuspended cells were lysed on ice, with 6 30-s pulses of a Sonics Vibra-Cell 130-watt VCX 130 probe-tip sonicator (50% amplitude output) and a 30-s rest on ice between each pulse. Lysates were cleared by centrifugation (36635 x g, 30 min, 4 °C) and then added to 1 mL slurry volume of Pierce Glutathione Agarose Beads (Thermo Fisher 16100) pre-equilibrated in GST Lysis Buffer in a 50-mL conical tube on ice. Lysates were incubated with beads for 30 min at 4 °C with end-over-end rotation and then gently pelleted by centrifugation (500 x g, 1 min, room temp). Unbound lysate supernatants were carefully removed from the beads, and then the beads were washed 5 min per wash at 4 °C with end-over-end rotation and pelleted as before, using 10 mL of the following buffers per wash: twice with GST Wash Buffer A (20 mM Tris-HCl pH 7.5, 300 mM NaCl), twice with GST Wash Buffer B (20 mM Tris-HCl pH 7.5, 1 M NaCl), and then twice again with GST Wash Buffer A. All but ∼500 μL the final wash buffer was removed from the beads, the beads were resuspended in the remaining buffer and transferred to a 1.7-mL tube, and the remaining buffer was carefully removed after pelleting by centrifugation (1000 x g, 2 min, 4 °C). To the beads, 500 μL Cleavage Buffer (20 mM Tris-HCl pH 7.5, 100 mM NaCl, 0.01% [v/v] Triton X-100, 10% [v/v] glycerol, 5 mM DTT [freshly added]) was added, followed by 10 μL PreScission Protease (Cytiva 27-0843-01). SBP-tagged proteins were cleaved from the bead-bound GST by incubation at 4 °C with end-over-end rotation for ∼18 h.

Beads were pelleted by centrifugation (2000 x g, 2 min, 4 °C), and supernatants were carefully collected and transferred to a new pre-chilled tube on ice. To clear residual beads from the supernatants, tubes were centrifuged again (16100 x g, 5 min, 4 °C), and supernatants (final protein preparations) were transferred to pre-chilled tubes on ice, aliquoted to small volumes, snap-frozen with liquid nitrogen, and stored at −80 °C. To assess purity and measure concentration, each protein preparation was run alongside a standard curve of known amounts of BSA (Fisher Scientific BP1600) on a pre-cast Bio-Rad TGX SDS-PAGE gel in Tris-glycine-SDS running buffer (2.5 mM Tris, 19.2 mM glycine, 0.01% [w/v] SDS, pH 8.3) at 150 V. The gel was stained with GelCode Blue (Thermo Fisher 24590), destained with water, and imaged with a Bio-Rad ChemiDoc MP imager (Coomassie Blue setting). Densitometry analysis was performed with Bio-Rad Image Lab v6.1 software to determine band intensities of SBP-tagged proteins and the sums of band intensities in each BSA standard; using Microsoft Excel software, SBP-tagged protein concentrations were calculated by fitting to a best-fit line of the BSA standards.

### In vitro transcription (IVT) of biotinylated and unlabeled RNA

Double-stranded DNA templates for IVT were generated by PCR using Q5 High-Fidelity DNA Polymerase (NEB M0491), pTETRISv1-*Xist*-2kb wild-type/mutant as template, and primers specific to *Xist* that add the SP6 promoter sequence upstream of Repeat A (Table S6). DNA templates were run on an agarose-TAE gel containing ethidium bromide, and excised DNA bands were purified using the GenCatch Gel Extraction Kit (Epoch Life Science 2260250). DNA concentration was measured with a NanoDrop Lite.

To generate internally biotinylated RNA, IVT was performed overnight (∼16-24 h) at 37 °C using the MEGAscript SP6 Transcription Kit (Thermo Fisher AM1330) following manufacturer instructions, with the addition of biotinylated UTP (MilliporeSigma 11388908910 or TriLink Biotechnologies N-5005) to a final concentration of 0.5 mM (such that the molar ratio of biotinylated UTP to UTP was 1:10) and 0.5 pmol DNA template per 20 μL of reaction. To generate large-scale amounts of RNA, reaction volumes were frequently scaled up to 100 μL. IVT products were treated with TURBO DNase (0.8 μL per 20 μL reaction, 15 min at 37 °C) and purified via standard TRIzol-chloroform extraction, per manufacturer instructions (Thermo Fisher 15596026). RNA concentration was measured with a NanoDrop Lite, and purity was assessed by urea-PAGE with Promega Diamond (Promega H1181) staining. RNA was stored in small aliquots at −80 °C.

To generate unlabeled RNA for RBNQ, the same procedure was performed with the following differences: biotinylated UTP was omitted, RNA was purified using the MEGAclear Transcription Clean-Up Kit (Thermo Fisher AM1908) following manufacturer instructions except that elution was performed at 37 °C to preserve native RNA folding, and RNA concentration was measured with a Qubit 2.0 fluorometer (RNA high-sensitivity or broad-range kit, Thermo Fisher Q32852 and Q10211).

### Equilibrium RNA bind-n-qPCR (RBNQ)

The following protocol is an adaptation of one published previously ^55^, modified to enable equilibrium assessment of binding and accurate comparison to input. SBP-tagged proteins were first diluted into room-temp RBNQ Binding Buffer with Random RNA (3 mM MgCl_2_, 25 mM Tris-HCl pH 8.05, 150 mM KCl, 1 mg/mL BSA [Thermo Fisher AM2616], 2 U/mL SUPERase-In [Thermo Fisher Scientific AM2694], 50 nM random 16-nt RNA [IDT, bases randomized by machine-mixing, purified by standard desalting]) to a range of different protein concentrations (∼1-1400 nM). Using a room-temp magnetic separator for buffer exchanges, a volume of 250 μL well-mixed Dynabeads MyOne Streptavidin T1 (Thermo Fisher 65602) was washed once with 250 μL RNase-free water, twice with 250 μL RBNQ Blocking Buffer (3 mM MgCl_2_, 25 mM Tris-HCl pH 8.05, 150 mM KCl, 1 mg/mL BSA [Thermo Fisher AM2616], 2 U/mL SUPERase-In [Thermo Fisher AM2694], 1 mg/mL yeast tRNA [Thermo Fisher AM7119]), and twice with 250 μL RBNQ Binding Buffer with Random RNA. Beads were then resuspended in 125 μL RBNQ Binding Buffer with Random RNA, and 10 μL of the well-mixed slurry was aliquoted to each of 12 1.7-mL tubes. (Volumes of beads and washes were scaled proportionately for experiments with more than 12 binding reactions.) To each tube, 30 μL of SBP-tagged protein (at various concentrations) was added and gently mixed by pipet, and tubes were incubated for 10 min (in a time-staggered manner from this point forward to ensure consistent handling) at room temp with end-over-end rotation for proteins to bind to the beads. Tubes were placed on the magnetic separator; supernatants were removed; beads were carefully resuspended in 30 μL of 0.1 nM in-vitro-transcribed, unlabeled Repeat A RNA in RBNQ Binding Buffer with Random RNA; and tubes were incubated for 30 min at room temp with end-over-end rotation to enable RNA-protein binding reactions to equilibrate. Meanwhile, 30 μL of each 0.1 nM in-vitro-transcribed, unlabeled Repeat A RNA was transferred to prechilled tubes on ice for “input RNA” samples. Keeping uniform, time-staggered treatment, tubes were placed on the magnetic separator, unbound supernatants were carefully and completely removed, beads were resuspended in 100 μL of Elution Buffer (25 mM Tris-HCl pH 8.05, 4 mM D-biotin [Thermo Fisher B1595], 8% [v/v] dimethyl sulfoxide [DMSO]), and tubes were incubated for 30 min at 37 °C. Tubes were placed on the magnetic separator, and “eluate RNA” supernatants were transferred to pre-chilled tubes on ice. “Input RNA” samples were brought up to 100 μL by adding 70 μL Elution Buffer and mixing well.

For RBNQ assays examining differences in protein binding to wild-type versus mutant Repeat A RNA, the same protocol was followed except that the volumes of Dynabeads and bead washes were cut in half (i.e., 10 μL Dynabead slurry was used per binding reaction and washes were with 125 μL) and that a single concentration of protein was used (near the pre-determined K_d_ for wild-type RNA). Replicate RBNQ experiments used protein preps that were purified on separate days; protein concentrations of the independent preps were measured at the same time to ensure accuracy and precision of RBNQ data.

Reverse transcription was performed with the Applied Biosystems High-Capacity cDNA Reverse Transcription Kit (Thermo Fisher 4368814) according to manufacturer instructions: 20-μL randomer-primed reverse transcription reactions were prepared in 0.2-mL PCR strip-tubes on ice, each containing 0.5 μL 40 U/μL RNaseOUT RNase Inhibitor (Thermo Fisher 10777019) and 5 μL of input or eluate RNA. Reverse transcription was performed with the following thermal cycler program: 25 °C for 10 min, 37 °C for 120 min, 85 °C for 5 min, hold at 4 °C. All reverse transcription products were first diluted 2-fold by thoroughly mixing with 20 μL nuclease-free water. For robust quantification of binding-curve RBNQ experiments via qPCR, standard curves were constructed by preparing 2-fold serial dilutions from each input sample (20 μL previous standard mixed with 20 μL nuclease-free water), generating a total of 8 standards per standard curve. For fixed-concentration RBNQ comparing WT and mutant Repeat A, 6-standard standard curves were constructed via 2.5-fold serial dilutions from each input sample.

To set up qPCR, a master mix was generated that contained, for each 10-μL qPCR reaction: 5 μL iTaq Universal SYBR Green Supermix (Bio-Rad 1725124), 3 μL nuclease-free water, 0.5 μL 10 μM forward primer, and 0.5 μL 10 μM reverse primer (see Table S6 for oligonucleotide sequences). In Hard-Shell 96-Well PCR Plates (Bio-Rad HSP9601), qPCR reactions were prepared in technical triplicate (technical duplicate for standard-curve samples in fixed-concentration WT vs. mutant Repeat A RBNQ experiments) by combining 9 μL qPCR master mix and 1 μL of input or eluate reverse transcription product. Standard-curve qPCR reactions from each input and corresponding eluate(s) were always run on the same plate. Plates were firmly sealed with Microseal ’B’ PCR Plate Sealing Film (Bio-Rad MSB1001), mixed several times by inversion and low intensity vortexing, and briefly pulsed down in a centrifuge (up to 3000 x g) to collect the volumes and remove bubbles. Reactions were analyzed on a Bio-Rad C1000 Touch Thermal Cycler equipped with a CFX96 Real-Time System with the following standard program: 95 °C for 10 min, 40 cycles of (95 °C for 15 s, 56 °C for 30 s, 72 °C for 30 s, plateread), followed by melt curve analysis: 95 °C for 10 s, 65 °C for 31 s, 61 cycles of (65 °C for 5 s +0.5 °C/cycle Ramp 0.5 °C/s, plateread). All qPCR primer pairs were confirmed via agarose gel electrophoresis to specifically amplify a single product of the expected size. Data were analyzed using Bio-Rad CFX Manager software (v3.1), first setting “Cq Determination Mode” to “Regression” and then using each standard curve (Cq vs. % input, with the average value from the most concentrated standard set to 100% input) to convert Cq values to “% input” values for each technical replicate of each eluate sample. For binding-curve RBNQ experiments, input-normalized values from each of two experimental replicates were plotted together in GraphPad Prism (v9.5.0), with the average values and standard deviation of qPCR technical triplicates being represented by dots and error bars, respectively. The equilibrium dissociation constant K_d_ and Hill slope h were determined in GraphPad Prism by fitting to the equation Y=B_max_*X^h^/(K_d_^h^ + X^h^) with Analyze > XY analyses > nonlinear regression (curve fit) > Binding - Saturation > Specific binding with Hill Slope. For fixed-concentration WT vs. mutant Repeat A RBNQ experiments, each qPCR technical triplicate value was plotted as a dot in GraphPad Prism (v9.5.0), with the average values being represented by bars. Significance was determined via two-tailed t-tests comparing the triplicate-averaged values from independent biological replicates.

### Preparation of nuclear extracts for biotinylated RNA pulldown

Between 14 and 20 75-85% confluent 15-cm dishes of E14 mESCs were each washed three times with 10 mL ice-cold PBS before harvesting by scraping into 1.5 mL ice-cold PBS. Cells were transferred to pre-chilled conical tubes and pelleted by centrifugation (1136 x g, 5 min, 4 °C), supernatants were removed, and cells were gently resuspended by serological pipet collectively in 7 mL ice-cold PBS and consolidated in one 15-mL conical tube before pelleting by centrifugation (1136 x g, 5 min, 4 °C). The volume of the cell pellet was estimated (∼1.2-1.5 mL), and cells were resuspended by serological pipet in 2 volumes of ice-cold NE Buffer A (10 mM HEPES-KOH pH 7.9, 1.5 mM MgCl_2_, 10 mM KCl, with the following added fresh: 0.5 mM DTT, 0.5 mM PMSF, 1:200 protease inhibitor cocktail [Sigma P8340]). IGEPAL CA-630 was added to a final concentration of 0.1% (v/v), and the cell suspension was gently inverted to mix before incubating on ice for 5 min. The cell suspension was placed in a pre-chilled Dounce homogenizer and homogenized with 10 up-down strokes of a tight-fitting pestle, taking care avoid introducing bubbles. The mixture was pelleted by centrifugation (2148 x g, 4 min, 4 °C), and the supernatant (cytoplasmic extract) was removed. To prepare nuclear extract, the pellet of nuclei was resuspended by serological pipet in 1.5 original-cell-pellet-volumes of ice-cold NE Buffer C (20 mM HEPES-KOH pH 7.9, 25% [v/v] glycerol, 420 mM KCl, 1.5 mM MgCl_2_, 0.2 mM EDTA-NaOH pH 8.0, with the following added fresh: 0.5 mM DTT, 0.5 mM PMSF, 1:200 protease inhibitor cocktail [Sigma P8340]) and incubated end-over-end at 4 °C for 30 min. The mixture was pelleted by centrifugation (5000 x g, 5 min, 4 °C), and supernatant (nuclear extract) was transferred to a pre-chilled tube on ice. From the pellet of nuclei and nuclear debris, the extraction process was repeated with another 1.5 original-cell-pellet-volumes of ice-cold NE Buffer C. The nuclear extract samples were cleared by centrifugation (16,100 x g, 15 min, 4 °C), and the supernatants were pooled together to a volume of ∼3-6 mL. Using Slide-A-Lyzer 10-kDa MWCO devices (Thermo Fisher 88404), cleared nuclear extracts were dialyzed twice at 4 °C into NE Buffer D (20 mM HEPES-KOH pH 7.9, 20% [v/v] glycerol, 100 mM KCl, 0.2 mM EDTA-NaOH pH 8.0, with 0.5 mM DTT added fresh); one dialysis step lasted 2-3 hours and the other lasted ∼18 hours. To remove precipitated insoluble material, dialyzed nuclear extracts were pelleted by centrifugation (16,100 x g, 15 min, 4 °C), and supernatants (final nuclear extracts) were pooled, measured by Bradford assay (Bio-Rad 5000006) against a standard curve of BSA to determine protein concentration (∼2.5-3 μg/μL), and snap-frozen in liquid nitrogen before storing at −80 °C. Aliquots were shipped on dry ice for proteomics analysis of nuclear extracts (biotinylated RNA pulldown inputs).

### Biotinylated RNA pulldown

The following protocol is a modification of one published previously ^3^, modified to increase the amount of RNA and protein, to clarify crosslinking conditions, and to detail the elution of proteins with RNase A. In a pre-chilled tube on ice for each pulldown sample, a volume of nuclear extract containing 1100 μg protein was brought to 500 μL with NE Buffer D, after which 1 mL ice cold Incubation Buffer (150 mM NaCl, 15 mM Tris-HCl pH 7.5, 0.5 mM EDTA-NaOH pH 8.0, 0.025% [v/v] IGEPAL CA-630, with the following added fresh: 1 mM DTT, 1:100 protease inhibitor cocktail [Sigma P8340], 100 U/mL SUPERase-In RNase inhibitor [Thermo Fisher AM2696]) and 11 μg internally biotinylated, in-vitro-transcribed RNA were added. Tubes were incubated end-over-end at 4 °C overnight.

The next day, for each pulldown, 25 μL slurry volume of Dynabeads M-280 streptavidin beads (Thermo Fisher 11205D) was washed twice with 500 μL ice-cold Incubation Buffer (without DTT, protease inhibitor cocktail, and SUPERase-In) with aid of a magnetic separator, keeping the beads in the second wash on ice until ready for addition of crosslinked RNA-protein mixtures. The RNA-protein mixtures were transferred to individual wells of a pre-chilled 6-well cell culture plate (Genesee Scientific 25-105) on ice. To induce crosslinks with UV light, the plate on ice (with lid removed) was placed in a Stratagene UV Stratalinker 1800 fitted with 254-nm bulbs and irradiated with the auto-crosslink setting (2500 x 100 μJ/cm^2^; value of 2500 entered on the digital screen). Washed beads were resuspended in crosslinked RNA-protein mixtures and incubated end-over-end at 4 °C overnight.

The next day, the beads were washed (5 min per wash, end-over-end at 4 °C) with 500 μL of the following buffers per wash: twice with ice-cold PD Wash Buffer A (150 mM NaCl, 15 mM Tris-HCl pH 7.5, 0.5 mM EDTA-NaOH pH 8.0, 1 mM DTT, 1:100 protease inhibitor cocktail [Sigma P8340]), twice with room-temp (to prevent SDS precipitation) PD Wash Buffer B (450 mM NaCl, 15 mM Tris-HCl pH 7.5, 0.5 mM EDTA-NaOH pH 8.0, 0.025% [v/v] IGEPAL CA-630, 0.05% [w/v] SDS, 0.1 mM DTT, 1:100 protease inhibitor cocktail [Sigma P8340]), and twice with ice-cold PD Wash Buffer C (150 mM NaCl, 15 mM Tris-HCl pH 7.5, 0.5 mM EDTA-NaOH pH 8.0, 1:100 protease inhibitor cocktail [Sigma P8340]). Mixtures were transferred to new pre-chilled tubes on ice before each of the final two washes (resuspending the beads in PD Wash Buffer C before transferring). After complete removal of the final wash, beads were resuspended in 20 μL of PD Wash Buffer C containing 0.005 μg/μL RNase A (Thermo Fisher EN0531, a 1:20,000 dilution). Tubes were incubated at 37 °C for 30 min, gently resuspending every 5 min to keep beads well suspended. Supernatants (first eluates) were transferred to pre-chilled tubes on ice. To collect residual proteins, beads were again resuspended in 15 μL of PD Wash Buffer C, placed on the magnetic separator, and supernatants were added to the first eluates and mixed for a total eluate volume of ∼35 μL. A volume of 5 μL of each eluate was analyzed by SDS-PAGE on a 4-20% polyacrylamide gel (Bio-Rad 4561094) and silver staining with the Pierce Silver Stain Kit (Thermo Fisher 24612). The remaining eluates were snap-frozen in liquid nitrogen and stored at −80 °C prior to shipping on dry ice for proteomics analysis.

### Repeat A proteomics sample analysis

Nuclear extract (input) samples: For in-solution Suspension Trap (S-Trap) digestion, 11.5 μL (∼2 μg/μL) of each nuclear extract sample (inputs for RNA pulldowns) was mixed with in 11.5 μL of 10% SDS solution for 23 μL of initial-volume S-Trap protocol (ProtiFi K02-micro-10). After overnight trypsin digestion, peptides were reconstituted with 25 μL of 5% acetonitrile and 0.1% formic acid, and absorbances were measured. Due to higher absorbance, they were further diluted 2 times, and 0.6 pmole of yeast alcohol dehydrogenase (ADH) was spiked in for quantitative sample-to-sample comparison, retention time reproducibility, and mass accuracy throughout the runs. A 3.8-μL portion of each sample (independent replicate input samples, each analyzed three times) was injected into a to Waters nanoACQUITY UPLC coupled to a Thermo Scientific Orbitrap Fusion Lumos Tribrid mass spectrometer. The proteins were searched against the SwissProt Mouse database (V9) in Mascot Search Engine in Proteome Discoverer v2.1, followed by export to Scaffold v4 for false-discovery-rate (FDR) and intensity-based absolute quantitation (iBAQ) analysis.

RNA pulldown samples: In-solution S-Trap digestion was performed with 30-μL initial volume for each sample. After digesting with trypsin, samples were reconstituted with 20 μL 5% acetonitrile, 0.1% (v/v) formic acid in water following 0.2 pmole ADH spike-in. A 3.8 μL portion of each sample (independent replicate RNA pulldown samples, each analyzed once) was then injected to a Waters nanoAcquity UPLC coupled to a Thermo Scientific Orbitrap Fusion Lumos Tribrid mass spectrometer. After data acquisition, raw files were loaded to Proteome Discoverer software, and database SwissProt Mouse was used for the Mascot Search Engine followed by Scaffold analysis. The Scaffold analysis was carried out at 1% protein FDR.

### Repeat A proteomics data analysis

Nuclear extract (input) samples: iBAQ data were exported from Scaffold to Microsoft Excel for processing. All iBAQ values of 0 or −1 were considered missing values and converted to values of 0. Proteins deemed as sample contaminants (yeast ADH, all keratins, and spurious entry “744”) were removed, and proteins with multiple entries (Hnrnpa3 [2 entries], Hnrnpk [2 entries], Tmpo [2 entries], Hnrnpc [3 entries], Cpsf4 [2 entries]) were combined into one entry, summing their iBAQ values within each sample replicate. Within each sample replicate, all protein iBAQ values were normalized to total signal by dividing each protein’s iBAQ value by the sum of all iBAQ values within the sample replicate, then multiplying by an arbitrary scaling factor of 1 million. To ensure robust estimation of relative protein abundance in the nuclear extracts, 149 proteins were removed for which 5 or more of the 9 normalized iBAQ values were missing. For the 2826 proteins remaining, nonmissing (nonzero) normalized iBAQ values were averaged, and the base-2 logarithm of this average was calculated to compare values to those from biotinylated RNA pulldowns on the same logarithmic scale (below).

RNA pulldown samples: iBAQ data were exported from Scaffold to Microsoft Excel and processed using Microsoft Excel and Python. All iBAQ values of 0 or −1 were considered missing values and converted to values of 0. Proteins deemed as sample contaminants were removed: yeast ADH, all keratins, and the 8 proteins with nonmissing iBAQ values in the “no RNA” control. Proteins with multiple entries (Hnrnpa3 [2 entries], Hnrnpc [2 entries]) were combined into one entry, summing their iBAQ values within each sample replicate. Within each sample replicate, all protein iBAQ values were normalized to total signal by dividing each protein’s iBAQ value by the sum of all iBAQ values within the sample replicate, then multiplying by an arbitrary scaling factor of 1 million. (Due to intra-group variability of total-sample iBAQ values [e.g., within the WT samples], normalization to total-sample iBAQ signal was deemed more appropriate than normalization to the yeast ADH spike-in. One of four WT replicates and one of three ΔCCUGC replicates were removed from the analysis due to substantially poorer total protein detection.) To consider only proteins recovered reproducibly in the RNA pulldowns, the 844 proteins with iBAQ signal in at least one sample were filtered to a list of 644 proteins that fit at least one of two criteria: (1) at least 2 of the 3 WT values were nonmissing [614 proteins], or (2) all of the values for at least one mutant group were nonmissing [30 additional proteins]. Each nonmissing value was replaced with its base-2 logarithm to transform the data to a roughly normal distribution, and missing values were replaced according to the nature of their missingness: when only 1 value of a sample group was missing, it was assumed to be missing-at-random and was replaced with a not-a-number (NaN) to omit it from averages and statistical tests; when 2 or 3 values of a sample group were missing, these were assumed to be missing-not-at-random and underwent “Perseus-style” imputation, replacing them with values generated at random from a normal distribution centered at (iBAQ_log2_mean) – 2*(iBAQ_log2_sd) with a standard deviation of 0.3*[(iBAQ_log2_mean) – 2*(iBAQ_log2_sd)], where iBAQ_log2_mean and iBAQ_log2_sd were the average and standard deviation, respectively, of all nonmissing values (total-normalized and log2-converted) in the dataset.

Two-sample t-tests were performed comparing the log2-transformed replicate values for WT and ΔGC-core pulldowns from three independent experiments, and the proteins recovered more strongly with WT and with *p* < 0.05 were considered the set of GC-rich core-interacting proteins highlighted in Figure 3B.

The heatmap in Figure 3D was generated by first filtering rows of proteins for those classified as SR proteins ^57^ and then using Morpheus (https://software.broadinstitute.org/morpheus), keeping column order fixed but enabling hierarchical clustering of rows with “metric” set to “one minus Pearson correlation” and “linkage method” set to “average.”

### Gene ontology analysis

Gene ontology of gene or protein sets were analyzed with the Molecular Signatures Database (MSigDB) at https://www.gsea-msigdb.org/gsea/msigdb ^151,152^, using mouse collections for all sets except for the sets of *SPEN*-correlated genes from GTEx and TCGA, which used the human collections. The top overlapping gene sets in GO biological process (GO:BP) and GO cellular component (GO:CC) were determined and reported with their enrichment p-values. Gene/protein classifications were defined by the following sources: SR proteins ^57^, nuclear speckles (MSigDB GO term GOCC_NUCLEAR_SPECK.v2023.2.Mm for mouse and MSigDB GO term GOCC_NUCLEAR_SPECK.v2024.1.Hs for human), paraspeckles ^153^, and nucleolus (MSigDB GO term GOCC_NUCLEOUS.v2023.2.Mm).

### Halo-SPEN immunoprecipitation for proteomics analysis

Immunoprecipitation for proteomics analysis generally followed the RIP protocol above but with modifications made to accommodate larger inputs and a different method to reverse formaldehyde-induced crosslinks.

For each immunoprecipitation, well-mixed slurry volume of 50 μL Protein A/G PLUS-Agarose beads (Santa Cruz sc-2003) was washed three times with 1 mL room-temp 0.5% (w/v) bovine serum albumin (BSA; Fisher Scientific BP1600) in PBS. For each wash, 1.7-mL tubes were gently inverted three times and pelleted by centrifugation (2000 x g, 1 min, room temp), and supernatants were removed carefully, leaving ∼10-20 μL to ensure no loss of beads. To washed beads, 300 μL 0.5% (w/v) BSA in PBS was added, followed by 15 μL 1 μg/μL anti-Halo antibody (Promega G9281). Tubes were incubated overnight at 4 °C with end-over-end rotation.

The following day, formaldehyde-crosslinked 10-million-cell pellets (7 pellets for Halo-SPEN-V5 WT, ΔIDR2 ΔIDR4, ΔSPOC, and no-rescue control IPs; 3 pellets for Halo-SPEN-V5 RRM1-4 and RRM2-4 IPs; see “RNA Immunoprecipitation” section) were thawed on ice and resuspended with 10 gentle up-down strokes of a P1000 pipet in 500 μL ice-cold RIPA Buffer (50 mM Tris-HCl pH 8.0, 150 mM KCl, 5 mM EDTA-NaOH pH 8.0, 1% [v/v] Triton X-100 [Fisher Scientific BP151], 0.5% [w/v] sodium deoxycholate, 0.1% [w/v] sodium dodecylsulfate) containing the following components added immediately before use: 0.5 μL per mL 1 M dithiothreitol (DTT), 10 μL per mL protease inhibitor cocktail (Sigma P8340), and 5 μL per mL 20 U/μL SUPERase-In RNase inhibitor (Invitrogen AM2696). Keeping tubes on ice, each cell suspension was sonicated twice for 30 s with a Sonics Vibra-Cell 130-watt VCX 130 probe-tip sonicator (30% amplitude output), with a 1-min rest on ice between sonication pulses. Tubes were centrifuged (15000 x g, 15 min, 4 °C), and 505 μL of each supernatant (cleared cell lysate) was transferred to a new pre-chilled 15-mL conical tube, pooling lysates for each genotype.

The antibody-bound beads from the day before were washed twice with 1 mL ice-cold fRIP Buffer; for each wash, 1.7-mL tubes were gently inverted three times and pelleted by centrifugation (2000 x g, 1 min, 4 °C), and supernatants were carefully removed, leaving ∼10-20 μL to ensure no loss of beads. A 1-mL portion of complete fRIP Buffer (25 mM Tris-HCl pH 7.5, 150 mL KCl, 5 mM EDTA-NaOH pH 8.0, 0.5% [v/v] IGEPAL CA-630 [Fisher Scientific ICN19859650]; containing the following components added immediately before use: 0.5 μL per mL 1 M DTT, 10 μL per mL Sigma protease inhibitor cocktail, and 5 μL per mL 20 U/μL SUPERase-In) was used to completely collect washed antibody-bound beads and transfer to the 15-mL tubes containing lysates. A volume of fRIP Buffer was then added to each tube such that the total volume of complete fRIP Buffer was the same as the volume of pooled lysates in complete RIPA Buffer.

The next day, beads were pelleted by centrifugation (2000 x g, 1 min, 4 °C), and supernatants were carefully removed. The beads were washed once with 1 mL ice-cold fRIP Buffer, which was used to carefully transfer them to new 1.7-mL tubes. Beads were then washed three times with 1 mL ice-cold Pol II ChIP Buffer (50 mM Tris-HCl pH 7.5, 140 mM NaCl, 1 mM EDTA-NaOH pH 8.0, 1 mM EGTA-NaOH pH 8.0, 1% [v/v] Triton X-100, 0.1% [w/v] sodium deoxycholate, 0.1% [w/v] sodium dodecylsulfate), once with ice-cold 1 mL High-Salt Pol II ChIP Buffer (50 mM Tris-HCl pH 7.5, 500 mM NaCl, 1 mM EDTA-NaOH pH 8.0, 1 mM EGTA-NaOH pH 8.0, 1% [v/v] Triton X-100, 0.1% [w/v] sodium deoxycholate, 0.1% [w/v] sodium dodecylsulfate), and once with 1 mL LiCl Buffer (20 mM Tris pH 8.0, 1 mM EDTA-NaOH pH 8.0, 250 mM LiCl, 0.5% [v/v] IGEPAL CA- 630, 0.5% [w/v] sodium deoxycholate); each wash included a 5-min end-over-end rotation at 4 °C. In the final wash, samples were transferred to a new 1.7-mL tube. The final wash was carefully removed, first with a P1000 pipet and then with a P20 pipet, to ensure no loss of beads while leaving a volume of ∼5 μL LiCl Buffer on the beads. A volume of 50 μL 2% (w/v) sodium dodecylsulfate was forcefully added to the beads to suspend them, and samples were incubated at 95 °C for 40 min to reverse crosslinks. A volume of 200 μL water was added, samples were vortexed at maximum power for 10 s, and centrifuged to pellet beads (2000 x g, 1 min, room temp). Supernatants were transferred to pre-chilled tubes on ice, snap-frozen in liquid nitrogen, and stored at −80 °C prior to shipping on dry ice for proteomics analysis.

### Halo-SPEN IP proteomics sample analysis

Samples were flash frozen on dry ice and subsequently dried using a Speed-Vac. Next, 23 µL of 1x lysis buffer (5% SDS, 50 mM TEAB pH 8.5) was added to the protein pellet, initiating the Suspension trap (S-Trap) protocol (ProtiFi K02-micro-10). The proteins underwent reduction by 10 mM TCEP and alkylation by 20 mM iodoacetamide. Following an overnight digestion at 37 °C, 40 µL of 50 mM TEAB elution buffer was added and spun at 4000 rpm for 1 minute. This was followed by a 1-min spin with 40 µL of 0.2% formic acid (FA) and, lastly, 40 µL of 50% acetonitrile (ACN). The elution was then pooled together and dried using a Speed-Vac.

The digested samples were reconstituted with 20 µL of 0.1% formic acid in 5% ACN, followed by centrifugation at 16000 x g for 15 minutes. Subsequently, 18 µL of the sample was transferred to non-binding HPLC vials.

The samples (2 µL injection) were analyzed on a TimsTOF Pro2 (Bruker) mass spectrometer, which was coupled to a nanoElute LC system from Bruker. Peptides were loaded and separated on a house-made 75-μm I.D. fused silica analytical column packed with 25 cm ReproSil-Pur C18-AQ (Dr. Maisch, GmbH, 120 Å pore size, 3 μm particle size) particles to a gravity-pulled tip. A 30-min gradient was employed for DDA-PASEF with a flow rate set at 500 nL/min. The captive nano-electrospray voltage was maintained at 1600 V, using one column configuration (no trap) and a solvent composition of Solvent A: 0.1% FA in water and Solvent B: 0.1% FA in ACN. The analytical column was changed between the two batches of sample runs: the first batch consisted of the first biological replicates of no-rescue and Halo-SPEN-V5 WT IPs and only replicates of ΔIDR2 ΔIDR4 and ΔSPOC IPs, and the second batch contained all other IP samples (Table S2). Therefore, comparisons to no-rescue control were made within each batch.

DDA-PASEF method: The method consists of 10 MS/MS PASEF scans per topN acquisition cycle, with ramp and accumulation times of 100 ms, covering an m/z range from 100 to 1700 and an ion mobility range (1/K0) from 0.70 to 1.30 V s/cm^2^. Collision energy settings followed a linear function of ion mobility, ranging from 20 eV at 0.6 V s/cm^2^ to 59 eV at 1.6 V s/cm^2^, using default parameters. Calibration of the instrument was performed using three ions from the ESI-L Tuning Mix (Agilent) [m/z 622, 922, 1222].

### Halo-SPEN IP proteomics data analysis

The custom workflow ‘LFQ-MBR’ from FragPipe version 21.1 was used ^154^, including database search with MSFragger (version 4.0), deep-learning prediction rescoring with MSBooster, Percolator and ProteinProphet (Philosopher version 5.1.0) for PSM validation and protein inference. The raw .d files from Bruker were searched against *Mus musculus* reference proteome FATSA file UP000000589 from Uniprot (downloaded on 11/20/2023), appended with common contaminant proteins, containing in total 25483 sequence entries. Decoy reversed sequences were appended to the search database. The default MSFragger search parameters were used, except precursor and fragment mass tolerances were set to 20 ppm; enzyme: trypsin (not cutting before P); peptide length 7-30. Variable modifications were set to oxidation of Met and acetylation of protein N-t, and fixed modifications to carbamidomethylation of Cys. MSBooster, Percolator and ProteinProphet default options were used, and results were filtered by 1% FDR at protein level. The searched results were imported to Scaffold Q/S 5.2.2. Total spectrum counts were then exported to Microsoft Excel to calculate enrichments of proteins relative to no-rescue control (and between RRM1-4 and RRM2-4 samples) within each experimental replicate (Table S2). In Excel, all entries containing “keratin”, “albumin”, “Ig gamma”, “Ig kappa”, or “REVERSE” were considered contaminants and removed from the analysis. Data were visualized with GraphPad Prism (v9.5.0).

### Immunoprecipitations for analysis by western blot

Halo-SPEN-V5 WT and near-full-length mutant constructs expressed at low levels, and repeated attempts to visualize them (or endogenous SPEN) via western blot of lysate samples (up to 60-100 μg per lane) were unsuccessful. Therefore, an immunoprecipitation step was used prior to western blot analysis in order to enrich constructs from larger amounts of lysate (1000 μg) and to reduce nonspecific signal on the membrane.

For each immunoprecipitation, 25 μL well-mixed slurry volume of Protein A/G PLUS-Agarose beads (Santa Cruz sc-2003) was washed three times with 1 mL room-temp 0.5% (w/v) bovine serum albumin (BSA; Fisher Scientific BP1600) in PBS. For each wash, 1.7-mL tubes were gently inverted three times and pelleted by centrifugation (2000 x g, 1 min, room temp), and supernatants were removed carefully, leaving ∼10-20 μL to ensure no loss of beads. To washed beads, 300 μL 0.5% (w/v) BSA in PBS was added, followed by 5 μg anti-V5 antibody (Thermo Fisher R960-25). Tubes were incubated overnight at 4 °C with end-over-end rotation.

After 3 d of treatment with 1000 ng/mL doxycycline hyclate, approximately 70-90% confluent 15-cm dishes of Halo-SPEN-V5-expressing RMCE-*Xist* Δ*Spen* cells were washed twice with 15 mL ice-cold PBS and harvested by scraping into 0.8 mL ice-cold PBS before transferring to a 1.7-mL tube. Cells were pelleted (1200 x g, 5 min, 4 °C) before removing supernatants and storing at −20 °C for later processing. Cell pellets were thawed on ice and resuspended with 10 up-down strokes of a P1000 pipet in 500 μL ice-cold RIPA Lysis Buffer (10 mM Tris-HCl pH 7.5, 140 mM NaCl, 1 mM EDTA-NaOH pH 8.0, 0.5 mM EGTA-NaOH pH 8.0, 1% [v/v] IGEPAL CA-630, 0.1% [w/v] sodium deoxycholate, 0.1% [w/v] SDS; 1 mM PMSF and 1:100 protease inhibitor cocktail [Sigma P8340-5ML] added fresh). Using a Sonics Vibracell 130-watt VCX 130 probe-tip sonicator (30% amplitude output), mixtures were sonicated on ice, twice for 10 s with a 1-min rest in between before pelleting cell debris (16,100 x g, 15 min, 4 °C). Cleared lysates (supernatants) were transferred to new, pre-chilled tubes on ice, and protein concentrations were measured against a standard curve of BSA (Fisher Scientific BP1600-100) via Bradford assay using 1x Bio-Rad Protein Assay Dye Reagent (Bio-Rad 5000006). Antibody-bound beads from the day before were washed twice with 1 mL ice-cold RIPA Lysis Buffer (not containing PMSF or protease inhibitor cocktail) before adding 1000 μg of lysate, brought to a total volume of 500 μL with the addition of RIPA Lysis Buffer. Tubes were incubated overnight at 4 °C with end-over-end rotation. The next day, beads were washed three times with 1 mL ice-cold RIPA Lysis Buffer (not containing PMSF or protease inhibitor cocktail). The final wash was removed, leaving approximately 10-20 μL residual volume on top of the beads, to which 6 μL 4X NuPAGE LDS Sample Buffer (Thermo Fisher NP0007) containing 200 mM DTT was added and mixed by gentle flicking. Samples were heated at 95 °C for 5 min, allowed to cool to room temp, and pulsed down in a microcentrifuge to collect condensation prior to analyzing by SDS-PAGE and western blot.

### Western blots

For western blots not requiring enrichment by immunoprecipitation, lysates were prepared and measured via Bradford assay as described above. Samples for SDS-PAGE were prepared by adding one volume 4X Laemmli sample buffer (Bio-Rad 1610747) containing 10% (v/v) β-mercaptoethanol to three volumes lysate. The exception to this was the right-most western blot in Figure S5B, in which samples were instead prepared with 4x NuPAGE LDS Sample Buffer (Thermo Fisher NP0007) containing 200 mM DTT. All samples were heated at 95 °C for 5 min, allowed to cool to room temp, and pulsed down in a microcentrifuge to collect condensation prior to analyzing by SDS-PAGE and western blot.

For western blots to detect WT or near-full-length SPEN, protein samples in 1x NuPAGE LDS Sample Buffer (Thermo Fisher NP0007) containing 50 mM DTT were run on NuPAGE 3-8% Tris-Acetate pre-cast SDS-PAGE gels (Thermo Fisher EA0375BOX) alongside HiMark Pre-stained Protein Standard (Thermo Fisher LC5699) in 1x NuPAGE Tris-Acetate SDS Running Buffer (Thermo Fisher LA0041) at 150 V and at room temp. Proteins were wet-transferred to methanol-activated Immobilon-P PVDF membranes (MilliporeSigma IPVH00010) in cold 1x NuPAGE Transfer Buffer (Thermo Fisher NP0006) containing 20% methanol, for 1 h 45 min at 110 V and at 4 °C.

For all other western blots, protein samples in 1x Laemmli sample buffer (Bio-Rad 1610747) containing 2.5% (v/v) β-mercaptoethanol were run on TGX pre-cast SDS-PAGE gels (e.g., Bio-Rad 4561084) alongside Precision Plus all-blue protein ladder (Bio-Rad 1610373) in Tris-glycine-SDS running buffer (2.5 mM Tris, 19.2 mM glycine, 0.01% [w/v] SDS, pH 8.3) at 150 V and at room temp. Proteins were wet-transferred to methanol-activated Immobilon-P PVDF membranes (MilliporeSigma IPVH00010) in cold Tris-glycine-methanol-SDS transfer buffer (2.5 mM Tris, 19.2 mM glycine, 20% [v/v] methanol, 0.001% [w/v] SDS, pH 8.3) for 60 min at 100 V and at 4 °C.

After transfer, membranes were frequently cut horizontally with a razor blade to enable parallel detection of loading controls or co-immunoprecipitated proteins (Figure 4D). Membranes were blocked for at least 30 min on a room-temperature orbital shaker in 3% BSA (Fisher Scientific BP1600-100) or 5% nonfat milk (Bio-Rad 1706404) in tris-buffered saline (TBS; 20 mM Tris-HCl, 137 mM NaCl, pH 7.6) before adding primary antibody (see Table S6 for antibodies and concentrations) and incubating overnight on a 4 °C rocking platform. Membranes were washed with a quick rinse of TBS-T and then three times for 10 min in TBS-T on a room-temperature orbital shaker. An appropriate secondary antibody (see Table S6 for antibodies and concentrations) in 3% BSA or 5% milk in TBS was added and incubated for 30 min on a room-temperature orbital shaker. Membranes were washed as before with TBS-T. For detection of proteins probed with HRP-conjugated secondary antibodies, membranes were developed for 5 min with Clarity ECL substrate (Bio-Rad 1705061) or 3 min with SuperSignal West Femto Maximum Sensitivity ECL substrate (Thermo Fisher 34094) before placing in a clear plastic sheet protector to prevent drying while imaging. Membranes were imaged with a Bio-Rad ChemiDoc MP imager in chemiluminescence, DyLight 680, or DyLight 800 mode as appropriate. Protein markers were visualized by simultaneously imaging a second channel in DyLight 680 or AlexaFluor 680 mode.

### *SPEN* “guilt-by-association” co-expression analysis

To enhance prediction accuracy, we took a conservative and multi-tissue/multi-tumor type approach, relying on experimental data from major public datasets and selecting high-quality results, as described below. We used three datasets: Genotype-Tissue Expression (GTEx), The Cancer Genome Atlas (TCGA) Tumor, and TCGA Normal. The first dataset includes normal tissues taken from the entire human body.^155^ The second and third datasets include tumor samples or normal tissues from cancer patients (https://www.cancer.gov/tcga). For the sake of homogeneity and reproducibility, the shared source of data for these datasets was the Gene Expression Profiling Interactive Analysis (GEPIA) database.^156^ This computational resource relies on recomputed data from RNA-seq samples, which were processed through the University of California, Santa Cruz (UCSC) Xena project pipeline.^157^

Co-expression was assessed using the Pearson correlation coefficient of non-log transformed data between *SPEN* and other genes across available samples. Notably, for GTEx, we included only tissues present both in the GEPIA list and in the GTEx V8 version and, at the same time, which are associated, in aggregate, in the GEPIA GTEx list, with > 10 samples. For TCGA Normal and Tumor, the inclusion criterion was to have > 10 samples associated. Due to these criteria, we used 49 different tissues for GTEx, 33 tumor types for TCGA Tumor, and 15 normal tissues for TCGA Normal (see Table S3). We selected, for each admitted tissue of GTEx and of TCGA Normal and for each cancer type of TCGA Tumor, the top 100 most correlated genes with *SPEN* (largest Pearson correlation coefficients, based on the GEPIA data). In case of isoform entries (distinct Ensembl ID) pointing to the same gene symbol, we removed these occasional duplicates and extended the analysis up to reaching the top 100 genes for each tissue, cell or cancer type.

We aggregated these gene entries in three tables having as many genes as the distinct correlated genes found across the tissues/cancers (i.e., union of all genes of the dataset having at least one top correlation) of that dataset and as many columns as the included tissues/cancers (one table for each dataset, tabs in Table S3). This procedure was separately followed for every dataset. While doing so, data were binarized such that the only numerical elements of each column of this table were 1 (gene belonging to the top 100 correlated genes for the analyzed tissue or tumor type) and 0 (otherwise). Then, we calculated the sum of gene hits across all tissues/cancers of that dataset (sums by rows) and used each total as the score for the corresponding gene in that dataset. The motivation for this approach is that, once a correlation is ranked at the very top for a tissue/cancer, the second variable to account for is how frequently we observe it across the dataset. Our assumption was that the more a correlation is common in a dataset, the more likely it is that that correlation is (a) numerically reliable, (b) biologically relevant, and (c) cell- or tissue-independent, and therefore potentially involved in *SPEN* functions in multiple human cell types.

To assess the statistical significance of the score of each gene, separately, for each table, we simulated the outcome of having 100 hits for each column, but randomly distributed in it, and calculated how many times each score was reached, in a simulated row, in a sequence of 1 million (M) simulated matrixes. Therefore, the simulated matrixes had as many rows as those of the three datasets (1,780, 1,144, and 907, respectively) and as many columns as the tissues/cancers of the datasets (see above). The randomization algorithm was applied so that only valid rows were used in the simulation (rows having scores of 0 were not allowed). We used the simulation results, through the following two formulas, to calculate the statistical significance: (a) for all gene scores ≥ maximum score reached in the 1 M randomizations (MS1M), p-value = times MS1M is reached across 1 M randomizations / 1 M; (b) for all gene scores < MS1M, p-value = (times in which scores equaling that score are reached in the 1 M randomizations + times in which scores greater than that score are reached in the 1 M randomizations) / 1 M ^158^. These *p*-values were reassessed in terms of multiple hypothesis testing by using the Benjamini Hochberg false discovery rate (FDR) method ^159^, as implemented in MATLAB ^160^.

The described method to assess the statistical significance of gene scores can be regarded as conservative, since in these simulations we only considered genes having at least one hit (i.e., at least one tissue type in which the gene was a top-100 *SPEN* correlate), instead of the full set of genes for which expression data were available.

In the TCGA Tumor dataset, of the 103 genes significantly correlated with *SPEN*, 59 reside on chr1. These were omitted from comparisons to the GTEx and TCGA Normal datasets and GO analysis because their correlations with *SPEN*, a gene on chr1, cannot be decoupled from cancer-associated chr1 copy-number variations that presumably have a strong influence in this dataset.

Overlaps of genes among the three datasets and depictions via area-proportional Venn diagram were generated using DeepVenn (https://www.deepvenn.com/, ^161^).

To assess whether this method significantly enriches for genes that encode SPEN-interacting proteins, we first created a merged list of our 37 WT SPEN interactors (Table S2), our 146 SPEN RRM1-4 interactors (Table S2), and 207 SPOC domain interactors (“Ratio” value >2, ^20^). We removed all duplicate entries, resulting in a list of 347 genes. We matched these 347 genes, by gene symbol, with all genes that had collected at least one hit across the 49 GTEx tissue/cell types of our analysis. To prevent bias when matching gene symbols from distinct sources and of mixed nature (due to the presence of coding as well as non-coding genes in the GTEx dataset), this procedure was performed with respect to a pre-defined common set, including only coding genes recognized by the Human Genome Organization (HUGO, https://www.genenames.org/). We assessed if the matching genes were enriched or depleted for genes having the highest scores in the GTEx analysis (those with FDR < 0.10). For this task, we compared the proportion of HUGO genes with FDR < 0.10 in the matched list and in the full GTEx list (used as baseline). Those two proportions were compared through a two-tailed binomial test using R.

### *Xist* O-MAP

For *Xist* O-MAP, RMCE-*Xist* WT and Δ*Spen* mESCs were cultured as above. *Xist* expression was induced with 1 µg/mL doxycycline in all of 4 replicates for 72 h prior to O-MAP fixation. Cells were plated into 6-well plates (3.5x10^5^ cells per well; 6 wells per replicate) 24 hours prior to O-MAP. *Xist* O-MAP labeling was conducted in parallel on all replicates, as previously described ^97,162^, using *Xist*-targeting primary probes, and the SABER1–HRP O-MAP secondary probe described previously.^97^ Briefly, cells were fixed (2% paraformaldehyde, 10 min), permeabilized (0.5% Triton X-10, 10 min), and endogenous peroxidases were inactivated (0.5% H_2_O_2_, 10 min). Primary *Xist* O-MAP probes ^97^ were hybridized for 8 hours at 42 °C, in 40% formamide Hybridization Buffer.^97^ Cells were washed to remove excess unbound probes and hybridized to the HRP-conjugated O-MAP oligo (1 hour at 37 °C; 30% formamide). Samples were extensively washed, and *in situ* biotinylated (8 µM biotin tyramide; 1 mM H_2_O_2_; 10 min at 23 °C). Labeling reactions were quenched with sodium azide and ascorbate, and cells were harvested by scraping into 1x Dulbecco’s Phosphate Buffered Saline (dPBS), pooled, and stored at −80°C before use.

### *Xist* O-MAP-MS sample preparation

*Xist* O-MAP labeled cells (4.2×10^6^ cells per replicate; 4 biological replicates per experimental condition) were lysed, processed for streptavidin pulldown, and prepared for mass spectrometry using the same manipulations and reagents described previously.^162^ Briefly, cells were gently lysed (0.5% NP-40, 10 min) and centrifuged to remove supernatant (3000 x g, 5min at 4 °C, 2 times) for nuclear enrichment. Nuclei were more extensively lysed (0.1% SDS, 10 min) and sonicated with a Branson Probe Sonicator (4 cycles of 30 s at 0.7/1.3 seconds on/off intervals) in an ice block. Samples were reverse-crosslinked by boiling (1% SDS, 1 hour, 800 rpm mixing), cooled to 50°C, then sonicated for one additional cycle on ice, using the same parameters. Samples were centrifuged (15,000 x g, 10 min at 4 °C), and clarified lysates were gently mixed with Pierce Magnetic streptavidin beads (ThermoFisher 88817; 75 µL per sample) for 2 h, with end-over-end rocking, at 23 °C. Beads were washed twice with (Tris-HCl, 0.1% SDS, sodium azide, NaCl) and samples were moved to PCR strip tubes to improve sample recovery. Samples were then washed with the following series of buffers, with one-minute, off-magnet resuspensions between each washing step: (Tris-HCl, 0.1% SDS, sodium azide; two washes), 1x PBS (two washes), 1 M KCl Buffer (two washes), 2 M Urea Buffer (four washes), EPPS Buffer (two washes). Beads were then resuspended in new PCR strip tubes in 50 μL EPPS buffer. Samples were reduced (10 mM TCEP, 60 min at 700 rpm mixing), alkylated (10 mM IAM, 60 min at 700 rpm in the dark) and quenched with DTT (5 mM, 15 min at 700 rpm). Samples were then digested with LysC (1 μg per 100 µg total protein; 3 hours at 37 °C, 700 rpm mixing) and then trypsin (1 μg per 100 µg total protein; overnight at 37 °C, 700 rpm mixing). Beads were removed magnetically and samples were moved to fresh 0.2 mL PCR-strip tubes. Peptides were stage-tipped and analyzed by mass-spectrometry as described.^162^

### EU-RNA-seq sample preparation

The following protocol is a modification of others published previously,^104,163^ modified to include both unlabeled and biotinylated spike-in control RNAs to assess enrichment of nascent RNA, to detail sequencing library preparation, and to provide further clarity.

Spike-in control RNAs were generated by in vitro transcription of linearized control plasmids with the MEGAscript SP6 Transcription Kit (Thermo Fisher AM1330), following manufacturer’s protocol with 20-µL reactions. Unlabeled spike-in RNA was transcribed from 1 µg linearized pGEM Express (Promega P2561) with 5 µM each standard NTP. Biotin-labeled spike-in RNA was transcribed from 1 µg pTRI-Xef (Thermo Fisher AM1330) with 5 µM CTP, ATP, and GTP; 3.75 µM UTP; and 1.25 µM biotin-16-aminoallyl-UTP (TriLink Biotechnologies N-5005). Both reactions contained 0.5 µL 40 U/µL RNaseOUT. Reactions were incubated at 37 °C for 4 h on a thermal cycler, after which 1 µL TURBO DNase was added and mixed well before incubating at 37 °C for 15 min. In-vitro-transcribed RNAs were purified with the Monarch RNA Cleanup Kit (NEB T2050), eluting with 50 µL nuclease-free water. RNA concentrations were measured by Qubit (RNA Broad-Range Kit), and RNA size and integrity were confirmed by formaldehyde agarose gel electrophoresis. Aliquots containing 100 pg of both spike-in RNAs were prepared and stored at - 80 °C.

To metabolically label nascent cellular transcripts, the growth medium of approximately 80% confluent E14 WT cells (duplicate platings) and E14 Δ*Spen* cells (one plating each of two independent clonal lines) in 6-cm dishes was replaced with fresh growth medium containing 0.5 mM 5-ethynyluridine (EU, diluted from a pale-yellow 100-mM stock dissolved in DMSO, supplied in the Click-iT Nascent RNA Capture Kit, Thermo Fisher C10365). Exactly one hour later, growth medium was removed, cells were washed once with 3 mL room-temp PBS, and RNA was harvested in 1 mL TRIzol. RNA was prepared via standard TRIzol-chloroform extraction, according to manufacturer’s instructions, and resuspended in RNase-free water. RNA concentrations were measured by Qubit (RNA Broad-Range Kit), and rRNA integrity was confirmed by agarose gel electrophoresis.

To biotinylate EU-labeled RNA, 50-µL click reactions were assembled in 1.7-mL tubes at room temp following the Click-iT Nascent RNA Capture Kit manufacturer’s instructions, with each containing 3.15 µg EU-labeled RNA and 0.5 mM biotin azide; buffer additive 2 was added approximately 4 minutes after adding buffer additive 1, upon which reactions turned an intense reddish brown. Tubes were then immediately placed on a Vortex Genie 2 (Fisher Scientific 12-812) fitted with a foam tube holder attachment (Scientific Industries 504-0234-00) and gently shaken with a vortex intensity of 3 at room temp for 30 min. Tubes were removed and pulsed down to collect liquid, and then the following were added, in order: 1 µL 5 mg/mL linear acrylamide (Thermo Fisher AM9520), 50 µL 7.5 M ammonium acetate, and 700 µL ice-cold 100% ethanol. Tubes were inverted and vortexed for 3-5 s before incubating overnight (20 h) at −80 °C to precipitate RNA.

The next day, tubes were centrifuged at 13,000 x g for 20 min at 4 °C. Supernatants were removed carefully to avoid loss of the very small RNA pellets and washed twice with 700 µL ice-cold 75% ethanol, vortexing 3 s and centrifuging at 13,000 x g for 10 min at 4 °C each wash. The final wash was removed first by P1000 pipet and then pulsing down and carefully removing residual wash by P20 pipet. RNA pellets were air-dried for about 5 min, after which 10 µL ice-cold nuclease-free water was added. Tubes were incubated on ice for 1 h and then mixed by vortex and pipetting up and down with a P20 pipet. Concentrations of each resuspended RNA were measured by Qubit (RNA Broad-Range Kit). In 0.2-mL PCR strip-tubes, 1 µg biotinylated EU-labeled RNA was mixed with 100 pg of both spike-in control RNAs, brought to 12.3 µL with nuclease-free water, and stored at −80 °C.

The following day, biotinylated RNAs were enriched with streptavidin beads and used to generate RNA-seq libraries as follows. Using a magnetic separator, a volume of well-mixed Dynabeads MyOne T1 streptavidin beads equivalent to 5.5 μL per sample was washed three times with 500 μL room-temp Click-iT wash buffer 2. The beads were then resuspended to the original volume with room-temp Click-iT wash buffer 2, mixed well, and 5 μL washed beads were added to a new room-temp 1.7-mL tube for each sample.

RNA mixtures from the day before were thawed at room temp, and 12.7 µL of a room-temp master mix containing 12.5 µL 2x Click-iT RNA binding buffer and 0.2 µL 40 U/µL RNaseOUT was added and mixed well by P20 pipet. Tubes were incubated on a thermal cycler at 69 °C for 5 min and then allowed to returned to room temp. The entire 25-µL RNA samples were added to tubes containing 5 μL washed streptavidin beads, mixed well by pipet (keeping volume at the bottoms of the tubes), and incubated at room temp on an end-over-end rotator to prevent bead settling. Tubes were placed on a magnetic separator, and supernatants were removed by pipet. With aid of the magnetic separator to remove supernatants, beads were washed five times with 50 µL room-temp Click-iT wash buffer 1 and then five times with 50 µL room-temp Click-iT wash buffer 2, using a pipet to resuspend beads thoroughly at each wash step. To fully remove the final wash, tubes were briefly pulsed down, and residual wash was removed with a P20 pipet. Washed beads were resuspended in 5 µL room-temp Click-iT wash buffer 2 and transferred to 0.2-mL PCR strip-tubes.

Proceeding immediately, streptavidin-captured RNA was used for on-bead cDNA generation with the KAPA RNA HyperPrep Kit (Roche 08098140702) as follows. A master mix was prepared containing 10 µL 2x Fragment, Prime, and Elute Buffer and 5 µL nuclease-free water per sample (with 10% excess to allow for pipetting error), and 15 µL of this master mix was added to each 5-µL bead sample, mixing well by pipet. Samples were mixed again briefly by pipet (to keep beads in suspension) immediately before placing on a thermal cycler pre-warmed to 65 °C for 5 min. Tubes were then immediately placed on ice, and 10 µL first strand synthesis master mix was added to each and mixed well by pipet. Beads were resuspended by pipet immediately before placing on a thermal cycler with the following program: 25 °C for 10 min, 42 °C for 15 min, 70 °C for 15 min. Every 5 min during this program, the lid of the thermal cycler was opened, and beads were resuspended by pipet (∼3 quick up-down strokes) to keep in suspension. Tubes were returned to ice, and 30 µL second strand synthesis and A-tailing master mix was added to each and mixed well by pipet. Beads were resuspended by pipet immediately before placing on a thermal cycler with the following program: 16 °C for 30 min, 62 °C for 10 min. Every 5 min during this program, the lid of the thermal cycler was opened, and beads were resuspended by pipet (∼3 quick up-down strokes) to keep in suspension. After completion of second-strand cDNA synthesis, cDNA should be uncoupled from RNA immobilized on the beads. The entire mixtures were transferred to new 1.7-mL tubes, and supernatant cDNA was removed from beads using a magnetic separator and transferred to pre-chilled 0.2-mL PCR strip-tubes on ice. From this step onward, the KAPA RNA HyperPrep Kit manufacturer’s protocol was followed to generate RNA-seq libraries, using 1.5-μM adapter stocks 12 cycles of PCR for library amplification. Libraries were quantified by Qubit (dsDNA broad sensitivity kit, Thermo Fisher Q32850), and average fragment sizes were determined by agarose gel electrophoresis. Libraries were pooled and sequenced on an Illumina NextSeq1000 platform with a 100-cycle NextSeq 1000/2000 P2 XLEAP-SBS Reagent Kit (Illumina 20100987).

### EU-RNA-seq data processing and analysis

Using STAR (v2.7.11b, ^141^), EU-RNA-seq reads were aligned to a custom genome index containing the mouse GRCm38 (mm10) genome and sequences of the two linearized in-vitro transcription control plasmids (with reference to GTF file gencode.vM25.basic.annotation.gtf from https://www.gencodegenes.org/mouse/release_M25.html ^142^). Alignment was performed with single-pass mapping and the option “--outFilterMultimapNmax 1” to consider only uniquely mapping reads. Reads were assigned to each gene using featureCounts (Subread v2.1.1, ^143^) and a custom SAF file that contained sequences of both linearized in-vitro transcription control plasmids and definitions of each vM25 gene corresponding to its start and end position (i.e., ignoring exon definition to enable counting of nascent transcription reads originating from all exons and introns of a gene). In the output file, across samples, the number of reads assigned to pGEM Express (unlabeled IVT control) was 8-10% the number of reads assigned to pTRI-Xef (biotinylated IVT control), indicating uniform, successful enrichment of biotinylated (nascent) RNA. Mouse genes were pre-filtered to 11306 genes for which an average of 50 reads were assigned across the samples. DESeq2 (v1.32.0, ^49^) was used to determine differentially transcribed genes between WT and Δ*Spen* cells and to visualize by MA plot, using a significance threshold of p_adj_ < 0.05.

### SPEN RRM1-4 RIP-seq signal summing by gene

To determine the extent to which genes had transcripts associating with SPEN RRM1-4, the Halo RIP-seq peaks over the *Spen* locus were removed in the same fashion as the *Spen* and *Xist* peaks from the motif methods above. Next, a limited version of the GENCODE vM25 basic annotation GTF was created that removed all non-gene feature lines, and this limited GTF was provided to BEDTools intersect with the non-*Spen* peaks and the -wo and -s flags. The resultant output file contained one line for every peak that intersected a gene, the information of both the gene and the peak coordinates, and the number of nucleotides in that intersection. A custom Python script (rank_genes_by_peak_signal_3_31_25.py) was used to calculate the scaled signal over control for each peak as was calculated in the motif methods previously (the RIP RPM value squared and divided by the sum of the RIP RPM and the control-sample RIP RPM). For each gene, the scaled signals for all peaks over the annotated gene bounds were fractionally summed based on the percentage of the peak within the gene bounds.

### RNA stability assay

RMCE-*Xist* WT and mutant cells in wells of a 6-well plate were treated for 4 h with 5 μg/mL actinomycin D or an equivalent volume of DMSO vehicle control (“0 h”) before washing once with 1x PBS and harvesting with 1 mL TRIzol. RNA was prepared via standard TRIzol-chloroform extraction, according to manufacturer’s instructions, and resuspended in RNase-free water. 400 ng RNA was reverse-transcribed with random primers using the Applied Biosystems High-Capacity cDNA Reverse Transcription Kit (Thermo Fisher 4368814; see above) and analyzed by qPCR (see above) to measure *Xist* and *Gapdh* mRNA abundance, fitting to a serially-diluted standard curve of a representative sample for each primer pair to incorporate primer efficiencies (see Table S6 for primer sequences). *Gapdh*-normalized *Xist* abundance data were calculated relative to 0-h control, and results from three independent experiments were with GraphPad Prism (v9.5.0).

### Semiquantitative PCR analysis of *Xist* ectopic intron splicing

One microgram of total RNA from RMCE-*Xist* WT or intron mutant cells (see RNA-seq section) was reverse-transcribed with random primers using the Applied Biosystems High-Capacity cDNA Reverse Transcription Kit (Thermo Fisher 4368814; see above), and products were diluted 10-fold with nuclease-free water. PCR reactions (10 μL) were prepared containing 5 μL iTaq Universal SYBR Green Supermix (Bio-Rad 1725124), 3 μL nuclease-free water, 0.5 μL 10 μM forward primer JT328, and 0.5 μL 10 μM reverse primer JT329 (see Table S6 for primer sequences). For PCR templates, 1 μL 1:10-diluted reverse transcription product was added, or in the case of the “spliced” and “unspliced” controls, 1 μL 0.5 ng/μL pTETRISv1-*Xist*-2kb or pTETRISv1-*Xist*-2kb-β-globin/IgG-intron,^27^ respectively. PCR was run on a thermal cycler with the following program: 95 °C for 10 min, 24 cycles of (95 °C for 15 s, 60 °C for 30 s, 72 °C for 30 s, 12 °C hold. Twenty-four cycles was chosen due to this being in the “linear range” of amplification curves from qPCR analyzed previously; parallel semiquantitative PCRs were run with 22 and 26 cycles, which yielded similar results. PCR products were mixed with Purple Gel Loading Dye (NEB B7024S) to 1x concentration and run on a 2% agarose-TAE gel containing ethidium bromide and visualized with a Bio-Rad ChemiDoc MP imager (Ethidium Bromide setting) alongside 1 Kb Plus DNA Ladder (Thermo Fisher MPK10025). Band intensities were determined by densitometry with Bio-Rad Image Lab 6.1 software. For each sample, splicing efficiency was determined as the ratio of spliced band intensity to the sum of spliced and unspliced band intensities.

### Key Resources Table

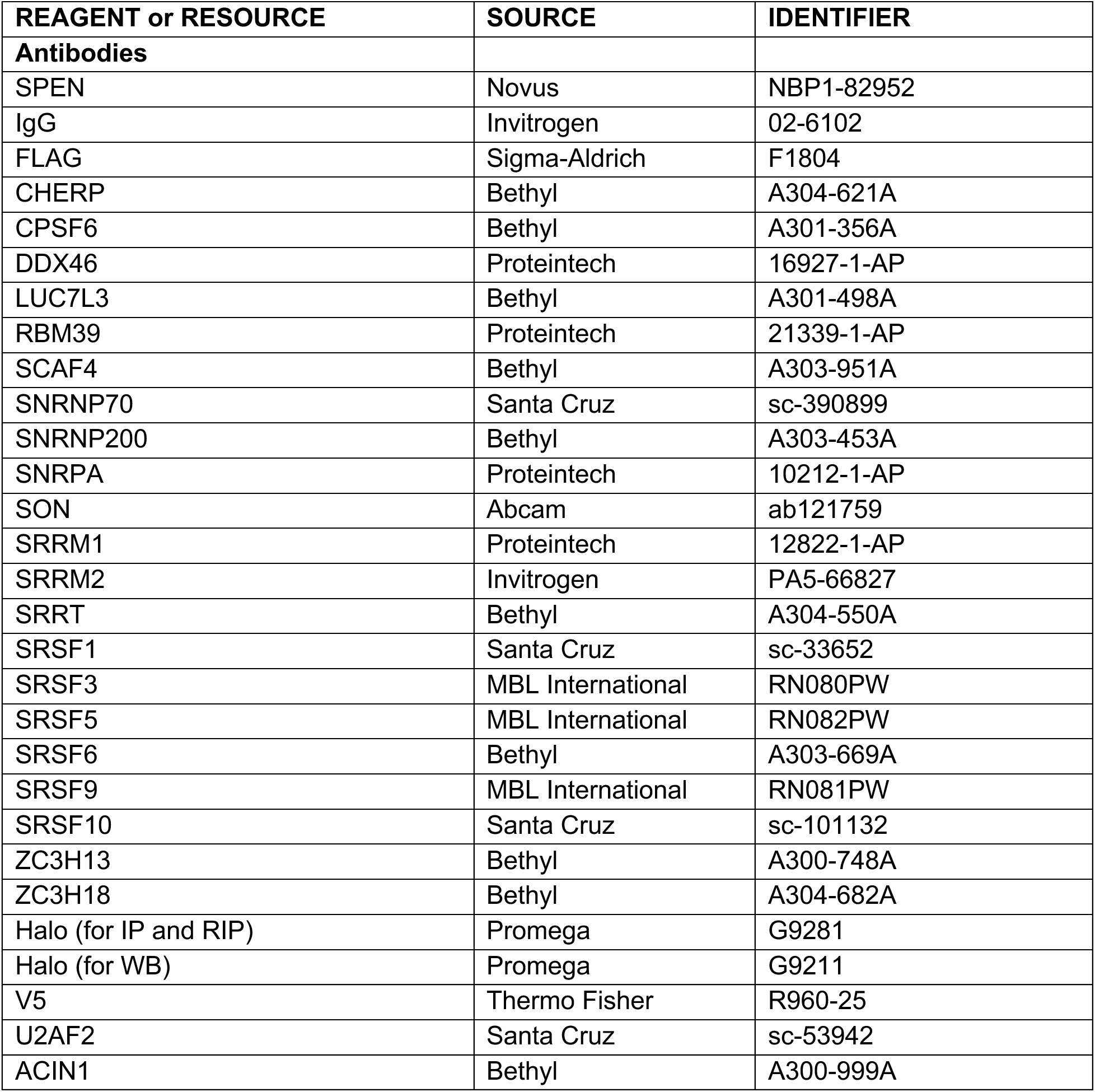
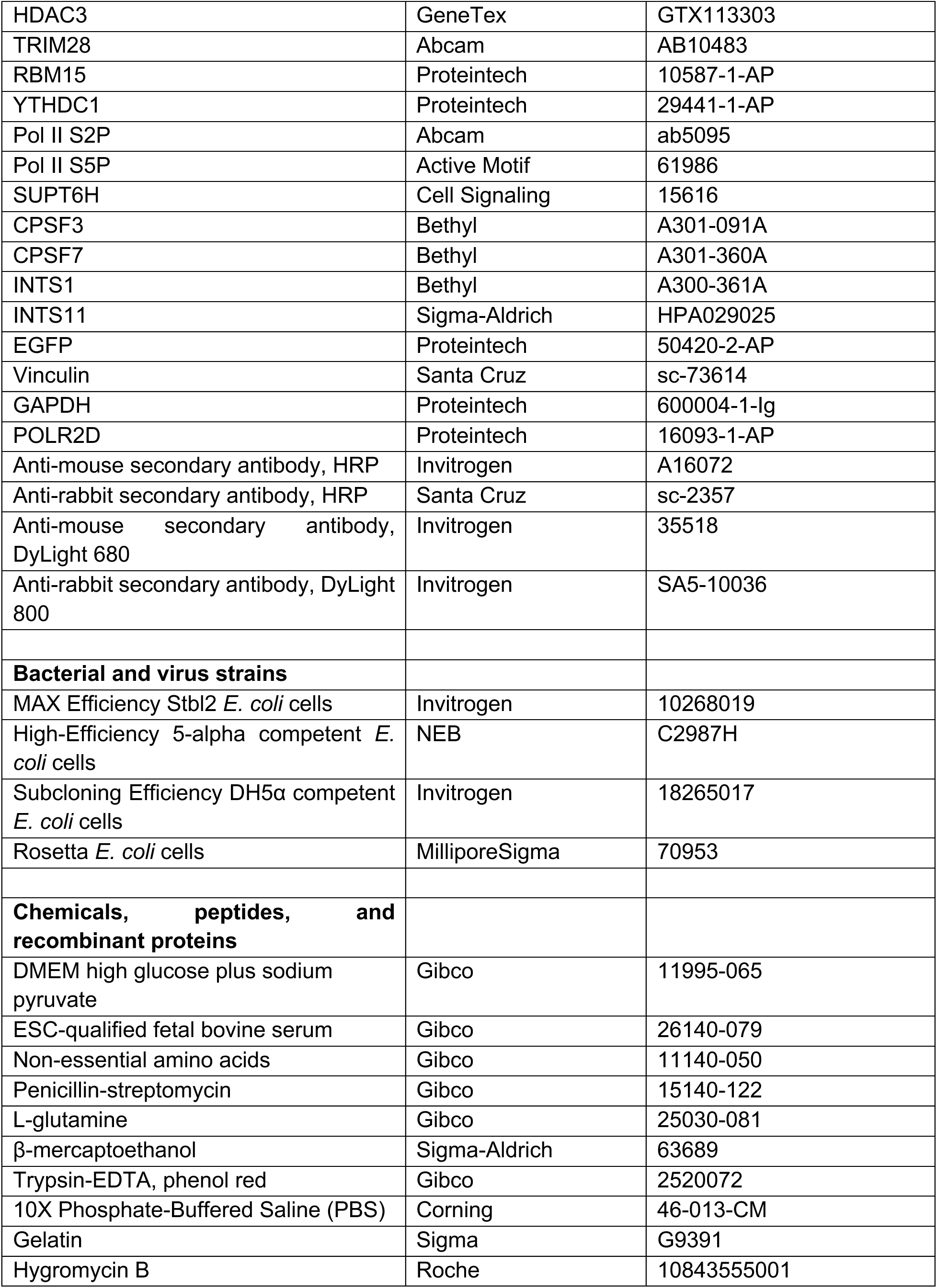
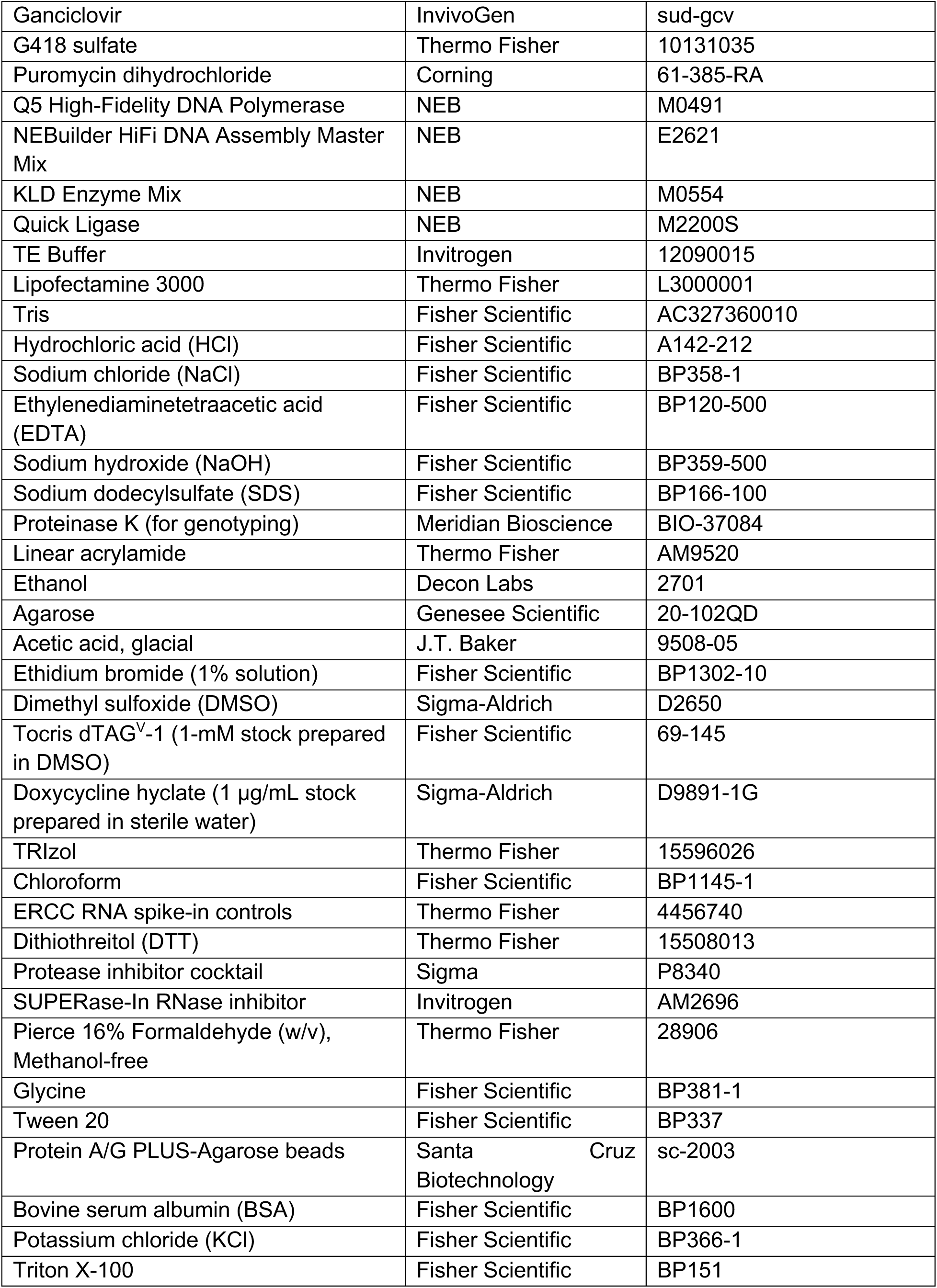
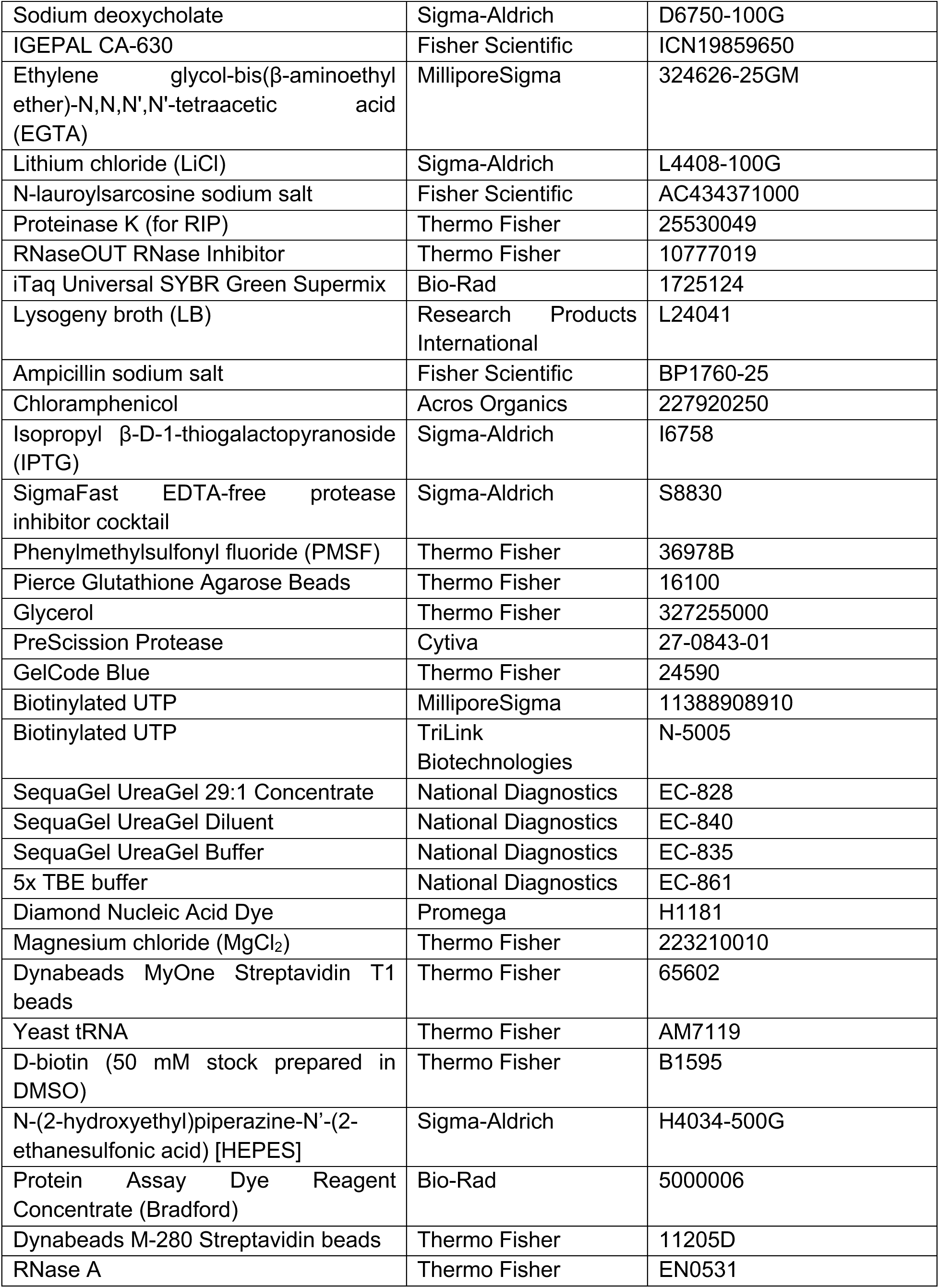
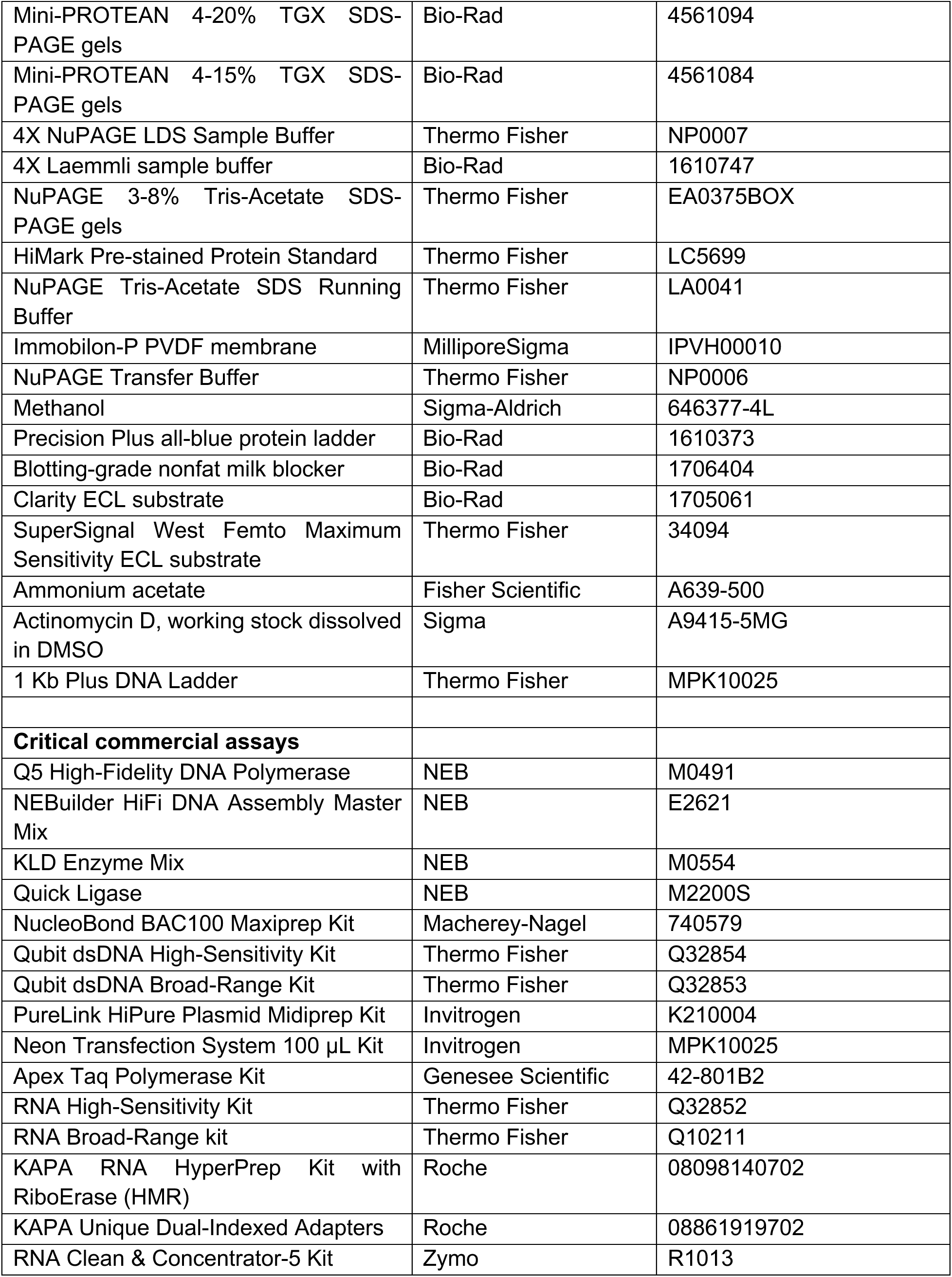
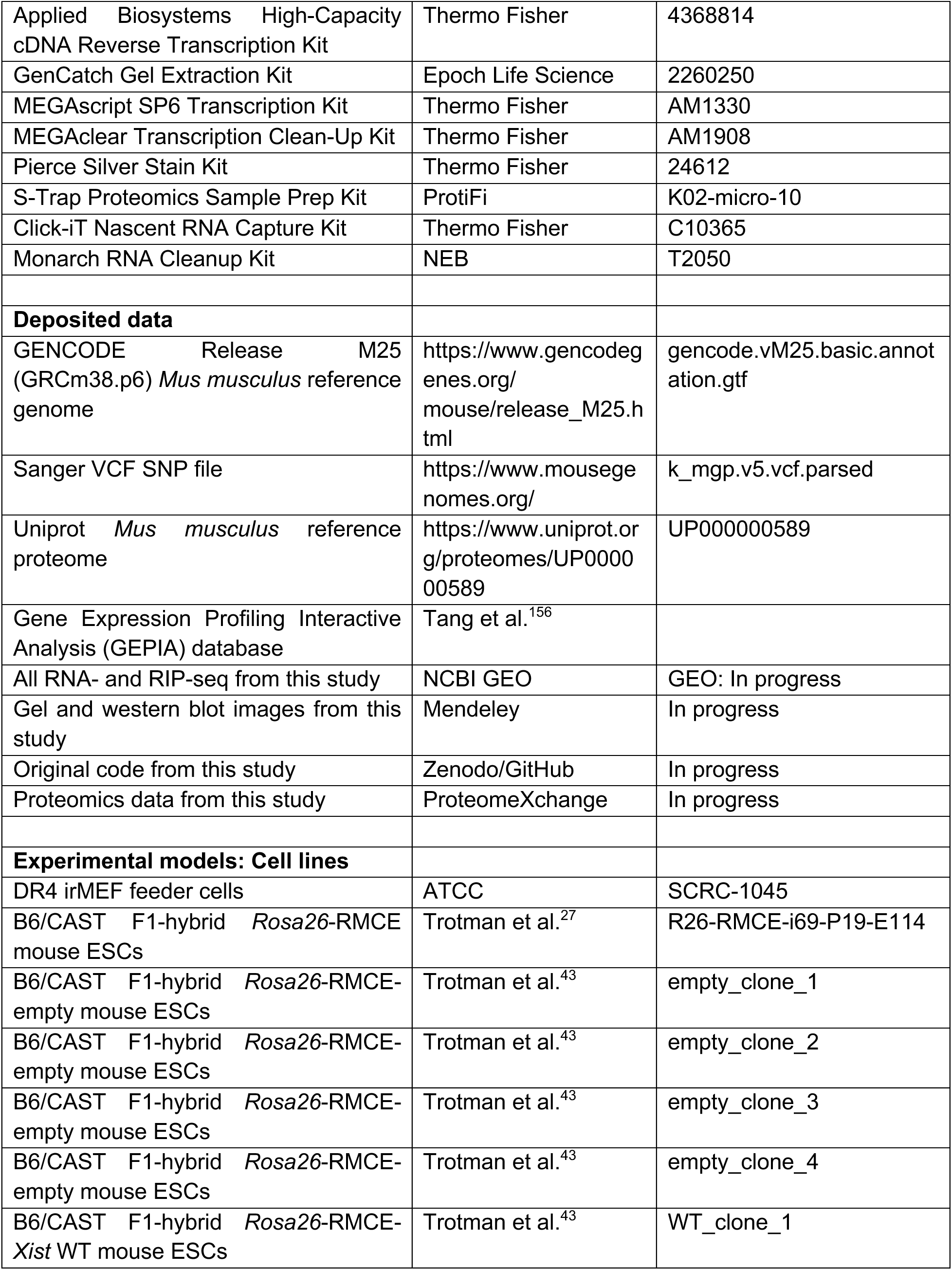
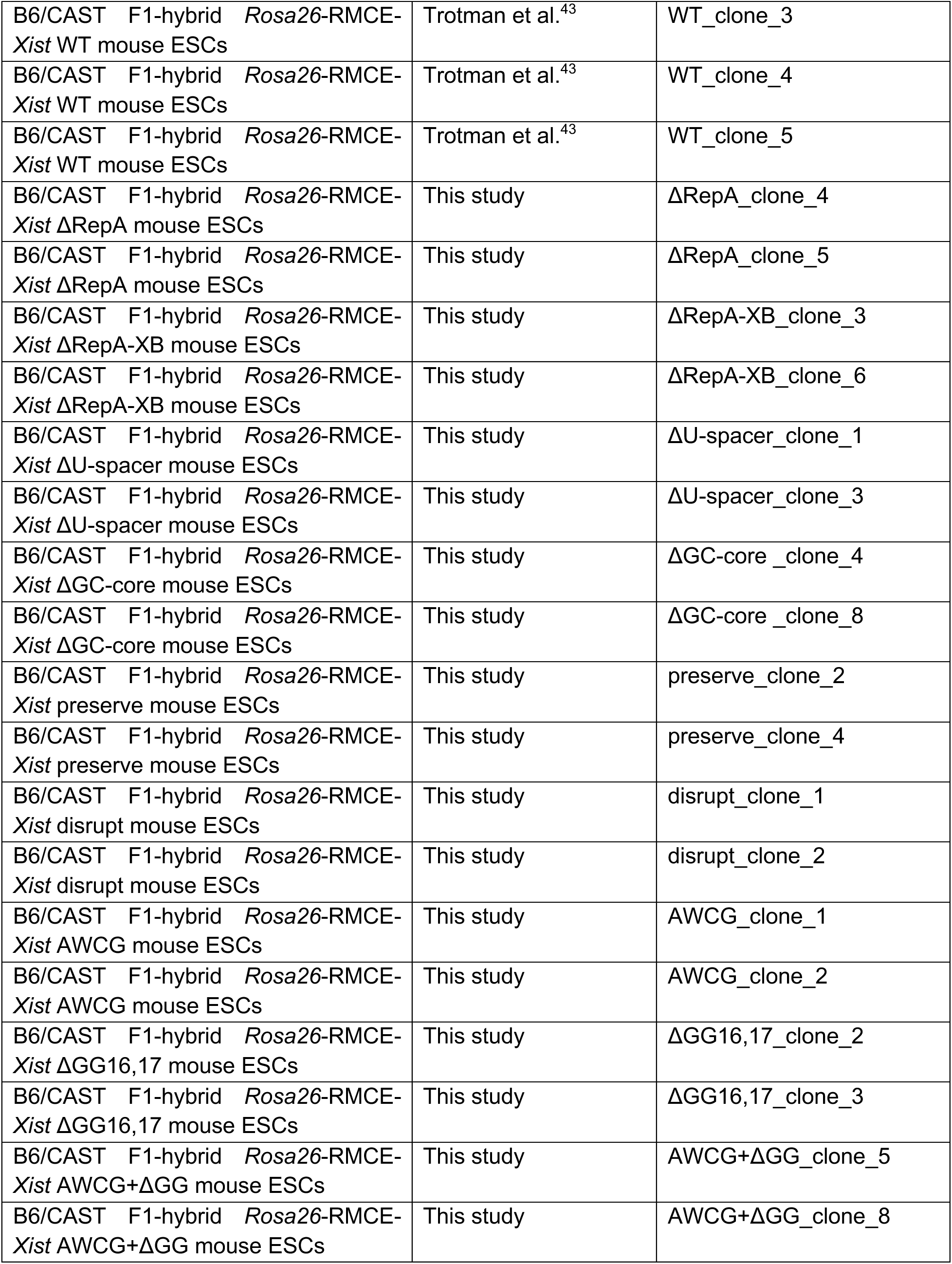
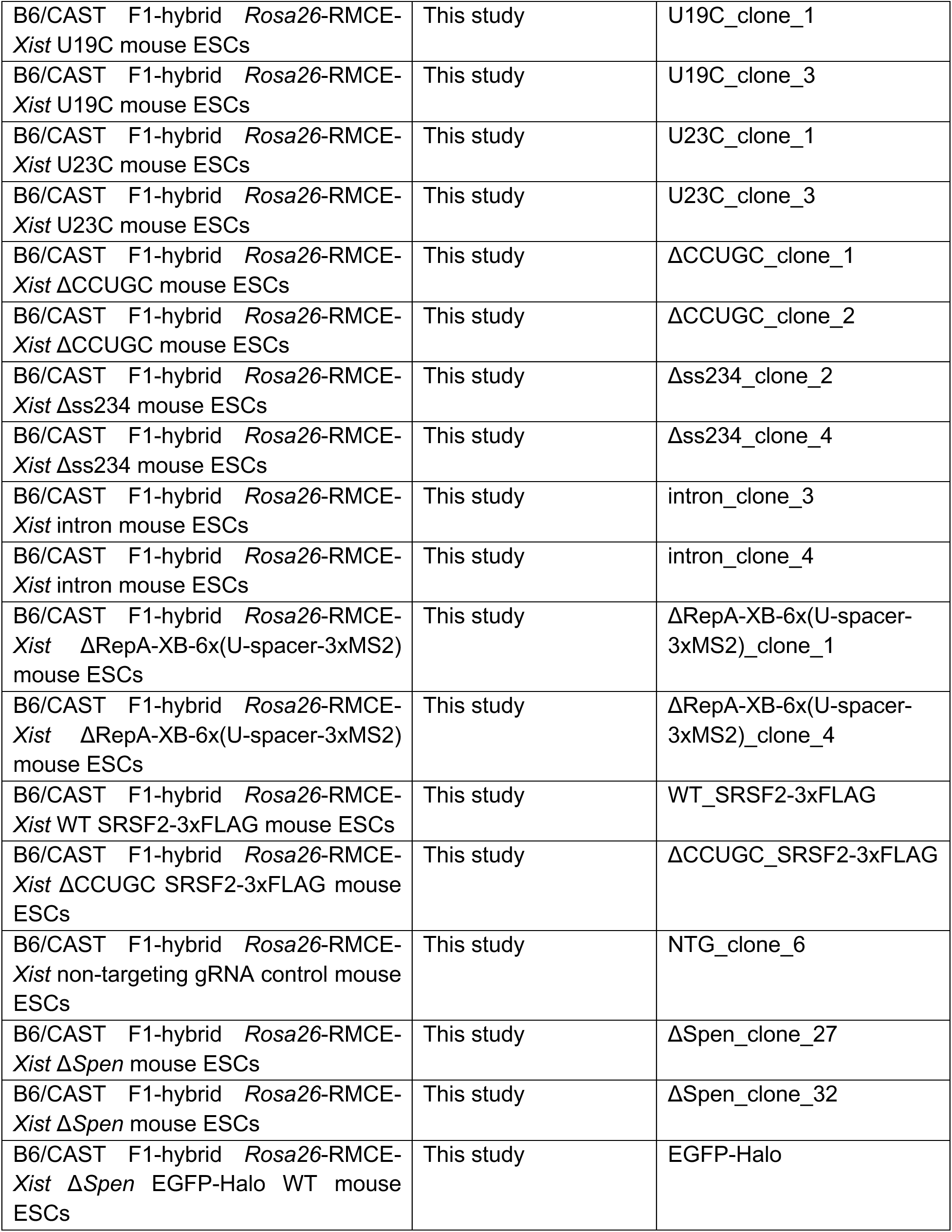
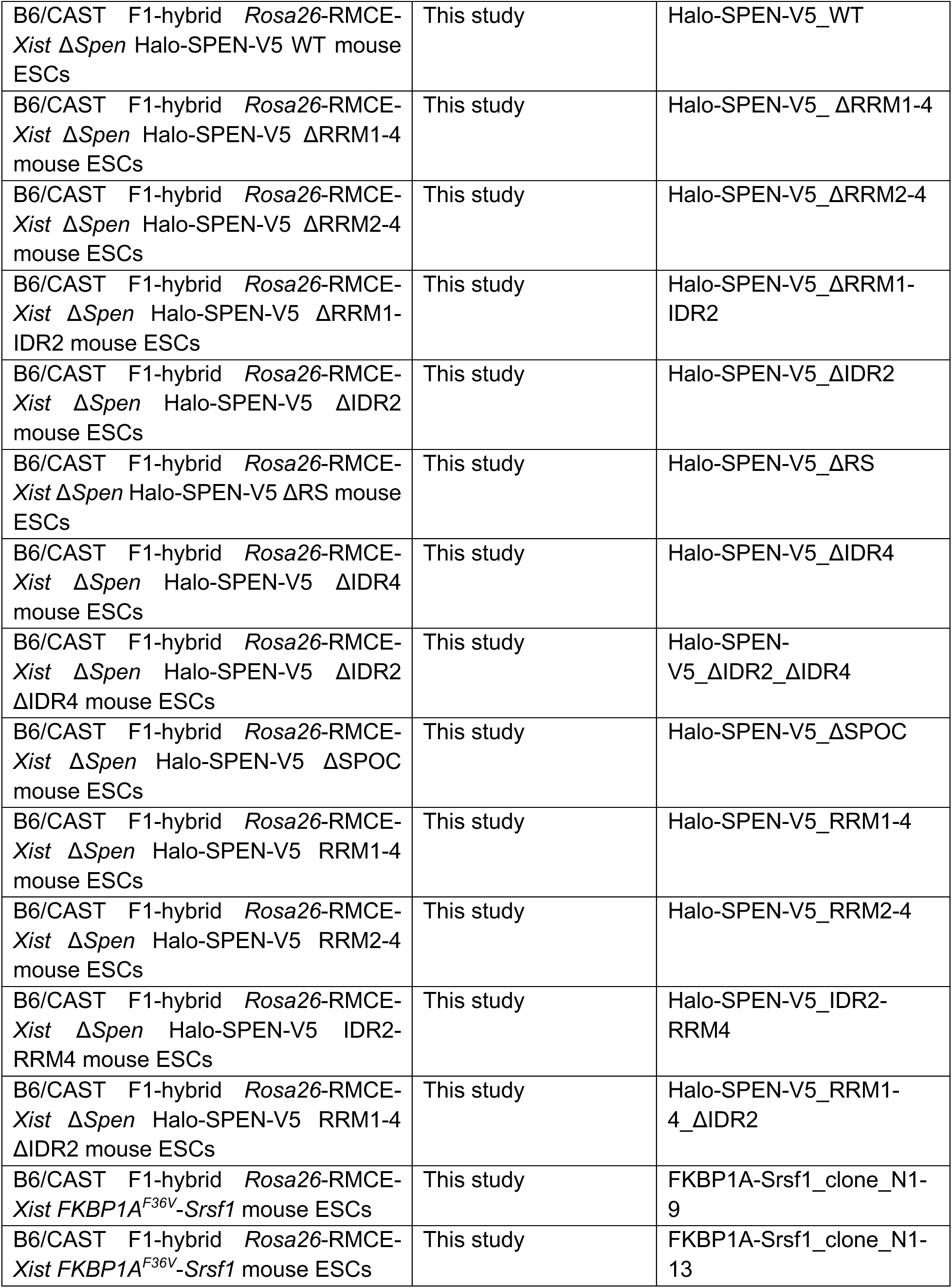
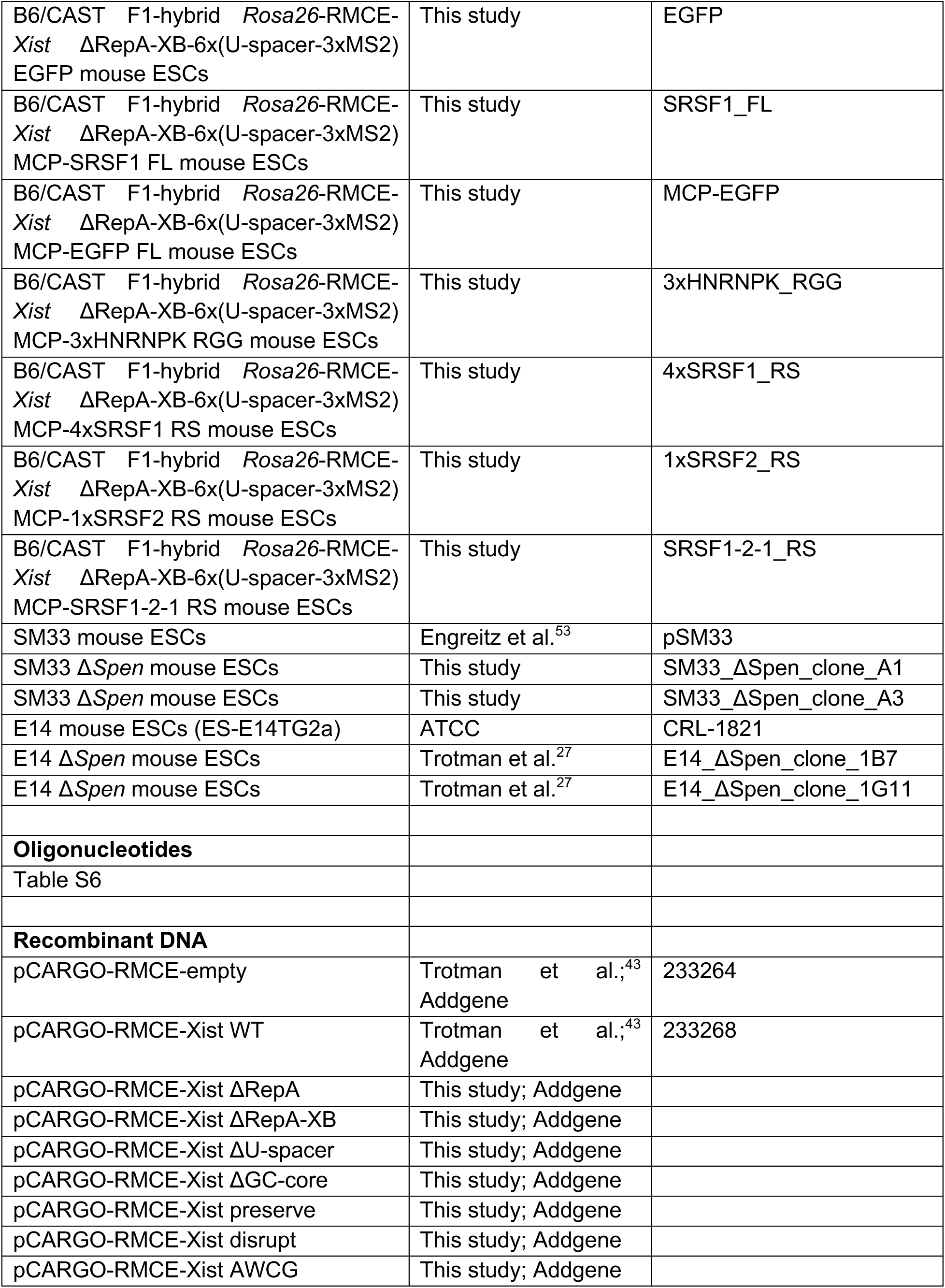
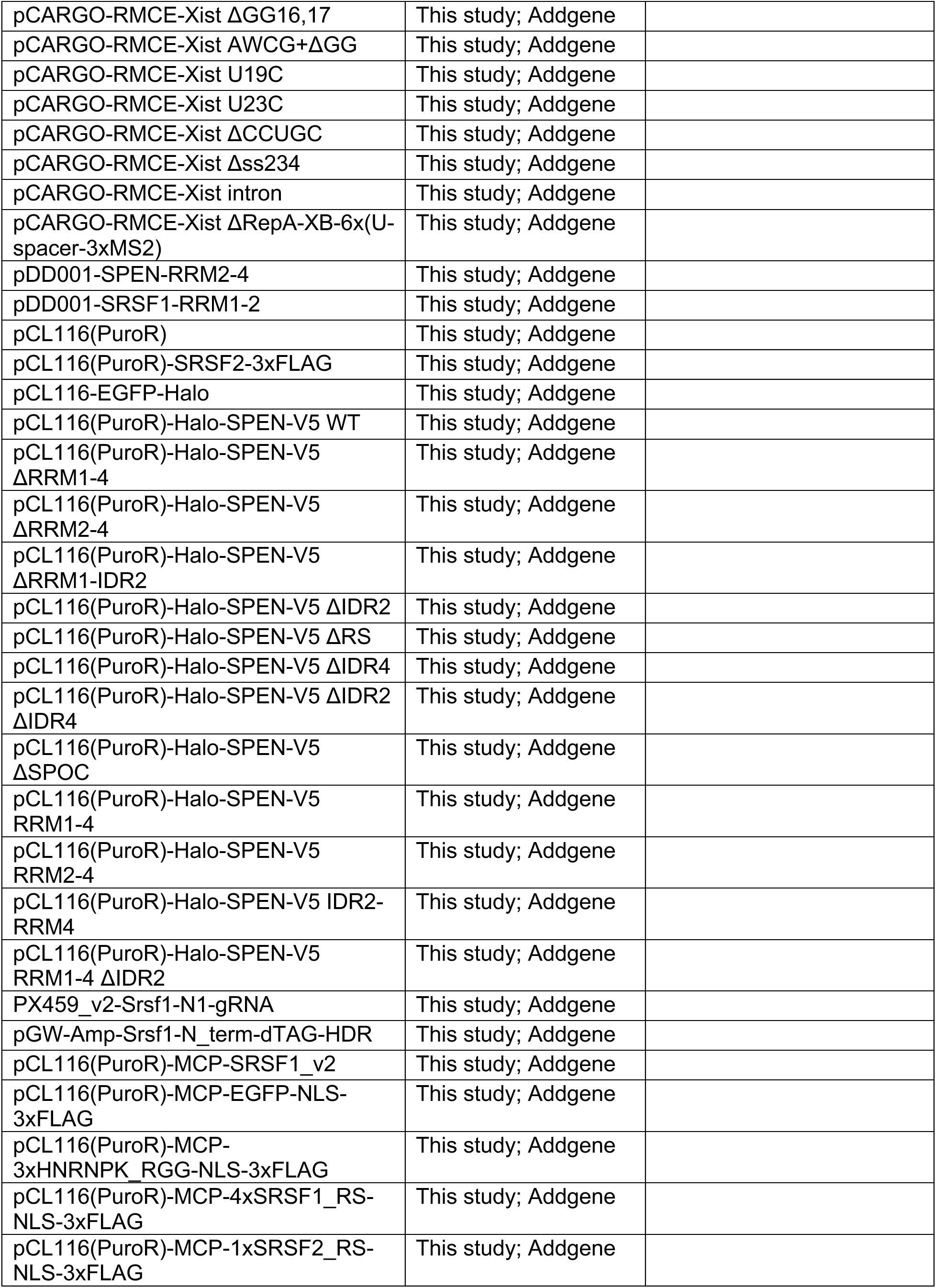
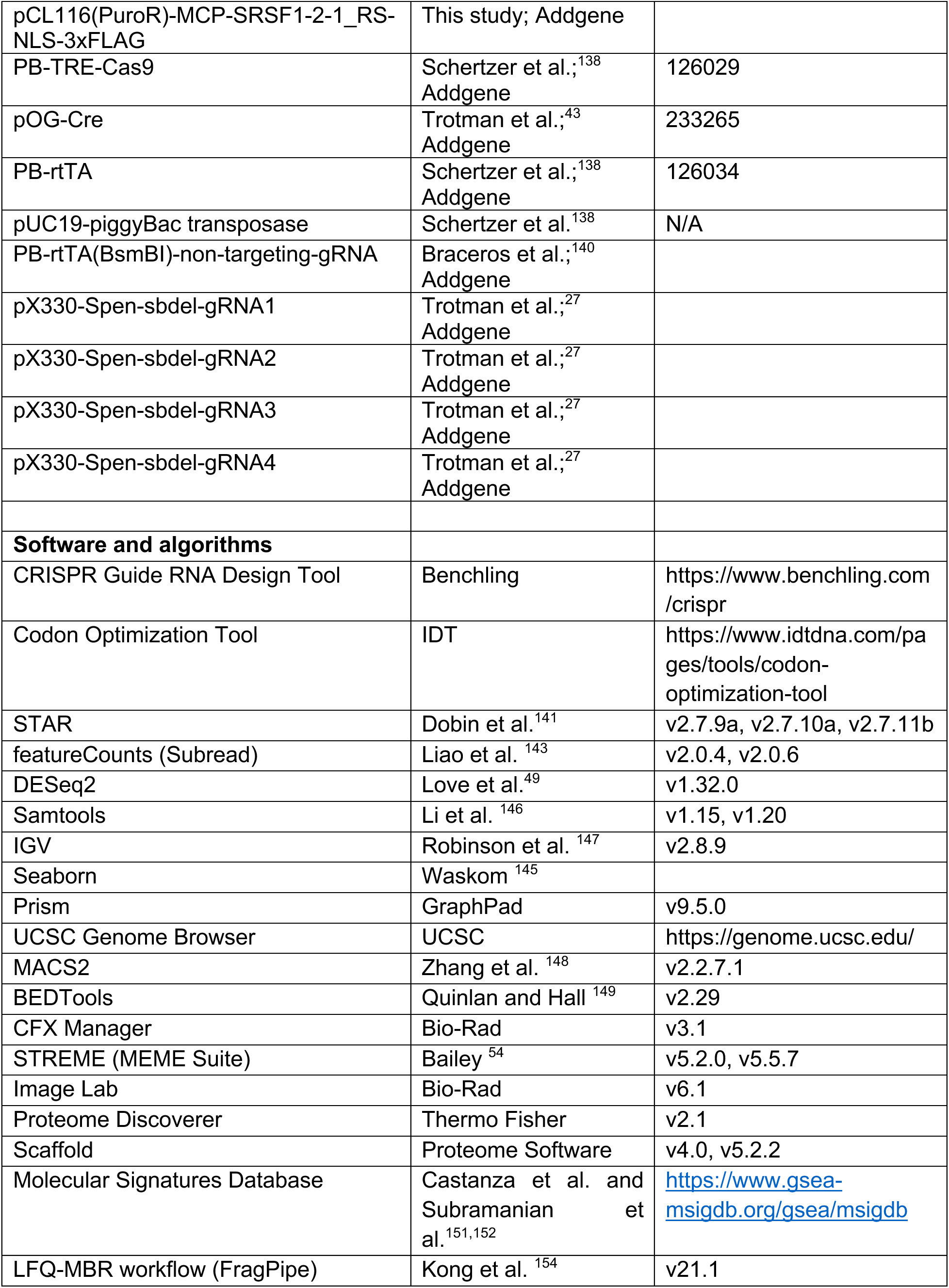
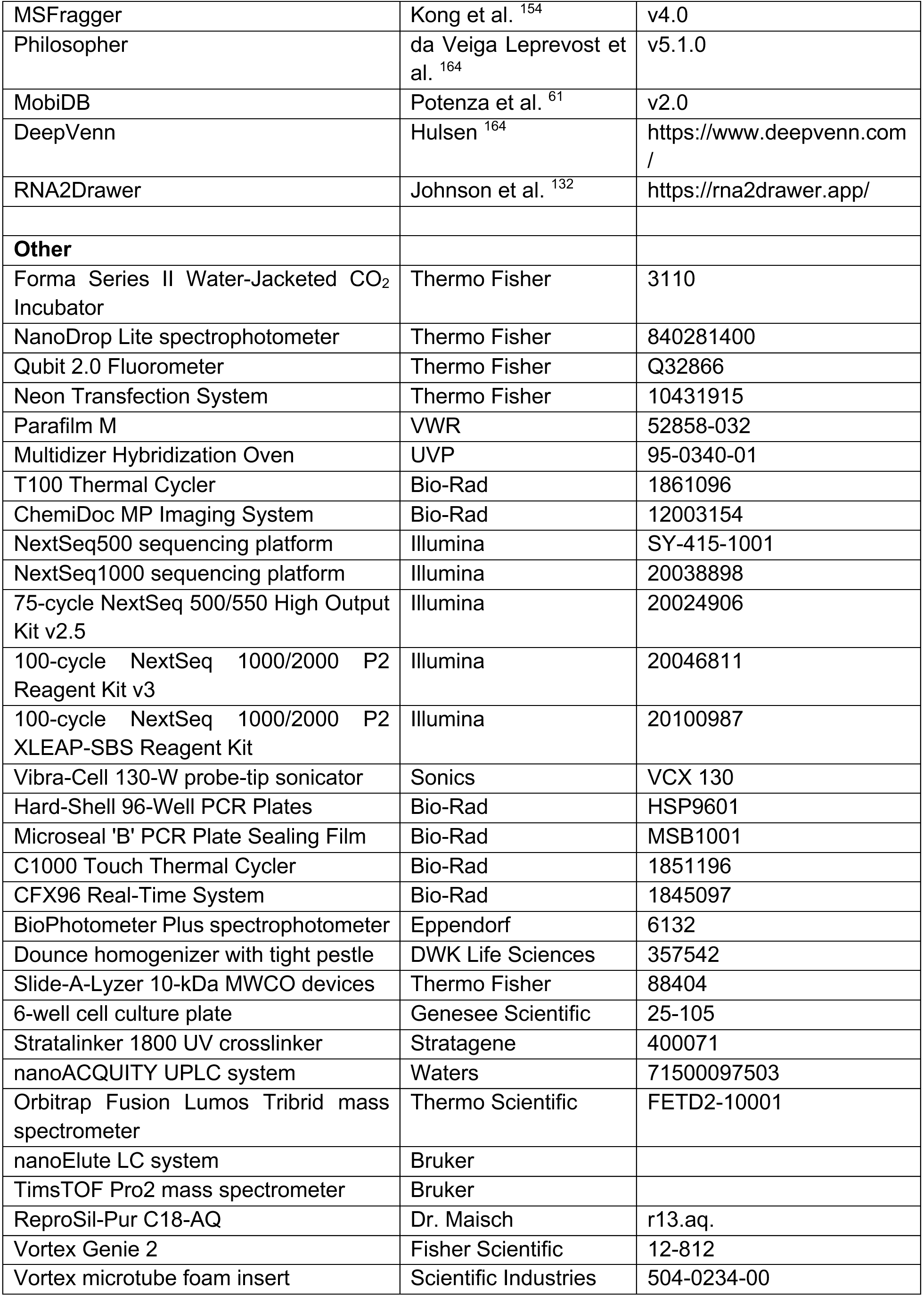

## REFERENCES

1. Loda, A., Collombet, S., and Heard, E. (2022). Gene regulation in time and space during X-chromosome inactivation. Nat Rev Mol Cell Biol 23, 231–249. 10.1038/s41580-021-00438-7.

2. Mattick, J.S., Amaral, P.P., Carninci, P., Carpenter, S., Chang, H.Y., Chen, L.L., Chen, R., Dean, C., Dinger, M.E., Fitzgerald, K.A., et al. (2023). Long non-coding RNAs: definitions, functions, challenges and recommendations. Nat Rev Mol Cell Biol 24, 430–447. 10.1038/s41580-022-00566-8.

3. Pintacuda, G., Wei, G., Roustan, C., Kirmizitas, B.A., Solcan, N., Cerase, A., Castello, A., Mohammed, S., Moindrot, B., Nesterova, T.B., and Brockdorff, N. (2017). hnRNPK Recruits PCGF3/5-PRC1 to the Xist RNA B-Repeat to Establish Polycomb-Mediated Chromosomal Silencing. Mol Cell 68, 955–969 e910. 10.1016/j.molcel.2017.11.013.

4. Pandya-Jones, A., Markaki, Y., Serizay, J., Chitiashvili, T., Mancia Leon, W.R., Damianov, A., Chronis, C., Papp, B., Chen, C.K., McKee, R., et al. (2020). A protein assembly mediates Xist localization and gene silencing. Nature 587, 145–151. 10.1038/s41586-020-2703-0.

5. Lu, Z., Guo, J.K., Wei, Y., Dou, D.R., Zarnegar, B., Ma, Q., Li, R., Zhao, Y., Liu, F., Choudhry, H., et al. (2020). Structural modularity of the XIST ribonucleoprotein complex. Nat Commun 11, 6163. 10.1038/s41467-020-20040-3.

6. Graindorge, A., Pinheiro, I., Nawrocka, A., Mallory, A.C., Tsvetkov, P., Gil, N., Carolis, C., Buchholz, F., Ulitsky, I., Heard, E., et al. (2019). In-cell identification and measurement of RNA-protein interactions. Nat Commun 10, 5317. 10.1038/s41467-019-13235-w.

7. Weidmann, C.A., Mustoe, A.M., Jariwala, P.B., Calabrese, J.M., and Weeks, K.M. (2021). Analysis of RNA-protein networks with RNP-MaP defines functional hubs on RNA. Nat Biotechnol 39, 347–356. 10.1038/s41587-020-0709-7.

8. Minks, J., Baldry, S.E., Yang, C., Cotton, A.M., and Brown, C.J. (2013). XIST-induced silencing of flanking genes is achieved by additive action of repeat a monomers in human somatic cells. Epigenetics Chromatin 6, 23. 1756-8935-6-23 [pii] 10.1186/1756-8935-6-23.

9. Smola, M.J., Christy, T.W., Inoue, K., Nicholson, C.O., Friedersdorf, M., Keene, J.D., Lee, D.M., Calabrese, J.M., and Weeks, K.M. (2016). SHAPE reveals transcript-wide interactions, complex structural domains, and protein interactions across the Xist lncRNA in living cells. Proc Natl Acad Sci U S A 113, 10322–10327. 10.1073/pnas.1600008113.

10. Chu, C., Zhang, Q.C., da Rocha, S.T., Flynn, R.A., Bharadwaj, M., Calabrese, J.M., Magnuson, T., Heard, E., and Chang, H.Y. (2015). Systematic discovery of Xist RNA binding proteins. Cell 161, 404–416. 10.1016/j.cell.2015.03.025.

11. McHugh, C.A., Chen, C.K., Chow, A., Surka, C.F., Tran, C., McDonel, P., Pandya-Jones, A., Blanco, M., Burghard, C., Moradian, A., et al. (2015). The Xist lncRNA interacts directly with SHARP to silence transcription through HDAC3. Nature 521, 232–236. 10.1038/nature14443.

12. Monfort, A., Di Minin, G., Postlmayr, A., Freimann, R., Arieti, F., Thore, S., and Wutz, A. (2015). Identification of Spen as a Crucial Factor for Xist Function through Forward Genetic Screening in Haploid Embryonic Stem Cells. Cell Rep 12, 554–561. 10.1016/j.celrep.2015.06.067.

13. Moindrot, B., Cerase, A., Coker, H., Masui, O., Grijzenhout, A., Pintacuda, G., Schermelleh, L., Nesterova, T.B., and Brockdorff, N. (2015). A Pooled shRNA Screen Identifies Rbm15, Spen, and Wtap as Factors Required for Xist RNA-Mediated Silencing. Cell Rep 12, 562–572. 10.1016/j.celrep.2015.06.053.

14. Wutz, A., Rasmussen, T.P., and Jaenisch, R. (2002). Chromosomal silencing and localization are mediated by different domains of Xist RNA. Nat Genet 30, 167–174. 10.1038/ng820.

15. Duszczyk, M.M., Wutz, A., Rybin, V., and Sattler, M. (2011). The Xist RNA A-repeat comprises a novel AUCG tetraloop fold and a platform for multimerization. RNA 17, 1973–1982. 10.1261/rna.2747411.

16. Lu, Z., Zhang, Q.C., Lee, B., Flynn, R.A., Smith, M.A., Robinson, J.T., Davidovich, C., Gooding, A.R., Goodrich, K.J., Mattick, J.S., et al. (2016). RNA Duplex Map in Living Cells Reveals Higher-Order Transcriptome Structure. Cell 165, 1267–1279. 10.1016/j.cell.2016.04.028.

17. Carter, A.C., Xu, J., Nakamoto, M.Y., Wei, Y., Zarnegar, B.J., Shi, Q., Broughton, J.P., Ransom, R.C., Salhotra, A., Nagaraja, S.D., et al. (2020). Spen links RNA-mediated endogenous retrovirus silencing and X chromosome inactivation. Elife 9. 10.7554/eLife.54508.

18. Button, A.C., Hall, S.D., Ashley, E.L., and McHugh, C.A. (2024). Dissection of protein and RNA regions required for SPEN binding to XIST A-repeat RNA. RNA 30, 240–255. 10.1261/rna.079713.123.

19. Kaufmann, C., and Wutz, A. (2023). IndiSPENsable for X Chromosome Inactivation and Gene Silencing. Epigenomes 7. 10.3390/epigenomes7040028.

20. Dossin, F., Pinheiro, I., Zylicz, J.J., Roensch, J., Collombet, S., Le Saux, A., Chelmicki, T., Attia, M., Kapoor, V., Zhan, Y., et al. (2020). SPEN integrates transcriptional and epigenetic control of X-inactivation. Nature 578, 455–460. 10.1038/s41586-020-1974-9.

21. Zylicz, J.J., Bousard, A., Zumer, K., Dossin, F., Mohammad, E., da Rocha, S.T., Schwalb, B., Syx, L., Dingli, F., Loew, D., et al. (2019). The Implication of Early Chromatin Changes in X Chromosome Inactivation. Cell 176, 182–197 e123. 10.1016/j.cell.2018.11.041.

22. Dixon-McDougall, T., and Brown, C.J. (2022). Multiple distinct domains of human XIST are required to coordinate gene silencing and subsequent heterochromatin formation. Epigenetics Chromatin 15, 6. 10.1186/s13072-022-00438-7.

23. Wang, Y., Zhong, Y., Zhou, Y., Tanaseichuk, O., Li, Z., and Zhao, J.C. (2019). Identification of a Xist silencing domain by Tiling CRISPR. Sci Rep 9, 2408. 10.1038/s41598-018-36750-0.

24. Rodermund, L., Coker, H., Oldenkamp, R., Wei, G., Bowness, J., Rajkumar, B., Nesterova, T., Susano Pinto, D.M., Schermelleh, L., and Brockdorff, N. (2021). Time-resolved structured illumination microscopy reveals key principles of Xist RNA spreading. Science 372. 10.1126/science.abe7500.

25. Hoki, Y., Kimura, N., Kanbayashi, M., Amakawa, Y., Ohhata, T., Sasaki, H., and Sado, T. (2009). A proximal conserved repeat in the Xist gene is essential as a genomic element for X-inactivation in mouse. Development 136, 139–146. 10.1242/dev.026427.

26. Nesterova, T.B., Wei, G., Coker, H., Pintacuda, G., Bowness, J.S., Zhang, T., Almeida, M., Bloechl, B., Moindrot, B., Carter, E.J., et al. (2019). Systematic allelic analysis defines the interplay of key pathways in X chromosome inactivation. Nat Commun 10, 3129. 10.1038/s41467-019-11171-3.

27. Trotman, J.B., Lee, D.M., Cherney, R.E., Kim, S.O., Inoue, K., Schertzer, M.D., Bischoff, S.R., Cowley, D.O., and Calabrese, J.M. (2020). Elements at the 5’ end of Xist harbor SPEN-independent transcriptional antiterminator activity. Nucleic Acids Res 48, 10500–10517. 10.1093/nar/gkaa789.

28. Royce-Tolland, M.E., Andersen, A.A., Koyfman, H.R., Talbot, D.J., Wutz, A., Tonks, I.D., Kay, G.F., and Panning, B. (2010). The A-repeat links ASF/SF2-dependent Xist RNA processing with random choice during X inactivation. Nat Struct Mol Biol 17, 948–954. 10.1038/nsmb.1877.

29. Colognori, D., Sunwoo, H., Kriz, A.J., Wang, C.Y., and Lee, J.T. (2019). Xist Deletional Analysis Reveals an Interdependency between Xist RNA and Polycomb Complexes for Spreading along the Inactive X. Mol Cell 74, 101–117 e110. 10.1016/j.molcel.2019.01.015.

30. Sakata, Y., Nagao, K., Hoki, Y., Sasaki, H., Obuse, C., and Sado, T. (2017). Defects in dosage compensation impact global gene regulation in the mouse trophoblast. Development 144, 2784–2797. 10.1242/dev.149138.

31. Cartegni, L., Wang, J., Zhu, Z., Zhang, M.Q., and Krainer, A.R. (2003). ESEfinder: A web resource to identify exonic splicing enhancers. Nucleic Acids Res 31, 3568–3571. 10.1093/nar/gkg616.

32. Radio, F.C., Pang, K., Ciolfi, A., Levy, M.A., Hernandez-Garcia, A., Pedace, L., Pantaleoni, F., Liu, Z., de Boer, E., Jackson, A., et al. (2021). SPEN haploinsufficiency causes a neurodevelopmental disorder overlapping proximal 1p36 deletion syndrome with an episignature of X chromosomes in females. Am J Hum Genet 108, 502–516. 10.1016/j.ajhg.2021.01.015.

33. Damrauer, J.S., Beckabir, W., Klomp, J., Zhou, M., Plimack, E.R., Galsky, M.D., Grivas, P., Hahn, N.M., O’Donnell, P.H., Iyer, G., et al. (2022). Collaborative study from the Bladder Cancer Advocacy Network for the genomic analysis of metastatic urothelial cancer. Nature communications 13, 6658. 10.1038/s41467-022-33980-9.

34. Heath, A.P., Ferretti, V., Agrawal, S., An, M., Angelakos, J.C., Arya, R., Bajari, R., Baqar, B., Barnowski, J.H.B., Burt, J., et al. (2021). The NCI Genomic Data Commons. Nat Genet 53, 257–262. 10.1038/s41588-021-00791-5.

35. Mendiratta, G., Ke, E., Aziz, M., Liarakos, D., Tong, M., and Stites, E.C. (2021). Cancer gene mutation frequencies for the U.S. population. Nat Commun 12, 5961. 10.1038/s41467-021-26213-y.

36. Reddy, A., Zhang, J., Davis, N.S., Moffitt, A.B., Love, C.L., Waldrop, A., Leppa, S., Pasanen, A., Meriranta, L., Karjalainen-Lindsberg, M.L., et al. (2017). Genetic and Functional Drivers of Diffuse Large B Cell Lymphoma. Cell 171, 481–494 e415. 10.1016/j.cell.2017.09.027.

37. Sarkozy, C., Hung, S.S., Chavez, E.A., Duns, G., Takata, K., Chong, L.C., Aoki, T., Jiang, A., Miyata-Takata, T., Telenius, A., et al. (2021). Mutational landscape of gray zone lymphoma. Blood 137, 1765–1776. 10.1182/blood.2020007507.

38. Schmitz, R., Wright, G.W., Huang, D.W., Johnson, C.A., Phelan, J.D., Wang, J.Q., Roulland, S., Kasbekar, M., Young, R.M., Shaffer, A.L., et al. (2018). Genetics and Pathogenesis of Diffuse Large B-Cell Lymphoma. N Engl J Med 378, 1396–1407. 10.1056/NEJMoa1801445.

39. Seiler, M., Peng, S., Agrawal, A.A., Palacino, J., Teng, T., Zhu, P., Smith, P.G., Cancer Genome Atlas Research, N., Buonamici, S., and Yu, L. (2018). Somatic Mutational Landscape of Splicing Factor Genes and Their Functional Consequences across 33 Cancer Types. Cell Rep 23, 282–296 e284. 10.1016/j.celrep.2018.01.088.

40. Tyner, J.W., Tognon, C.E., Bottomly, D., Wilmot, B., Kurtz, S.E., Savage, S.L., Long, N., Schultz, A.R., Traer, E., Abel, M., et al. (2018). Functional genomic landscape of acute myeloid leukaemia. Nature 562, 526–531. 10.1038/s41586-018-0623-z.

41. Legare, S., Cavallone, L., Mamo, A., Chabot, C., Sirois, I., Magliocco, A., Klimowicz, A., Tonin, P.N., Buchanan, M., Keilty, D., et al. (2015). The Estrogen Receptor Cofactor SPEN Functions as a Tumor Suppressor and Candidate Biomarker of Drug Responsiveness in Hormone-Dependent Breast Cancers. Cancer Res 75, 4351–4363. 10.1158/0008-5472.CAN-14-3475.

42. Chen, S., Francioli, L.C., Goodrich, J.K., Collins, R.L., Kanai, M., Wang, Q., Alfoldi, J., Watts, N.A., Vittal, C., Gauthier, L.D., et al. (2024). A genomic mutational constraint map using variation in 76,156 human genomes. Nature 625, 92–100. 10.1038/s41586-023-06045-0.

43. Trotman, J.B., Abrash, E.W., Murvin, M.M., Braceros, A.K., Li, S., Boyson, S.P., Salcido, R.T., Cherney, R.E., Bischoff, S.R., Kaufmann, K., et al. (2025). Isogenic comparison of Airn and Xist reveals core principles of Polycomb recruitment by lncRNAs. Mol Cell 85, 1117–1133 e1114. 10.1016/j.molcel.2025.02.014.

44. Kirk, J.M., Kim, S.O., Inoue, K., Smola, M.J., Lee, D.M., Schertzer, M.D., Wooten, J.S., Baker, A.R., Sprague, D., Collins, D.W., et al. (2018). Functional classification of long non-coding RNAs by k-mer content. Nat Genet 50, 1474–1482. 10.1038/s41588-018-0207-8.

45. Hughes, A.L., Szczurek, A.T., Kelley, J.R., Lastuvkova, A., Turberfield, A.H., Dimitrova, E., Blackledge, N.P., and Klose, R.J. (2023). A CpG island-encoded mechanism protects genes from premature transcription termination. Nature communications 14, 726. 10.1038/s41467-023-36236-2.

46. Koerner, M.V., Pauler, F.M., Hudson, Q.J., Santoro, F., Sawicka, A., Guenzl, P.M., Stricker, S.H., Schichl, Y.M., Latos, P.A., Klement, R.M., et al. (2012). A downstream CpG island controls transcript initiation and elongation and the methylation state of the imprinted Airn macro ncRNA promoter. PLoS Genet 8, e1002540. 10.1371/journal.pgen.1002540.

47. Younis, I., Berg, M., Kaida, D., Dittmar, K., Wang, C., and Dreyfuss, G. (2010). Rapid-response splicing reporter screens identify differential regulators of constitutive and alternative splicing. Mol Cell Biol 30, 1718–1728. 10.1128/MCB.01301-09.

48. Matsuura, R., Nakajima, T., Ichihara, S., and Sado, T. (2021). Ectopic Splicing Disturbs the Function of Xist RNA to Establish the Stable Heterochromatin State. Front Cell Dev Biol 9, 751154. 10.3389/fcell.2021.751154.

49. Love, M.I., Huber, W., and Anders, S. (2014). Moderated estimation of fold change and dispersion for RNA-seq data with DESeq2. Genome Biol 15, 550. 10.1186/s13059-014-0550-8.

50. Colognori, D., Sunwoo, H., Wang, D., Wang, C.Y., and Lee, J.T. (2020). Xist Repeats A and B Account for Two Distinct Phases of X Inactivation Establishment. Dev Cell 54, 21–32 e25. 10.1016/j.devcel.2020.05.021.

51. Schertzer, M.D., Braceros, K.C.A., Starmer, J., Cherney, R.E., Lee, D.M., Salazar, G., Justice, M., Bischoff, S.R., Cowley, D.O., Ariel, P., et al. (2019). lncRNA-Induced Spread of Polycomb Controlled by Genome Architecture, RNA Abundance, and CpG Island DNA. Mol Cell 75, 523–537 e510. 10.1016/j.molcel.2019.05.028.

52. Jachowicz, J.W., Strehle, M., Banerjee, A.K., Blanco, M.R., Thai, J., and Guttman, M. (2022). Xist spatially amplifies SHARP/SPEN recruitment to balance chromosome-wide silencing and specificity to the X chromosome. Nat Struct Mol Biol 29, 239–249. 10.1038/s41594-022-00739-1.

53. Engreitz, J.M., Pandya-Jones, A., McDonel, P., Shishkin, A., Sirokman, K., Surka, C., Kadri, S., Xing, J., Goren, A., Lander, E.S., et al. (2013). The Xist lncRNA exploits three-dimensional genome architecture to spread across the X chromosome. Science 341, 1237973. science.1237973 [pii] 10.1126/science.1237973.

54. Bailey, T.L. (2021). STREME: Accurate and versatile sequence motif discovery. Bioinformatics. 10.1093/bioinformatics/btab203.

55. Hinkle, E.R., Wiedner, H.J., Torres, E.V., Jackson, M., Black, A.J., Blue, R.E., Harris, S.E., Guzman, B.B., Gentile, G.M., Lee, E.Y., et al. (2022). Alternative splicing regulation of membrane trafficking genes during myogenesis. RNA 28, 523–540. 10.1261/rna.078993.121.

56. Aguilar, R., Spencer, K.B., Kesner, B., Rizvi, N.F., Badmalia, M.D., Mrozowich, T., Mortison, J.D., Rivera, C., Smith, G.F., Burchard, J., et al. (2022). Targeting Xist with compounds that disrupt RNA structure and X inactivation. Nature 604, 160–166. 10.1038/s41586-022-04537-z.

57. Cascarina, S.M., and Ross, E.D. (2022). Expansion and functional analysis of the SR-related protein family across the domains of life. RNA 28, 1298–1314. 10.1261/rna.079170.122.

58. Sliskovic, I., Eich, H., and Muller-McNicoll, M. (2022). Exploring the multifunctionality of SR proteins. Biochemical Society transactions 50, 187–198. 10.1042/BST20210325.

59. Malovannaya, A., Lanz, R.B., Jung, S.Y., Bulynko, Y., Le, N.T., Chan, D.W., Ding, C., Shi, Y., Yucer, N., Krenciute, G., et al. (2011). Analysis of the human endogenous coregulator complexome. Cell 145, 787–799. 10.1016/j.cell.2011.05.006.

60. Hein, M.Y., Hubner, N.C., Poser, I., Cox, J., Nagaraj, N., Toyoda, Y., Gak, I.A., Weisswange, I., Mansfeld, J., Buchholz, F., et al. (2015). A human interactome in three quantitative dimensions organized by stoichiometries and abundances. Cell 163, 712–723. 10.1016/j.cell.2015.09.053.

61. Potenza, E., Di Domenico, T., Walsh, I., and Tosatto, S.C. (2015). MobiDB 2.0: an improved database of intrinsically disordered and mobile proteins. Nucleic Acids Res 43, D315–320. 10.1093/nar/gku982.

62. Manley, J.L., and Krainer, A.R. (2010). A rational nomenclature for serine/arginine-rich protein splicing factors (SR proteins). Genes Dev 24, 1073–1074. 10.1101/gad.1934910.

63. Cazalla, D., Zhu, J., Manche, L., Huber, E., Krainer, A.R., and Caceres, J.F. (2002). Nuclear export and retention signals in the RS domain of SR proteins. Mol Cell Biol 22, 6871–6882. 10.1128/MCB.22.19.6871-6882.2002.

64. Markaki, Y., Gan Chong, J., Wang, Y., Jacobson, E.C., Luong, C., Tan, S.Y.X., Jachowicz, J.W., Strehle, M., Maestrini, D., Banerjee, A.K., et al. (2021). Xist nucleates local protein gradients to propagate silencing across the X chromosome. Cell 184, 6174–6192 e6132. 10.1016/j.cell.2021.10.022.

65. Van Nostrand, E.L., Pratt, G.A., Shishkin, A.A., Gelboin-Burkhart, C., Fang, M.Y., Sundararaman, B., Blue, S.M., Nguyen, T.B., Surka, C., Elkins, K., et al. (2016). Robust transcriptome-wide discovery of RNA-binding protein binding sites with enhanced CLIP (eCLIP). Nat Methods 13, 508–514. 10.1038/nmeth.3810.

66. Consortium, G. (2013). The Genotype-Tissue Expression (GTEx) project. Nat Genet 45, 580–585. 10.1038/ng.2653.

67. van Dam, S., Craig, T., and de Magalhaes, J.P. (2015). GeneFriends: a human RNA-seq-based gene and transcript co-expression database. Nucleic Acids Res 43, D1124–1132. 10.1093/nar/gku1042.

68. Stuart, J.M., Segal, E., Koller, D., and Kim, S.K. (2003). A gene-coexpression network for global discovery of conserved genetic modules. Science 302, 249–255. 10.1126/science.1087447.

69. Quackenbush, J. (2003). Genomics. Microarrays--guilt by association. Science 302, 240–241. 10.1126/science.1090887.

70. Yuan, Z., VanderWielen, B.D., Giaimo, B.D., Pan, L., Collins, C.E., Turkiewicz, A., Hein, K., Oswald, F., Borggrefe, T., and Kovall, R.A. (2019). Structural and Functional Studies of the RBPJ-SHARP Complex Reveal a Conserved Corepressor Binding Site. Cell Rep 26, 845–854 e846. 10.1016/j.celrep.2018.12.097.

71. Oswald, F., Rodriguez, P., Giaimo, B.D., Antonello, Z.A., Mira, L., Mittler, G., Thiel, V.N., Collins, K.J., Tabaja, N., Cizelsky, W., et al. (2016). A phospho-dependent mechanism involving NCoR and KMT2D controls a permissive chromatin state at Notch target genes. Nucleic Acids Res 44, 4703–4720. 10.1093/nar/gkw105.

72. Gregersen, L.H., Mitter, R., Ugalde, A.P., Nojima, T., Proudfoot, N.J., Agami, R., Stewart, A., and Svejstrup, J.Q. (2019). SCAF4 and SCAF8, mRNA Anti-Terminator Proteins. Cell 177, 1797–1813 e1718. 10.1016/j.cell.2019.04.038.

73. Altendorfer, E., Mochalova, Y., and Mayer, A. (2022). BRD4: a general regulator of transcription elongation. Transcription 13, 70–81. 10.1080/21541264.2022.2108302.

74. Yu, B., Qi, Y., Li, R., Shi, Q., Satpathy, A.T., and Chang, H.Y. (2021). B cell-specific XIST complex enforces X-inactivation and restrains atypical B cells. Cell 184, 1790–1803 e1717. 10.1016/j.cell.2021.02.015.

75. Andersen, P.R., Domanski, M., Kristiansen, M.S., Storvall, H., Ntini, E., Verheggen, C., Schein, A., Bunkenborg, J., Poser, I., Hallais, M., et al. (2013). The human cap-binding complex is functionally connected to the nuclear RNA exosome. Nat Struct Mol Biol 20, 1367–1376. 10.1038/nsmb.2703.

76. Polak, P., Garland, W., Rathore, O., Schmid, M., Salerno-Kochan, A., Jakobsen, L., Gockert, M., Gerlach, P., Silla, T., Andersen, J.S., et al. (2023). Dual agonistic and antagonistic roles of ZC3H18 provide for co-activation of distinct nuclear RNA decay pathways. Cell Rep 42, 113325. 10.1016/j.celrep.2023.113325.

77. Liu, J., Dou, X., Chen, C., Chen, C., Liu, C., Xu, M.M., Zhao, S., Shen, B., Gao, Y., Han, D., and He, C. (2020). N (6)-methyladenosine of chromosome-associated regulatory RNA regulates chromatin state and transcription. Science 367, 580–586. 10.1126/science.aay6018.

78. Patil, D.P., Chen, C.K., Pickering, B.F., Chow, A., Jackson, C., Guttman, M., and Jaffrey, S.R. (2016). m(6)A RNA methylation promotes XIST-mediated transcriptional repression. Nature 537, 369-+. 10.1038/nature19342.

79. Xiao, W., Adhikari, S., Dahal, U., Chen, Y.S., Hao, Y.J., Sun, B.F., Sun, H.Y., Li, A., Ping, X.L., Lai, W.Y., et al. (2016). Nuclear m(6)A Reader YTHDC1 Regulates mRNA Splicing. Mol Cell 61, 507–519. 10.1016/j.molcel.2016.01.012.

80. Nayler, O., Hartmann, A.M., and Stamm, S. (2000). The ER repeat protein YT521-B localizes to a novel subnuclear compartment. J Cell Biol 150, 949–962. 10.1083/jcb.150.5.949.

81. Bousard, A., Raposo, A.C., Zylicz, J.J., Picard, C., Pires, V.B., Qi, Y., Gil, C., Syx, L., Chang, H.Y., Heard, E., and da Rocha, S.T. (2019). The role of Xist-mediated Polycomb recruitment in the initiation of X-chromosome inactivation. EMBO Rep 20, e48019. 10.15252/embr.201948019.

82. Pandit, S., Zhou, Y., Shiue, L., Coutinho-Mansfield, G., Li, H., Qiu, J., Huang, J., Yeo, G.W., Ares, M., Jr., and Fu, X.D. (2013). Genome-wide analysis reveals SR protein cooperation and competition in regulated splicing. Mol Cell 50, 223–235. 10.1016/j.molcel.2013.03.001.

83. Feng, H., Bao, S., Rahman, M.A., Weyn-Vanhentenryck, S.M., Khan, A., Wong, J., Shah, A., Flynn, E.D., Krainer, A.R., and Zhang, C. (2019). Modeling RNA-Binding Protein Specificity In Vivo by Precisely Registering Protein-RNA Crosslink Sites. Mol Cell 74, 1189–1204 e1186. 10.1016/j.molcel.2019.02.002.

84. Segovia, D., Adams, D.W., Hoffman, N., Safaric Tepes, P., Wee, T.L., Cifani, P., Joshua-Tor, L., and Krainer, A.R. (2024). SRSF1 interactome determined by proximity labeling reveals direct interaction with spliceosomal RNA helicase DDX23. Proc Natl Acad Sci U S A 121, e2322974121. 10.1073/pnas.2322974121.

85. Daubner, G.M., Clery, A., Jayne, S., Stevenin, J., and Allain, F.H. (2012). A syn-anti conformational difference allows SRSF2 to recognize guanines and cytosines equally well. EMBO J 31, 162–174. 10.1038/emboj.2011.367.

86. Wheeler, E.C., Vora, S., Mayer, D., Kotini, A.G., Olszewska, M., Park, S.S., Guccione, E., Teruya-Feldstein, J., Silverman, L., Sunahara, R.K., et al. (2022). Integrative RNA-omics Discovers GNAS Alternative Splicing as a Phenotypic Driver of Splicing Factor-Mutant Neoplasms. Cancer Discov 12, 836–855. 10.1158/2159-8290.CD-21-0508.

87. Zhang, X., Smits, A.H., van Tilburg, G.B., Jansen, P.W., Makowski, M.M., Ovaa, H., and Vermeulen, M. (2017). An Interaction Landscape of Ubiquitin Signaling. Mol Cell 65, 941–955 e948. 10.1016/j.molcel.2017.01.004.

88. Nabet, B., Ferguson, F.M., Seong, B.K.A., Kuljanin, M., Leggett, A.L., Mohardt, M.L., Robichaud, A., Conway, A.S., Buckley, D.L., Mancias, J.D., et al. (2020). Rapid and direct control of target protein levels with VHL-recruiting dTAG molecules. Nat Commun 11, 4687. 10.1038/s41467-020-18377-w.

89. Nabet, B., Roberts, J.M., Buckley, D.L., Paulk, J., Dastjerdi, S., Yang, A., Leggett, A.L., Erb, M.A., Lawlor, M.A., Souza, A., et al. (2018). The dTAG system for immediate and target-specific protein degradation. Nat Chem Biol 14, 431–441. 10.1038/s41589-018-0021-8.

90. Abuhashem, A., and Hadjantonakis, A.K. (2022). Generation of knock-in degron tags for endogenous proteins in mice using the dTAG system. STAR Protoc 3, 101660. 10.1016/j.xpro.2022.101660.

91. Chen, C., Liu, W., Guo, J., Liu, Y., Liu, X., Liu, J., Dou, X., Le, R., Huang, Y., Li, C., et al. (2021). Nuclear m(6)A reader YTHDC1 regulates the scaffold function of LINE1 RNA in mouse ESCs and early embryos. Protein Cell 12, 455–474. 10.1007/s13238-021-00837-8.

92. Xu, W., Li, J., He, C., Wen, J., Ma, H., Rong, B., Diao, J., Wang, L., Wang, J., Wu, F., et al. (2021). METTL3 regulates heterochromatin in mouse embryonic stem cells. Nature 591, 317–321. 10.1038/s41586-021-03210-1.

93. Ji, X., Zhou, Y., Pandit, S., Huang, J., Li, H., Lin, C.Y., Xiao, R., Burge, C.B., and Fu, X.D. (2013). SR proteins collaborate with 7SK and promoter-associated nascent RNA to release paused polymerase. Cell 153, 855–868. 10.1016/j.cell.2013.04.028.

94. Guo, Y.E., Manteiga, J.C., Henninger, J.E., Sabari, B.R., Dall’Agnese, A., Hannett, N.M., Spille, J.H., Afeyan, L.K., Zamudio, A.V., Shrinivas, K., et al. (2019). Pol II phosphorylation regulates a switch between transcriptional and splicing condensates. Nature 572, 543–548. 10.1038/s41586-019-1464-0.

95. Das, R., Yu, J., Zhang, Z., Gygi, M.P., Krainer, A.R., Gygi, S.P., and Reed, R. (2007). SR proteins function in coupling RNAP II transcription to pre-mRNA splicing. Mol Cell 26, 867–881. 10.1016/j.molcel.2007.05.036.

96. Horiuchi, K., Kawamura, T., Iwanari, H., Ohashi, R., Naito, M., Kodama, T., and Hamakubo, T. (2013). Identification of Wilms’ tumor 1-associating protein complex and its role in alternative splicing and the cell cycle. J Biol Chem 288, 33292–33302. 10.1074/jbc.M113.500397.

97. Tsue, A.F., Kania, E.E., Lei, D.Q., Fields, R., McGann, C.D., Marciniak, D.M., Hershberg, E.A., Deng, X., Kihiu, M., Ong, S.E., et al. (2024). Multiomic characterization of RNA microenvironments by oligonucleotide-mediated proximity-interactome mapping. Nat Methods 21, 2058–2071. 10.1038/s41592-024-02457-6.

98. Huttlin, E.L., Ting, L., Bruckner, R.J., Gebreab, F., Gygi, M.P., Szpyt, J., Tam, S., Zarraga, G., Colby, G., Baltier, K., et al. (2015). The BioPlex Network: A Systematic Exploration of the Human Interactome. Cell 162, 425–440. 10.1016/j.cell.2015.06.043.

99. Oswald, F., Kostezka, U., Astrahantseff, K., Bourteele, S., Dillinger, K., Zechner, U., Ludwig, L., Wilda, M., Hameister, H., Knochel, W., et al. (2002). SHARP is a novel component of the Notch/RBP-Jkappa signalling pathway. EMBO J 21, 5417–5426. 10.1093/emboj/cdf549.

100. Trotman, J.B., Braceros, K.C.A., Cherney, R.E., Murvin, M.M., and Calabrese, J.M. (2021). The control of polycomb repressive complexes by long noncoding RNAs. Wiley Interdiscip Rev RNA 12, e1657. 10.1002/wrna.1657.

101. Kuroda, K., Han, H., Tani, S., Tanigaki, K., Tun, T., Furukawa, T., Taniguchi, Y., Kurooka, H., Hamada, Y., Toyokuni, S., and Honjo, T. (2003). Regulation of marginal zone B cell development by MINT, a suppressor of Notch/RBP-J signaling pathway. Immunity 18, 301–312. 10.1016/s1074-7613(03)00029-3.

102. Tsuji, M., Shinkura, R., Kuroda, K., Yabe, D., and Honjo, T. (2007). Msx2-interacting nuclear target protein (Mint) deficiency reveals negative regulation of early thymocyte differentiation by Notch/RBP-J signaling. Proc Natl Acad Sci U S A 104, 1610–1615. 10.1073/pnas.0610520104.

103. Palozola, K.C., Donahue, G., Liu, H., Grant, G.R., Becker, J.S., Cote, A., Yu, H., Raj, A., and Zaret, K.S. (2017). Mitotic transcription and waves of gene reactivation during mitotic exit. Science 358, 119–122. 10.1126/science.aal4671.

104. Palozola, K.C., Donahue, G., and Zaret, K.S. (2021). EU-RNA-seq for in vivo labeling and high throughput sequencing of nascent transcripts. STAR Protoc 2, 100651. 10.1016/j.xpro.2021.100651.

105. Wutz, A., Kaufmann, K., and Sting, S. (2025). A comprehensive CRISPR/Cas9-targeted base mutagenesis identifies an amino acid substitution separating functions of SPEN in X chromosome inactivation from development. Research Square. 10.21203/rs.3.rs-6717927/v1.

106. Cho, N.H., Cheveralls, K.C., Brunner, A.D., Kim, K., Michaelis, A.C., Raghavan, P., Kobayashi, H., Savy, L., Li, J.Y., Canaj, H., et al. (2022). OpenCell: Endogenous tagging for the cartography of human cellular organization. Science 375, eabi6983. 10.1126/science.abi6983.

107. Appel, L.M., Franke, V., Benedum, J., Grishkovskaya, I., Strobl, X., Polyansky, A., Ammann, G., Platzer, S., Neudolt, A., Wunder, A., et al. (2023). The SPOC domain is a phosphoserine binding module that bridges transcription machinery with co- and post-transcriptional regulators. Nature communications 14, 166. 10.1038/s41467-023-35853-1.

108. Zhang, L., Zhang, Y., Chen, Y., Gholamalamdari, O., Wang, Y., Ma, J., and Belmont, A.S. (2020). TSA-seq reveals a largely conserved genome organization relative to nuclear speckles with small position changes tightly correlated with gene expression changes. Genome Res 31, 251–264. 10.1101/gr.266239.120.

109. Bhat, P., Chow, A., Emert, B., Ettlin, O., Quinodoz, S.A., Strehle, M., Takei, Y., Burr, A., Goronzy, I.N., Chen, A.W., et al. (2024). Genome organization around nuclear speckles drives mRNA splicing efficiency. Nature 629, 1165–1173. 10.1038/s41586-024-07429-6.

110. Shopland, L.S., Johnson, C.V., Byron, M., McNeil, J., and Lawrence, J.B. (2003). Clustering of multiple specific genes and gene-rich R-bands around SC-35 domains: evidence for local euchromatic neighborhoods. J Cell Biol 162, 981–990. 10.1083/jcb.200303131.

111. Yoon, Y., Bournique, E., Soles, L.V., Yin, H., Chu, H.F., Yin, C., Zhuang, Y., Liu, X., Liu, L., Jeong, J., et al. (2025). RBBP6 anchors pre-mRNA 3’ end processing to nuclear speckles for efficient gene expression. Mol Cell 85, 555–570 e558. 10.1016/j.molcel.2024.12.016.

112. Collombet, S., Rall, I., Dugast-Darzacq, C., Heckert, A., Halavatyi, A., Le Saux, A., Dailey, G., Darzacq, X., and Heard, E. (2023). RNA polymerase II depletion from the inactive X chromosome territory is not mediated by physical compartmentalization. Nat Struct Mol Biol 30, 1216–1223. 10.1038/s41594-023-01008-5.

113. Rouviere, J.O., Salerno-Kochan, A., Lykke-Andersen, S., Garland, W., Dou, Y., Rathore, O., Molska, E.S., Wu, G., Schmid, M., Bugai, A., et al. (2023). ARS2 instructs early transcription termination-coupled RNA decay by recruiting ZC3H4 to nascent transcripts. Mol Cell 83, 2240–2257 e2246. 10.1016/j.molcel.2023.05.028.

114. Parker, M.T., Knop, K., Zacharaki, V., Sherwood, A.V., Tome, D., Yu, X., Martin, P.G., Beynon, J., Michaels, S.D., Barton, G.J., and Simpson, G.G. (2021). Widespread premature transcription termination of Arabidopsis thaliana NLR genes by the spen protein FPA. eLife 10. 10.7554/eLife.65537.

115. Hornyik, C., Terzi, L.C., and Simpson, G.G. (2010). The spen family protein FPA controls alternative cleavage and polyadenylation of RNA. Dev Cell 18, 203–213. 10.1016/j.devcel.2009.12.009.

116. Baurle, I., Smith, L., Baulcombe, D.C., and Dean, C. (2007). Widespread role for the flowering-time regulators FCA and FPA in RNA-mediated chromatin silencing. Science 318, 109–112. 10.1126/science.1146565.

117. Busch, A., and Hertel, K.J. (2012). Evolution of SR protein and hnRNP splicing regulatory factors. Wiley Interdiscip Rev RNA 3, 1–12. 10.1002/wrna.100.

118. Duret, L., Chureau, C., Samain, S., Weissenbach, J., and Avner, P. (2006). The Xist RNA gene evolved in eutherians by pseudogenization of a protein-coding gene. Science 312, 1653–1655. 10.1126/science.1126316.

119. Elisaphenko, E.A., Kolesnikov, N.N., Shevchenko, A.I., Rogozin, I.B., Nesterova, T.B., Brockdorff, N., and Zakian, S.M. (2008). A dual origin of the Xist gene from a protein-coding gene and a set of transposable elements. PLoS One 3, e2521. 10.1371/journal.pone.0002521.

120. McCarthy, R.L., Kaeding, K.E., Keller, S.H., Zhong, Y., Xu, L., Hsieh, A., Hou, Y., Donahue, G., Becker, J.S., Alberto, O., et al. (2021). Diverse heterochromatin-associated proteins repress distinct classes of genes and repetitive elements. Nat Cell Biol 23, 905–914. 10.1038/s41556-021-00725-7.

121. Tatarakis, A., Saini, H., Yu, J., Feng, W., Pinzon-Arteaga, C.A., and Moazed, D. (2025). Requirements for establishment and epigenetic stability of mammalian heterochromatin. Mol Cell 85, 3388–3406 e3312. 10.1016/j.molcel.2025.08.025.

122. Becker, J.S., McCarthy, R.L., Sidoli, S., Donahue, G., Kaeding, K.E., He, Z., Lin, S., Garcia, B.A., and Zaret, K.S. (2017). Genomic and Proteomic Resolution of Heterochromatin and Its Restriction of Alternate Fate Genes. Mol Cell 68, 1023–1037 e1015. 10.1016/j.molcel.2017.11.030.

123. Oswald, F., Winkler, M., Cao, Y., Astrahantseff, K., Bourteele, S., Knochel, W., and Borggrefe, T. (2005). RBP-Jkappa/SHARP recruits CtIP/CtBP corepressors to silence Notch target genes. Mol Cell Biol 25, 10379–10390. 10.1128/MCB.25.23.10379-10390.2005.

124. Shi, Y., Downes, M., Xie, W., Kao, H.Y., Ordentlich, P., Tsai, C.C., Hon, M., and Evans, R.M. (2001). Sharp, an inducible cofactor that integrates nuclear receptor repression and activation. Genes Dev 15, 1140–1151. 10.1101/gad.871201.

125. Ariyoshi, M., and Schwabe, J.W. (2003). A conserved structural motif reveals the essential transcriptional repression function of Spen proteins and their role in developmental signaling. Genes Dev 17, 1909–1920. 10.1101/gad.266203.

126. Chang, J.L., Lin, H.V., Blauwkamp, T.A., and Cadigan, K.M. (2008). Spenito and Split ends act redundantly to promote Wingless signaling. Dev Biol 314, 100–111. 10.1016/j.ydbio.2007.11.023.

127. Chen, F., and Rebay, I. (2000). split ends, a new component of the Drosophila EGF receptor pathway, regulates development of midline glial cells. Curr Biol 10, 943–946. 10.1016/s0960-9822(00)00625-4.

128. Doroquez, D.B., Orr-Weaver, T.L., and Rebay, I. (2007). Split ends antagonizes the Notch and potentiates the EGFR signaling pathways during Drosophila eye development. Mech Dev 124, 792–806. 10.1016/j.mod.2007.05.002.

129. Feng, Y., Bommer, G.T., Zhai, Y., Akyol, A., Hinoi, T., Winer, I., Lin, H.V., Cadigan, K.M., Cho, K.R., and Fearon, E.R. (2007). Drosophila split ends homologue SHARP functions as a positive regulator of Wnt/beta-catenin/T-cell factor signaling in neoplastic transformation. Cancer Res 67, 482–491. 10.1158/0008-5472.CAN-06-2314.

130. Kuang, B., Wu, S.C., Shin, Y., Luo, L., and Kolodziej, P. (2000). split ends encodes large nuclear proteins that regulate neuronal cell fate and axon extension in the Drosophila embryo. Development 127, 1517–1529. 10.1242/dev.127.7.1517.

131. Piovesan, D., Del Conte, A., Mehdiabadi, M., Aspromonte, M.C., Blum, M., Tesei, G., von Bulow, S., Lindorff-Larsen, K., and Tosatto, S.C.E. (2025). MOBIDB in 2025: integrating ensemble properties and function annotations for intrinsically disordered proteins. Nucleic Acids Res 53, D495–D503. 10.1093/nar/gkae969.

132. Johnson, P.Z., Kasprzak, W.K., Shapiro, B.A., and Simon, A.E. (2019). RNA2Drawer: geometrically strict drawing of nucleic acid structures with graphical structure editing and highlighting of complementary subsequences. RNA Biol 16, 1667–1671. 10.1080/15476286.2019.1659081.

133. Wu, B., Eliscovich, C., Yoon, Y.J., and Singer, R.H. (2016). Translation dynamics of single mRNAs in live cells and neurons. Science 352, 1430–1435. 10.1126/science.aaf1084.

134. Huang, X., Zheng, M., Wang, P., Mok, B.W., Liu, S., Lau, S.Y., Chen, P., Liu, Y.C., Liu, H., Chen, Y., et al. (2017). An NS-segment exonic splicing enhancer regulates influenza A virus replication in mammalian cells. Nature communications 8, 14751. 10.1038/ncomms14751.

135. Cherney, R.E., Eberhard, Q.E., Giri, G., Mills, C.A., Porrello, A., Zhang, Z., White, D., Trotman, J.B., Herring, L.E., Dominguez, D., and Calabrese, J.M. (2023). SAFB associates with nascent RNAs and can promote gene expression in mouse embryonic stem cells. RNA 29, 1535–1556. 10.1261/rna.079569.122.

136. Ran, F.A., Hsu, P.D., Wright, J., Agarwala, V., Scott, D.A., and Zhang, F. (2013). Genome engineering using the CRISPR-Cas9 system. Nat Protoc 8, 2281–2308. 10.1038/nprot.2013.143.

137. Grunwald, D., and Singer, R.H. (2010). In vivo imaging of labelled endogenous beta-actin mRNA during nucleocytoplasmic transport. Nature 467, 604–607. 10.1038/nature09438.

138. Schertzer, M.D., Thulson, E., Braceros, K.C.A., Lee, D.M., Hinkle, E.R., Murphy, R.M., Kim, S.O., Vitucci, E.C.M., and Calabrese, J.M. (2019). A piggyBac-based toolkit for inducible genome editing in mammalian cells. RNA 25, 1047–1058. 10.1261/rna.068932.118.

139. Cong, L., Ran, F.A., Cox, D., Lin, S., Barretto, R., Habib, N., Hsu, P.D., Wu, X., Jiang, W., Marraffini, L.A., and Zhang, F. (2013). Multiplex genome engineering using CRISPR/Cas systems. Science 339, 819–823. 10.1126/science.1231143.

140. Braceros, A.K., Schertzer, M.D., Omer, A., Trotman, J.B., Davis, E.S., Dowen, J.M., Phanstiel, D.H., Aiden, E.L., and Calabrese, J.M. (2023). Proximity-dependent recruitment of Polycomb repressive complexes by the lncRNA Airn. Cell Rep 42, 112803. 10.1016/j.celrep.2023.112803.

141. Dobin, A., Davis, C.A., Schlesinger, F., Drenkow, J., Zaleski, C., Jha, S., Batut, P., Chaisson, M., and Gingeras, T.R. (2013). STAR: ultrafast universal RNA-seq aligner. Bioinformatics 29, 15–21. 10.1093/bioinformatics/bts635.

142. Mudge, J.M., Carbonell-Sala, S., Diekhans, M., Martinez, J.G., Hunt, T., Jungreis, I., Loveland, J.E., Arnan, C., Barnes, I., Bennett, R., et al. (2025). GENCODE 2025: reference gene annotation for human and mouse. Nucleic Acids Res 53, D966–D975. 10.1093/nar/gkae1078.

143. Liao, Y., Smyth, G.K., and Shi, W. (2014). featureCounts: an efficient general purpose program for assigning sequence reads to genomic features. Bioinformatics 30, 923–930. 10.1093/bioinformatics/btt656.

144. Keane, T.M., Goodstadt, L., Danecek, P., White, M.A., Wong, K., Yalcin, B., Heger, A., Agam, A., Slater, G., Goodson, M., et al. (2011). Mouse genomic variation and its effect on phenotypes and gene regulation. Nature 477, 289–294. nature10413 [pii] 10.1038/nature10413.

145. Waskom, M. (2021). seaborn: statistical data visualization. Journal of Open Source Software 6. 10.21105/joss.03021.

146. Li, H., Handsaker, B., Wysoker, A., Fennell, T., Ruan, J., Homer, N., Marth, G., Abecasis, G., Durbin, R., and Genome Project Data Processing, S. (2009). The Sequence Alignment/Map format and SAMtools. Bioinformatics 25, 2078–2079. 10.1093/bioinformatics/btp352.

147. Robinson, J.T., Thorvaldsdottir, H., Winckler, W., Guttman, M., Lander, E.S., Getz, G., and Mesirov, J.P. (2011). Integrative genomics viewer. Nat Biotechnol 29, 24–26. 10.1038/nbt.1754.

148. Zhang, Y., Liu, T., Meyer, C.A., Eeckhoute, J., Johnson, D.S., Bernstein, B.E., Nusbaum, C., Myers, R.M., Brown, M., Li, W., and Liu, X.S. (2008). Model-based analysis of ChIP-Seq (MACS). Genome Biol 9, R137. 10.1186/gb-2008-9-9-r137.

149. Quinlan, A.R., and Hall, I.M. (2010). BEDTools: a flexible suite of utilities for comparing genomic features. Bioinformatics 26, 841–842. 10.1093/bioinformatics/btq033.

150. Bailey, T.L. (2021). STREME: accurate and versatile sequence motif discovery. Bioinformatics 37, 2834–2840. 10.1093/bioinformatics/btab203.

151. Castanza, A.S., Recla, J.M., Eby, D., Thorvaldsdottir, H., Bult, C.J., and Mesirov, J.P. (2023). Extending support for mouse data in the Molecular Signatures Database (MSigDB). Nat Methods 20, 1619–1620. 10.1038/s41592-023-02014-7.

152. Subramanian, A., Tamayo, P., Mootha, V.K., Mukherjee, S., Ebert, B.L., Gillette, M.A., Paulovich, A., Pomeroy, S.L., Golub, T.R., Lander, E.S., and Mesirov, J.P. (2005). Gene set enrichment analysis: a knowledge-based approach for interpreting genome-wide expression profiles. Proc Natl Acad Sci U S A 102, 15545–15550. 10.1073/pnas.0506580102.

153. Yamazaki, T., and Hirose, T. (2015). The building process of the functional paraspeckle with long non-coding RNAs. Front Biosci (Elite Ed) 7, 1–41. 10.2741/715.

154. Kong, A.T., Leprevost, F.V., Avtonomov, D.M., Mellacheruvu, D., and Nesvizhskii, A.I. (2017). MSFragger: ultrafast and comprehensive peptide identification in mass spectrometry-based proteomics. Nat Methods 14, 513–520. 10.1038/nmeth.4256.

155. Consortium, G.T. (2020). The GTEx Consortium atlas of genetic regulatory effects across human tissues. Science 369, 1318–1330. 10.1126/science.aaz1776.

156. Tang, Z., Li, C., Kang, B., Gao, G., Li, C., and Zhang, Z. (2017). GEPIA: a web server for cancer and normal gene expression profiling and interactive analyses. Nucleic Acids Res 45, W98–W102. 10.1093/nar/gkx247.

157. Goldman, M., Craft, B., Swatloski, T., Cline, M., Morozova, O., Diekhans, M., Haussler, D., and Zhu, J. (2015). The UCSC Cancer Genomics Browser: update 2015. Nucleic Acids Res 43, D812–817. 10.1093/nar/gku1073.

158. Mielke, P.W., and Berry, K.J. (2001). Permutation methods : a distance function approach (Springer).

159. Benjamini, Y., and Hochberg, Y. (1995). Controlling the false discovery rate: a practical and powerful approach to multiple hypothesis testing. Journal of the Royal Statistical Society 57, 289–300. https://www.jstor.org/stable/2346101.

160. MathWorks. MATLAB version: 9.13.0 (R2022b). https://www.mathworks.com.

161. Hulsen, T. (2022). DeepVenn -- a web application for the creation of area-proportional Venn diagrams using the deep learning framework Tensorflow.js. arXiv, arXiv:2210.04597. 10.48550/arXiv.2210.04597.

162. Kania, E.E., Fenix, A., Marciniak, D.M., Lin, Q., Bianchi, S., Hristov, B., Li, S., Camplisson, C.K., Fields, R., Beliveau, B.J., et al. (2024). Nascent transcript O-MAP reveals the molecular architecture of a single-locus subnuclear compartment built by RBM20 and the TTN RNA. bioRxiv. 10.1101/2024.11.05.622011.

163. Chang, Y., Wang, X., Yang, J., Tien, J.C., Mannan, R., Cruz, G., Zhang, Y., Vo, J.N., Magnuson, B., Mahapatra, S., et al. (2024). Development of an orally bioavailable CDK12/13 degrader and induction of synthetic lethality with AKT pathway inhibition. Cell Rep Med 5, 101752. 10.1016/j.xcrm.2024.101752.

164. da Veiga Leprevost, F., Haynes, S.E., Avtonomov, D.M., Chang, H.Y., Shanmugam, A.K., Mellacheruvu, D., Kong, A.T., and Nesvizhskii, A.I. (2020). Philosopher: a versatile toolkit for shotgun proteomics data analysis. Nat Methods 17, 869–870. 10.1038/s41592-020-0912-y.

